# Population coding under the scale invariance of high-dimensional noise

**DOI:** 10.1101/2024.08.23.608710

**Authors:** S. Amin Moosavi, Sai Sumedh R. Hindupur, Hideaki Shimazaki

## Abstract

High-dimensional scale-invariant neural activity is ubiquitous across brain regions and species, but its implications for information coding remain unclear. Here, we ask how stimulus information in the high-dimensional activity of mouse V1 scales with neuron number: does it saturate due to noise correlations, or increase without bound as subpopulations grow? Contrary to previous reports, we find that leading noise components that scale linearly with population size, and thus can limit information, are not sufficiently aligned with the signal to impose a bound. This conclusion follows from two scale-invariant power-law properties of neuronal responses in mouse V1: the noise eigenspectrum and alignment of noise components with the signal. We show that population subsampling links the observed power-law exponents to information boundedness, and that information scaling depends on the full eigenspectrum rather than its leading modes. Finally, we prove that, under subsampling, information-limiting correlations, if present, are differential correlations. Our findings clarify how information scales in high-dimensional neuronal activity under scale-invariant noise.

**Teaser:** Scale-invariant noise in mouse V1 does not limit stimulus information as population size increases.

## Introduction

Neural activity is highly variable yet structured in a high-dimensional space (*1–3*), ubiquitously exhibiting quasi-universal, scale-invariant power-law spectra of noise components across various brain regions and species (*4–6*). Recent advances in theoretical and empirical studies suggest the potential mechanistic and functional origins of these scale-invariant properties (*4–8*). However, their impact on population coding remains poorly understood.

Classic works on neuronal activity in the monkey middle temporal (MT) area have revealed that the activity of single neurons is highly informative about the animal’s perceptual sensitivity in discriminating stimulus directions (*9, 10*), suggesting that neurons encode stimuli in an overly redundant manner. In redundant coding, the information conveyed about a stimulus increases sublinearly as the size of the neural population grows, in contrast to the linear increase observed with independent coding (*10, 11*). A hallmark of redundant coding is shared noise in neural activity across repeated trials, which obscures the signal quantified by the change in the average population activity in response to a change in stimulus (*12–14*). Recent high-throughput techniques for the simultaneous recording of hundreds or thousands of neurons have made it possible to visualize the sublinear increase in the conveyed information as a function of the observed number of neurons (*15–18*).

Redundant coding can be so strong that it may limit information: Coded information may saturate at its maximum value in the limit of a large number of neurons (*10, 19, 20*). Here, we assessed information scaling in large-scale recordings of neurons in mouse primary visual cortex (V1) (*21*) and tested the boundedness in substantially larger networks beyond observed populations. To this end, we used two scale-invariant properties revealed by the data: the scale-invariant noise spectrum and the scale-invariant relationship between the magnitude of noise components and their alignment with the signal. The approach transcends earlier large-scale studies (*15–18, 21*) by replacing oversimplified assumptions about covariance scaling with the quasi-universal scaling observed in the data. Consequently, while these studies reported that information is bounded, we show that noise correlations, despite their detrimental effect on stimulus coding, do not limit the information. Therefore, neural populations have the capacity to encode stimulus information without a bound. These results demonstrate that the quasi-universal scaling of neural noise covariance provides a foundation for revealing the scaling and boundedness of population codes.

To establish the general principles underlying these findings, we developed a population coding theory based on the scaling properties of noise covariance under population subsampling. We derive a necessary and sufficient condition expressed in terms of covariance-eigenvalue scaling and the corresponding signal angular occupations, under which information remains bounded in the large-population limit. This condition highlights the critical need to consider the full spectrum of high-dimensional noise when predicting the scaling and boundedness of information coding in large neuronal populations. We further elucidate the constraints imposed by the covariance scaling on the power-law exponent of its eigenvalue spectrum, and clarify how this exponent relates to the boundedness of information. Finally, we characterize the incremental information gain by adding a neuron and relate the bounded regime to the specific noise structure known as differential correlations, which has been proposed to fundamentally limit information (*22*).

## Results

### Scaling laws for subsampled neuronal populations

The relationship between a signal and noise in the activity of *N* neurons evoked by multiple presentations of a stimulus (*s*) is represented in an *N*-dimensional space, where each dimension corresponds to the activity of a neuron (Fig. 1A). In this space, a vector **f**_*N*_ (*s*) = (*f*_1_(*s*), *f*_2_(*s*), …, *f*_*N*_ (*s*))^T^ represents trial-averaged ensemble activity, and the signal denoted by 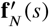 is the change in **f**_*N*_ (*s*) caused by an infinitesimal change in the stimulus. Noise is the deviation from **f**_*N*_ (*s*) during each stimulus presentation, quantified by a noise covariance matrix **Σ**_*N*_ that is represented by its eigenvectors **u**_*i,N*_ and eigenvalues *λ*_*i,N*_ where *i* = 1, …, *N*. These eigenvalues are indexed in descending order: *λ*_1,*N*_ ≥ *λ*_2,*N*_ ≥ ⋯ ≥ *λ*_*N,N*_. The overlap between 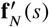 and each **u**_*i,N*_ is measured by the angle *θ*_*i,N*_ between the two vectors (Fig. 1B). Throughout this work, we focus on how these eigenvalues and angles scale as the population size *N* increases.

**Fig. 1.**
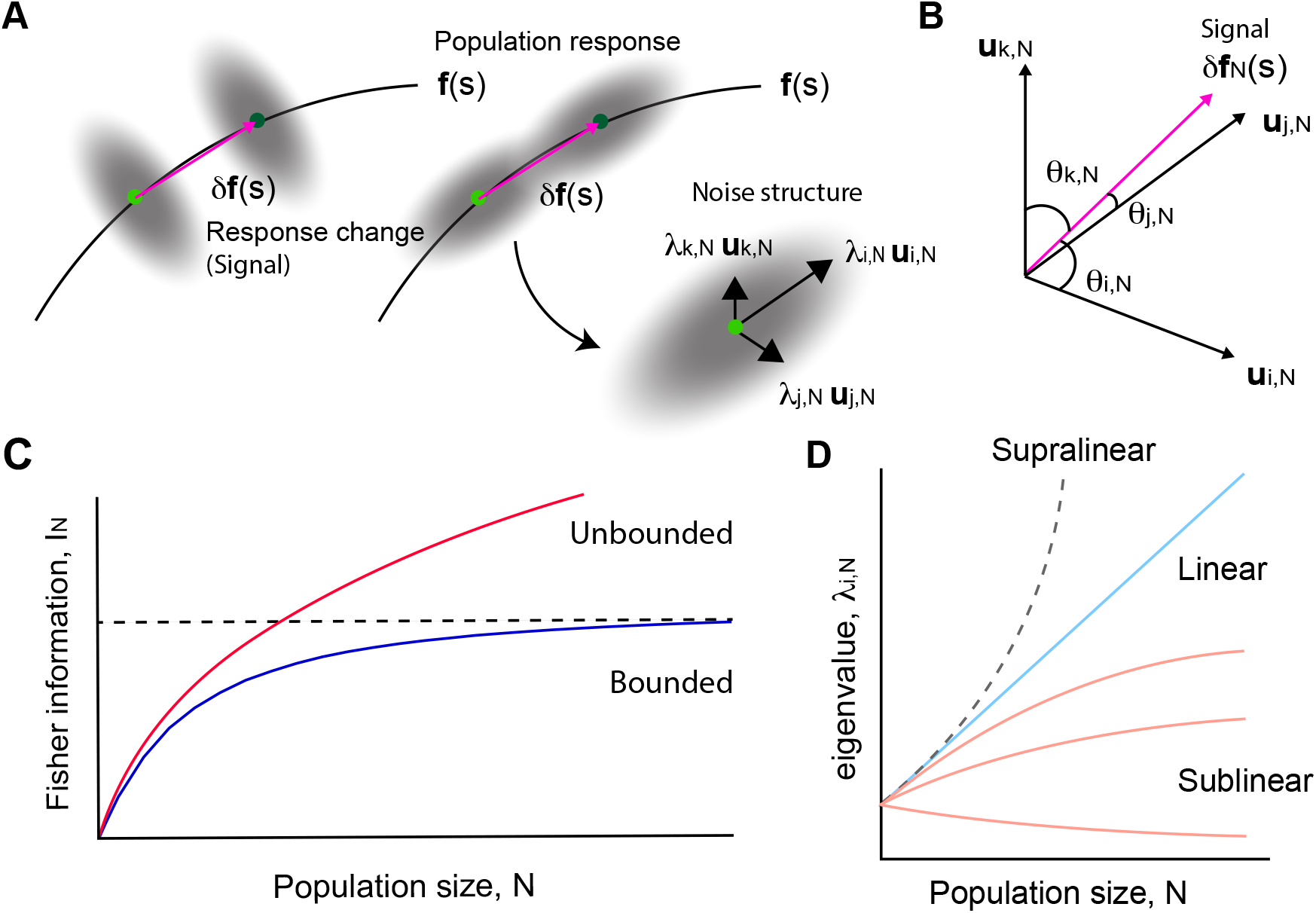
Schematic illustrating necessary and sufficient conditions for bounded and unbounded LFI. **A**, A schematic of a subspace spanned by three neurons. The mean activity of neurons 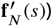 encodes the stimulus *s* as first-order statistics (solid black line). In addition to the change in the mean responses (green dots) to the two close stimuli (*δ***f**_*N*_ (*s*), magenta arrow, a finite-size analog of **f** ^′^ (*s*)), the covariability (gray areas) affects stimulus coding beneficially (left) or detrimentally (right) when compared to circular independent noise. Noise covariance is quantified by eigenvectors **u**_*i,N*_, **u**_*j,N*_, **u**_*k,N*_ and their respective noise strengths *λ*_*i,N*_, *λ*_*j,N*_, *λ*_*k,N*_. **B**, A three-dimensional view of the orientation of *δ***f**_*N*_ (*s*) with respect to the covariance eigenvectors. *θ*_*i,N*_ is the angle between *δ***f**_*N*_ (*s*) and the eigenvector **u**_*i,N*_. **C**, The scaling of bounded (blue) and unbounded (red) linear Fisher information. **D**, Supralinear, linear, and sublinear eigenvalues as a function of *N*. Supralinear eigenvalues are prohibited for subsampled populations.

The coding capacity is quantitatively measured by the linear Fisher information (LFI) defined as 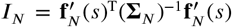 (*13, 19, 20, 22, 23*). We note that, although *I*_*N*_ depends on the stimulus, in this study we focus on its scaling with population size rather than variations with stimulus. Therefore, we constrained our analysis to small stimulus variations around a fixed value *s*. Using the noise components’ eigenvalues and the corresponding angles with 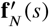, the LFI can equivalently be expressed as (text S1):

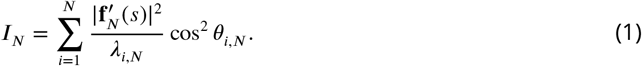

The LFI gives an upper bound on the information that linear decoders can read out. In other words, the optimal linear decoder—obtained by multiplying the inverse of the covariance matrix 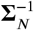 with the signal vector 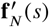—achieves a discrimination precision equal to the LFI (*13*).

The alignment of each noise component with the signal 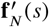 is measured by cos^2^ *θ*_*i,N*_ termed angular occupation, which, due to the completeness of the orthonormal basis, satisfies

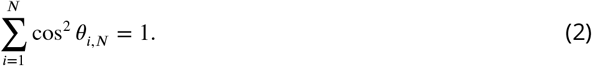

Noise components that are not orthogonal to the signal 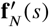 (cos^2^ *θ*_*i,N*_ ≠ 0) contribute to the LFI. In correlated population responses, the alignment of 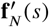 with larger components impairs the coding accuracy, and alignment with smaller ones improves the coding accuracy, compared to independent neurons.

With the above representation, we examined the scaling of the LFI with increasing subpopulation size *N* (Fig. 1C). Each subpopulation is constructed by randomly subsampling *N* neurons from a total of *N*_*t*_ recorded neurons (*N* ≤ *N*_*t*_). Three factors contribute to the scaling of LFI: the signal magnitude, the noise components’ strength, and their alignments. The signal magnitude 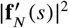 scales linearly with *N* under the random subsampling,

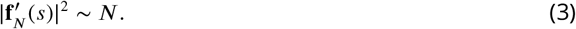

This is because 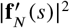 is a summation of *N* nonnegative variables, which increases linearly with *N* if these variables have a finite, nonzero mean (text S1). Thus, whether the LFI is limited or not depends on how *λ*_*i,N*_ scales against the linear 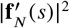 and how the angular occupation cos^2^ *θ*_*i,N*_ changes with *N*. Therefore, to assess the LFI scaling, from here onward, we categorized the covariance eigenvalues based on their behavior versus *N* as supralinear, linear, and sublinear eigenvalues (Fig. 1D, see text S2 for definition). Linear eigenvalues, if they exist, appear at early indices in the eigenspectrum in descending order, followed by the sublinear eigenvalues. These linear eigenvalues can potentially limit information growth if their associated noise directions sufficiently overlap with the signal direction.

The eigenvalues of the covariance matrix of subsampled populations are fundamentally constrained: the total variance must grow linearly with the population size *N*,

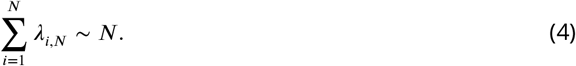

This follows from the fact that the sum of the covariance eigenvalues equals the total variance, which is itself the sum of the individual variances of all *N* neurons. Because each neuron’s variance is independent of the subsample size *N*, the total variance, like the signal magnitude, scales proportionally with *N*.

This introduces several crucial constraints on the eigenspectrum of the covariance matrix (text S2). First, the subsampled systems cannot have supralinear eigenvalues. Second, consider eigenvalue sequences that, for a fixed rank *i*, scale linearly with *N*. If their slopes remain above a fixed positive value, then their number must converge to a finite value; otherwise, their sum grows faster than *N*. The remaining eigenvalues, whose number then grows linearly with *N*, are sublinear, and thus dominate the spectrum by number in the large-*N* limit. Third, if the slopes of the linear eigen-values are allowed to shrink with the eigenvalue ranking such that their sum over rank is finite (i.e., summable), the spectrum may contain a countably infinite family of fixed-rank linear eigen-value sequences. In this case, linear total-variance scaling alone does not constrain the number of sublinear eigenvalues.

Having established the spectral scaling constraints that govern information boundedness under population subsampling, we then turned to experimental data to assess how the noise co-variance spectrum scales in mouse V1 and how its alignment with 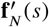 influences information boundedness.

### Unbounded information scaling in mouse V1 neurons

We analyzed stimulus-evoked responses of a population of mouse V1 neurons recorded by two-photon imaging (see Materials and Methods). The experiments were designed to examine the population’s ability to detect small changes in orientations of grating stimuli (*21*). In each trial, the stimulus orientation was randomly drawn from a uniform distribution in the range [43°, 47°]. Following Stringer et al. (*21*), we grouped the trials into two adjacent angular ranges to quantify local discriminability between the two groups (Fig. 2A). Although the presented stimuli are finite and discrete, the question here is how local discriminability scales with subpopulation size. The dataset includes 17,631–21,469 neurons recorded from five mice with ~1000 repeated trials per degree while animals passively viewed stimuli.

**Fig. 2.**
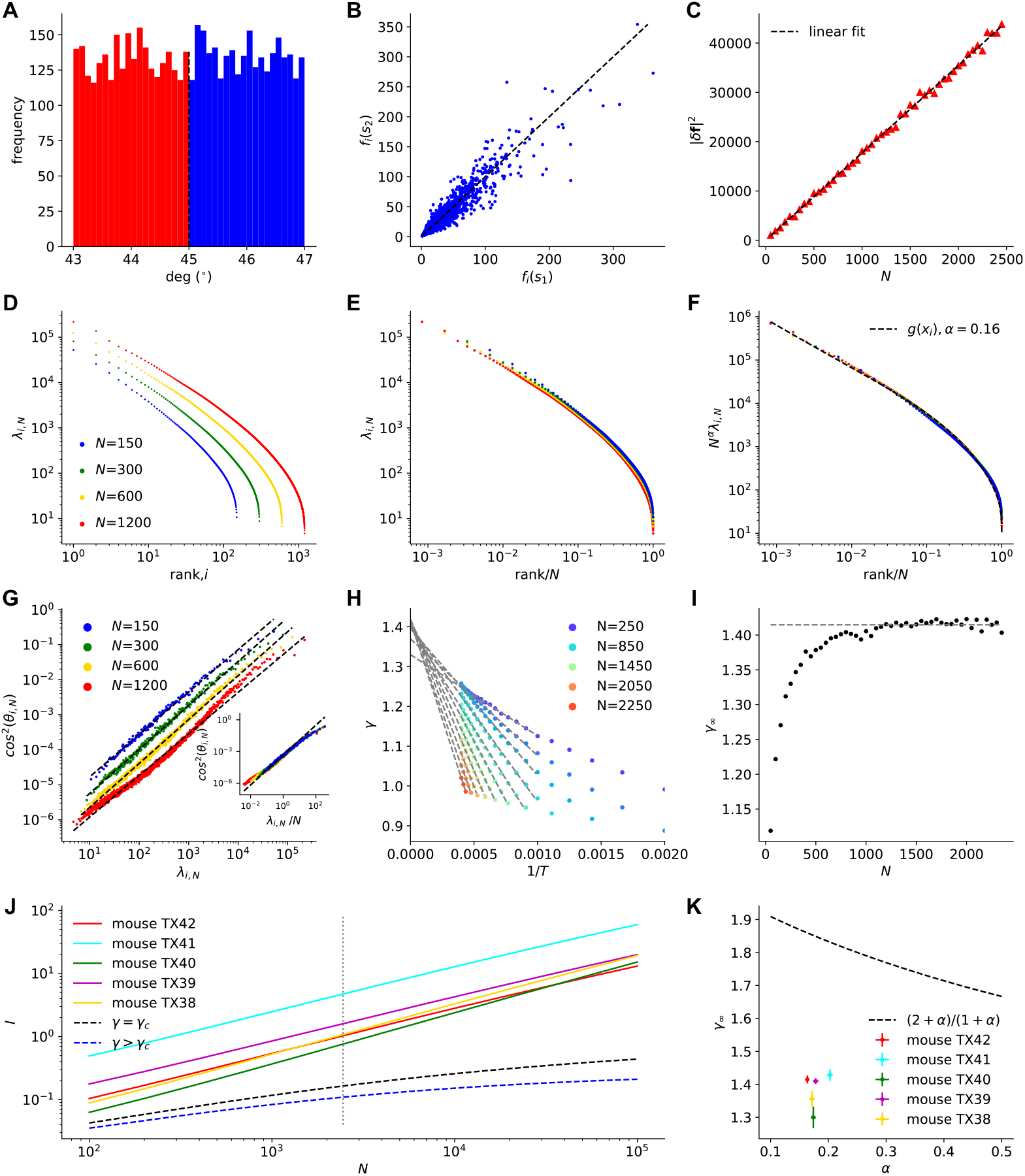
Scaling analysis of mouse V1 neurons. **A**, Histogram of grating stimulus orientations presented to mouse TX42. Stimuli with orientations from 43° to 45°, including 45° (red), are categorized as Stimulus 1 and those above 45° and up to 47° (blue) as Stimulus 2. **B**, Mean responses *f*_*i*_(*s*) of neurons (*i* = 1, …, 18,982) to Stimuli 1 (*s* = *s*_1_) vs 2 (*s* = *s*_2_) for TX42. *f*_*i*_(*s*) is the mean response over trials in each stimulus category. Dashed line, identity. **C**, Mean response changes |*δ***f**_*N*_|^2^ vs subpopulation size *N*, where *δ***f**_*N*_ = **f**_*N*_ (*s*_1_) − **f**_*N*_ (*s*_2_), for TX42. Values are averaged over 200 randomly chosen subpopulations. Dashed line, linear fit. **D**, Noise eigenspectra *λ*_*i,N*_ for different subpopulation sizes *N* in TX42. **E**, Eigenspectra vs the normalized rank, *x*_*i*_ = *i*/*N*. **F**, Rescaled eigenspectra *N*^*α*^*λ*_*i,N*_, showing collapse onto the fitted scaling function *g*(*x*_*i*_) (dashed line, Eq. 6). For TX42, *α* = 0.164 ± 0.002; fitted parameters of *g*(*x*_*i*_) are *a* = 2.752, exp(*b*) = 9.611, *c* = 2.298, and *ν* = 0.254. **G**, cos^2^ *θ*_*i,N*_ vs *λ*_*i,N*_ for different *N* in TX42 using all trials (2500 per stimulus). Dashed lines, linear fits. Inset: cos^2^ *θ*_*i,N*_ vs *λ*_*i,N*_ /*N*. **H**, Fitted exponent *γ* in Eq. 7 vs 1/*T* for different *N*, where *T* is the number of trials. Solid lines, linear fits; intercepts estimate *γ* at an infinite trial number. **I**, Intercept *γ*_∞_ vs *N*. Mean *γ*_∞_ for *N* = 1000–2500 (dashed line) was 1.415 ± 0.006. **J**, Inferred LFI vs *N* for TX42 and four other mice (solid lines). The LFI was computed from Eqs. 5 and 7, using the parameters fitted to each mouse. Dashed lines show theoretical LFI resulting in critical (black, *γ* = *γ*_*c*_) and bounded information (blue, *γ* > *γ*_*c*_) for TX42. Vertical dashed line, neuron number used for parameter estimation. **K**, Estimated *γ*_∞_ vs *α* for all mice. Dashed line, critical boundary (Eq. 8); error bars, 2SD.

These neurons exhibited linearly scaling 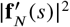 (Fig. 2B,C), as expected. Figure 2D shows that the eigenspectrum of the noise covariance matrix scales systematically with subpopulation size. When the eigenspectrum is represented using the normalized index *x*_*i*_ = *i*/*N* (Fig. 2E), the eigen-spectrum is in agreement with the following scale invariance,

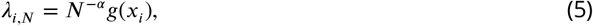

where *α* > 0. Accordingly, plotting *N*^*α*^*λ*_*i,N*_ against *x*_*i*_ produces a collapse of data onto a scale-invariant function *g*(*x*_*i*_) that is independent of subpopulation size (Fig. 2F). For mouse TX42, the exponent obtained by finite-size scaling collapse of the data (see Materials and Methods) was *α* = 0.164. This exponent appears quasi-universal among all the mice in the experiment (*α* = 0.18 ± 0.01, mean ± standard deviation (SD) across five mice; figs. S1–S5).

By carefully constructing an ansatz consistent with the subsampling constraints and the observed scaling collapse, we found the following form for the scale-invariant function:

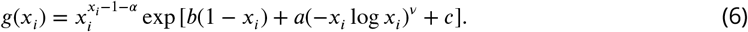

The exponential function accounts for the cutoff that occurs at *x*_*i*_ = 1 by definition, whereas the power-law part explains the behavior of larger eigenvalues in the *x*_*i*_ → 0 limit. In particular, considering Eq. 5, the power law is defined such that linear eigenvalues are permitted as *x*_*i*_ → 0 when *α* > 0; and the total variance, i.e., the sum of all eigenvalues, scales linearly with *N*. While the functional form of *g*(*x*_*i*_) is independent of *N*, its domain *x*_*i*_ ∈ [1/*N*, 2/*N*, …, 1] depends on *N*. Unlike *α*, the remaining fitted parameters of *g*(*x*_*i*_) varied across mice.

The equations above specify the composition of the eigenspectrum in the large-*N* limit. At small normalized rank (*x*_*i*_ ≪ 1), the scaling function follows a power law, 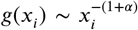. This power law describes scaling with eigenvalue rank, whereas Eq. 5 describes scaling with subpopulation size *N*. When *α* > 0, the eigenvalue at any fixed rank *i* grows linearly with *N* because *λ*_*i,N*_ ~ *i*^−(1+*α*)^*N*. Namely, for every fixed positive integer *N*_*c*_, the first *N*_*c*_ eigenvalue sequences increase linearly with *N*. Hence, in the large-*N* limit, the spectrum contains a countably infinite family of fixed-rank linear eigenvalue sequences. The contribution of this fixed-rank linear family is compatible with the linear-total-variance constraint (Eq. 4) when the linear coefficients *i*^−(1+*α*)^ are summable, i.e., *α* > 0. Although this family is countably infinite, eigenvalues at ranks proportional to *N* scale as *N*^−*α*^ and are therefore sublinear. Thus, sublinear eigenvalues dominate the spectrum by number in the large-*N* limit.

The alignment of the noise eigenvectors linked to linear or sublinear eigenvalues with 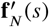 determines whether the LFI is bounded. The angular occupations cos^2^ *θ*_*i,N*_ and eigenvalues *λ*_*i,N*_ show linear dependency on a logarithmic scale (Fig. 2G), indicating another scale-invariant relation:

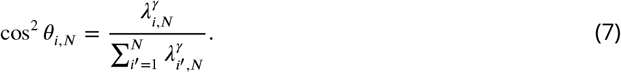

The eigenvectors tied to the larger eigenvalues occupy a large portion of the angular occupations, indicating that the linear eigenvalues align with 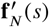 and potentially limit the information. By contrast, the eigenvectors associated with the smaller eigenvalues exhibit smaller angular occupations (nearly orthogonal to 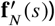. Yet, due to the sublinearity, their contributions to the coded information can be substantial. From Eqs. 1, 4, and 7, one can see *I*_*N*_ ~ *N* for *γ* = 1 and *I*_*N*_ → *I*_∞_ for *γ* = 2, where *I*_∞_ = lim_*N*→∞_ *I*_*N*_ < ∞ denotes the limiting LFI (Materials and Methods; text S3). Therefore, a critical exponent *γ* = *γ*_*c*_ exists between one and two, where the information changes behavior from unbounded to bounded. We find that this critical exponent is given by

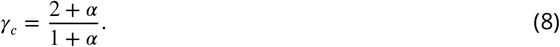

Specifically, in the large-*N* limit, the LFI scales as

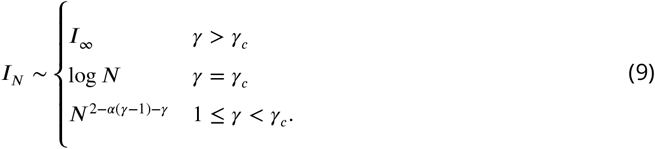

For *α* = 0.164 in mouse TX42, the critical exponent is *γ*_*c*_ = 1.859. The exponent *γ* estimated by extrapolating to the infinite-trial limit, denoted *γ*_∞_, is *γ*_∞_ = 1.415 ± 0.006 (Fig. 2H,I). As *γ*_∞_ is smaller than *γ*_*c*_, the result indicates that the LFI of the neurons in this dataset increases without bound. In this regime, the LFI asymptotically scales as a power law with the exponent 2 −*α*(*γ* − 1) −*γ* = 0.517, which is a sublinear but unbounded function of *N*. Such sublinear, unbounded behavior was consistently observed across all five mice (Fig. 2J,K; see figs. S1–S5 for individual results). Accordingly, we conclude that the mouse V1 neurons’ correlated activity causes redundancy, yet it is insufficient to limit information in the large-*N* limit. Therefore, the only constraints on the information capacity are the limited stimulus information available in the retina (*24*) and the total number of neurons in V1 and related areas, which are not imposed in this theory.

We performed the analyses above using the raw sample eigenvalues and eigenvectors. However, because finite-sample estimation introduces biases into both the eigenspectrum and the angular occupations, we also developed a finite-sample bias-correction method. The procedure consists of three steps: iterative shrinkage of the eigenspectrum, an iterative Wishart-surrogate correction for angular occupations, and an analytical perturbative correction for residual bias in the estimated signal vector. Together, these steps yielded reliable bias-corrected estimates of both the eigenspectrum and the angular occupations (fig. S6). See the Materials and Methods section for details of the algorithms. We applied the method to empirical data (fig. S7) and verified that the main conclusions were robust to these corrections: after bias correction, the exponents were *α* = 0.10 ± 0.02, *γ* = 1.26 ± 0.20, and *γ*_*c*_ = 1.91 ± 0.01 (mean ± SD across five mice), with *γ* < *γ*_*c*_ in every animal.

### Linear total variance of subsampled populations constrains eigenvalue scaling, power-law exponent, and boundedness of the LFI

Here we summarize the general consequences for scale-invariant eigenspectra (Eq. 5) imposed by the linear scaling of the total variance required for subsampled populations (Eq. 4). Since there are no supralinear eigenvalues for subsampled populations, the LFI (Eq. 1) can be decomposed into the contributions by components with linear and sublinear eigenvalues:

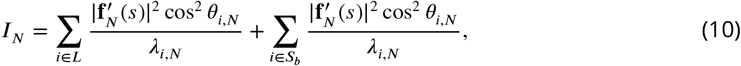

where *L* and *S*_*b*_ are sets of linear and sublinear eigenvalues, and 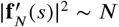.

Because the two contributions are nonnegative, *I*_*N*_ is bounded if and only if both are bounded. We first consider the case in which the fixed-rank linear family is finite. At large *N*, the number of elements in set *L* approaches a constant integer *c* ≥ 0; consequently, the number of elements in set *S*_*b*_ linearly increases with *N*: |*L*| → *c* and |*S*_*b*_| → ∞ as *N* → ∞. The first term remains bounded because *L* is finite and, for each *i* ∈ *L, N*/*λ*_*i,N*_ → 1/*b*_*i*_ < ∞. Therefore, in this first scenario, the boundedness of the second summation is necessary and sufficient for bounded information: LFI boundedness is determined by the sublinear components, not by the linear eigenvalues. Defining the occupation of cos^2^ *θ*_*i,N*_ from sublinear eigenvectors by 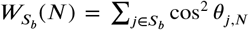, the boundedness condition can be written as (text S4)

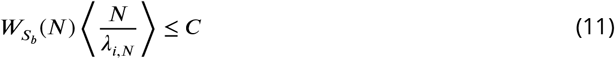

for all sufficiently large *N*, where *C* > 0 is a finite constant independent of *N*. For 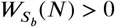, we define 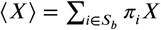, where 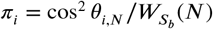 if 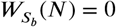, the sublinear contribution vanishes. Equivalently, the necessary and sufficient condition for bounded LFI is that 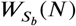 decreases to zero at a rate equal to or faster than ⟨*N*/*λ*_*i,N*_⟩^−1^, which sets the critical rate of convergence. This critical rate is determined by the scaling of the sublinear eigenvalues together with their associated angular occupations. This condition can equivalently be expressed in terms of the angular occupation of the linear eigenvectors, *W*_*L*_(*N*), using 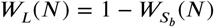. Specifically, 1 − *W*_*L*_(*N*) must decrease to zero at least as fast as the critical rate set by the sublinear components. This shows that alignment of the signal with the linear-eigenvalue subspace (*W*_*L*_(*N*) → 1) is necessary but not sufficient for bounded LFI.

Next, we consider the case in which the family of fixed-rank linear eigenvalue sequences is countably infinite and their linear slopes (*b*_*i*_), defined by *λ*_*i,N*_ ~ *b*_*i*_*N*, form a summable sequence, 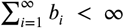. In this case, in addition to the boundedness of the sublinear-eigenvalue contribution, the angular occupations of the fixed-rank linear eigenvalue sequences must decay sufficiently rapidly with rank for their cumulative contribution to remain bounded. This scenario is consistent with the experimental results presented above, where the eigenvalue slopes decay according to a power law, *b*_*i*_ ~ *i*^−(1+*α*)^ with *α* > 0. Indeed, for 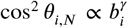 from Eq. 7, the linear-eigenvalue contribution reduces to 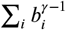, and its summability, i.e., convergence of the summation 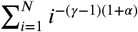 yields the critical exponent *γ*_*c*_ given in Eq. 8. The full-spectrum calculation in text S3 further shows that the same condition ensures boundedness of the total LFI, including the cumulative contribution of the sublinear eigenvalues.

Furthermore, combining power-law eigenspectral scaling with the linear scaling of the total variance imposes important constraints on the power-law exponent and, consequently, on the boundedness of the LFI (Materials and Methods; full derivation in texts S5 and S6). Because the scale-invariant function is defined on *x*_*i*_ = *i*/*N* ∈ [1/*N*, 1], the *N*-dependence enters through the left edge of the eigenspectrum, corresponding to the leading eigenvalues. Thus, the *x*_*i*_ → 0 behavior determines the scaling of the total variance, even though the variance is defined as the sum over the full eigenspectrum. The results are summarized in Table 1. Given the power-law form 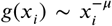, we showed that the constraint of the linear total variance (Eq. 4) leads to *μ* = 1 + *α* if *α* is positive. In this regime, the leading eigenvalues scale linearly with population size, *λ*_*i,N*_ ~ *N* (for fixed *i*). The selected functional form for *g*(*x*_*i*_) in Eq. 6, specifically 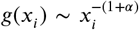 in the *x*_*i*_ → 0 limit, is consistent with the linear scaling of the total variance (Eq. 4) and permits linear eigenvalues with *α* > 0. Such linear scaling can potentially bound the LFI, provided that the corresponding eigenvectors align sufficiently with the signal direction.

**Table 1.** Scaling parameter, power-law exponent, and information boundedness under subsampling. For a scale-invariant power-law eigenspectrum, the rows summarize how the scaling parameter *α* constrains the power-law exponent *μ*, the scaling of the leading eigenvalues *λ*_*i,N*_, and whether the LFI is bounded or unbounded under the linear scaling of the total variance.

| Scaling parameter $\alpha$ | Power-law exponent $\mu$ | Scaling of leading eigenvalues | Linear Fisher information |
| --- | --- | --- | --- |
| $\lambda_{i,N} = N^{-\alpha} g(x_i) \quad (x_i = i/N)$ | $g(x_i) \sim x_i^{-\mu} \quad (x_i \ll 1)$ | $\lambda_{i,N} \sim N^{\mu-\alpha} \quad (\text{fixed } i)$ | $I_N$ |
| $\alpha > 0$ | $\mu = 1 + \alpha > 1$ | $\lambda_{i,N} \sim N \quad (\text{linear})$ | bounded/unbounded |
| $\alpha = 0$ | $\mu < 1$ | $\lambda_{i,N} \sim N^\mu \quad (\text{sublinear})$ | unbounded |

By contrast, when *α* = 0, the same constraint imposes *μ* < 1, indicating that the eigenspectrum must have a slope shallower than 1 if the eigenspectrum in the normalized index collapses in the limit of large *N*. In this case, the leading eigenvalues scale sublinearly, *λ*_*i,N*_ ~ *N*^*μ*^ (for fixed *i*). This means that all eigenvalues scale sublinearly, making the information an unbounded function of *N*, regardless of the behavior of angular occupations (text S1). Notably, this yields a sufficient condition requiring only the noise eigenspectrum: given 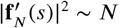, observing *α* = 0 (equivalently *μ* < 1) establishes unboundedness without estimating the signal–noise alignment. Indeed, when neural activities examined in the previous section are individually normalized by their nonzero mean activities (*6*) or standard deviations (*4, 5*), we observed that the slopes of leading eigenvalues satisfy *μ* < 1 (fig. S8), further supporting that the LFI is not bounded in this dataset.

We note that, if a scale-invariant relationship between the eigenvalues and their angular occupations holds (Eq. 7), one obtains an analytical expression for the LFI (Eq. 9), from which its boundedness can be determined as discussed in the previous section.

We validated these predictions by simulating population activity of neurons using the scale-invariant eigenspectrum defined by Eqs. 5 and 6 for *α* > 0 and a modified function for *α* = 0 (Fig. 3A top and bottom, respectively). For the *α* = 0 simulations, we replaced the leading-edge exponent 1 +*α* in Eq. 6 with an independently chosen *μ* = 0.5, while retaining the remaining parameters fitted to TX42. The simulation procedure is described in Materials and Methods. For the positive-*α* case, we used *α* = 0.16, close to the estimate from the mouse data. We confirmed the emergence of linear eigenvalues with *α* > 0, and their absence when *α* = 0 (Fig. 3B). For *α* > 0, we considered the signal and noise component directions given by Eq. 7 with *γ* = 2 and *γ* = 0 (Fig. 3C, top), which theoretically result in bounded and unbounded LFI, respectively. We also included an unbounded LFI example with *γ* = 1.415, which is close to the estimate from the mouse data. For *α* = 0, we expect unbounded LFI regardless of *γ*. Therefore, we consider an example with *γ* = 2 (Fig. 3C, bottom), which would have resulted in the bounded LFI if *α* were positive.

**Fig. 3.**
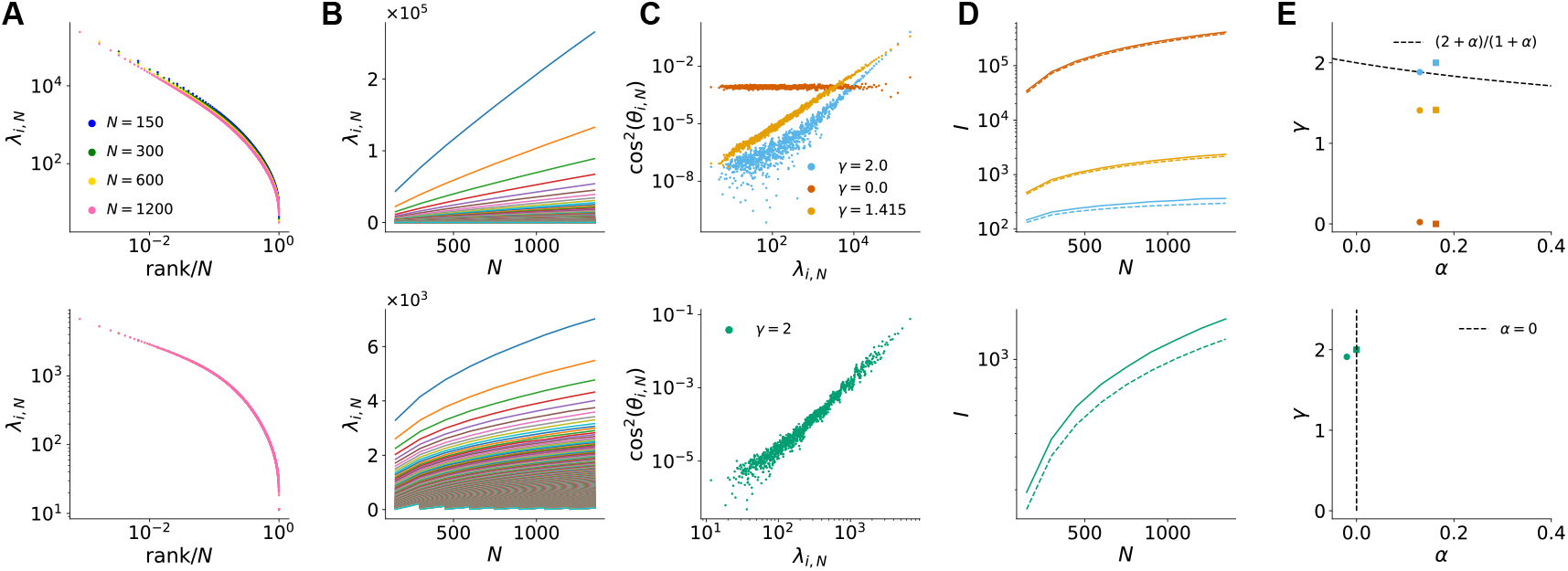
Verification of theory by simulated neural activity. **A**, Eigenspectra (generated as described in the text for *α* = 0.16 and *α* = 0) as a function of normalized rank for simulated populations with the scaling parameter *α* = 0.16 (observed value for mouse TX42, top) and *α* = 0 (bottom) for *N* = 150, 300, 600, 1200 neurons. **B**, Eigenvalues as a function of the subpopulation size *N* with *α* = 0.16 (top) and *α* = 0 (bottom), showing the emergence of linear eigenvalues only when *α* > 0. **C**, Relation between the angular occupation cos^2^ *θ*_*i,N*_ and eigenvalue *λ*_*i,N*_. Top, simulations with *α* = 0.16 using Eq. 7 with *γ* = 2 (a value resulting in bounded LFI for any *α* > 0, blue), *γ* = 0 (random signal, red), and *γ* = 1.415 (observed value for mouse TX42, yellow). Bottom, simulation with *α* = 0 and *γ* = 2. Plots are shown for *N* = 1200. **D**, Theoretical (dashed) and bias-corrected (solid) estimates of the LFI (*17, 25*) for the simulated populations. For *α* = 0.16, the bias-corrected estimates show weaker growth for *γ* = 2 than for *γ* = 0 and *γ* = 1.415, consistent with bounded and unbounded behavior, respectively. For *α* = 0, the bias-corrected estimate remains unbounded. **E**, True (squares) and estimated (circles) values of *γ* as a function of *α*. The dashed lines indicate the critical boundaries. The bias-correction methods were applied to this analysis.

We found that, as expected, the bias-corrected estimates of the LFI (*17, 25*) show weaker growth for *γ* = 2 than for *γ* = 0 when *α* > 0, consistent with bounded and unbounded behavior, respectively (Fig. 3D, top). For *α* = 0, the bias-corrected estimate of the LFI is consistent with unbounded behavior (Fig. 3D, bottom). However, because the bias-corrected LFI is evaluated only over the observed range of population sizes, it cannot by itself reveal the asymptotic scaling behavior at larger *N*.

Alternatively, our approach uses the fitted scaling ansatz to predict the behavior of larger neuronal populations. We corroborated that the estimated scaling exponents *α* and *γ* are consistent with the theoretical criterion for LFI boundedness, with the inferred parameter pairs tracking the boundary between bounded and unbounded regimes (Fig. 3E). For *α* > 0, we could correctly infer whether the LFI was bounded or unbounded from the estimated exponents, including for the experimentally observed value *γ* = 1.415. When *γ* is large, however, it induces extremely small cos^2^ *θ*_*i,N*_ (Fig. 3C, top), making them difficult to estimate reliably from a limited number of samples. As a result, the inferred parameter pairs can be biased toward the theoretical boundary; therefore, the results in this regime should be interpreted with caution and may require more data. For *α* = 0, by contrast, all eigenvalues remain sublinear under subsampling, so the LFI is theoretically unbounded regardless of *γ* (text S5). In this case, the estimated *α* was slightly negative (Fig. 3E, bottom), which can arise from finite-sample effects when the true *α* is zero and therefore indicates that the LFI is unbounded (text S5).

Finally, when we applied the dimension-reduction method to estimate the LFI (*16*) to simulated data whose LFI is theoretically unbounded, we observed saturating LFI (text S7, figs. S9 and S10). These results demonstrate that apparent information saturation can arise as an artifact of the analysis method focusing on leading eigenvalues, rather than as a property of the underlying population code, and highlight the importance of examining the full eigenvalue spectrum.

### Information-limiting correlations in subsampled populations are differential correlations

While we provided the scaling conditions for bounded and unbounded information under the linear scaling of the total variance, the connection between our theory and previously proposed information-limiting correlations needs clarification. Moreno-Bote et al. (*22*) introduced differential correlations as shared noise aligned with stimulus-dependent changes in the mean population response. Because this noise lies along the signal direction, it cannot be averaged away by adding neurons and can cause the LFI to saturate. More specifically, they showed that when the noise covariance has the following additive rank-one differential-correlation structure, the LFI is bounded:

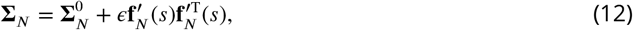

where *ϵ* is a positive constant and 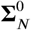 is a covariance matrix whose corresponding LFI

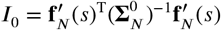

diverges with *N*. The resulting LFI is bounded above by *ϵ*^−1^ and is given by

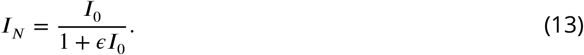

They further proposed that only differential correlations can limit the information. However, the original proof was restricted to specific noise covariances, leaving the claim unsubstantiated for general covariance matrices. Here, we prove that the information-limiting condition in our theory and the existence of differential correlations are not in general equivalent, but become identical under the subsampling constraint, where we can observe subsamples of varying size *N* of a fixed population, i.e., the total population is fixed and does not grow. To this end, we first quantitatively explain how the addition of a subsampled neuron to a population changes or does not change the LFI.

Recursively adding a neuron to a population creates a sequence of subsampled populations whose covariance matrices form a nested structure, with each larger covariance matrix containing that of the smaller subsampled population. Given *N* neurons, consider adding a single neuron whose mean response is *f*_*N*+1_(*s*), variance Σ_*N*+1_, and covariance **c**_*N*_ with the existing *N* neurons.

The covariance matrix of the resulting population of *N* + 1 neurons is

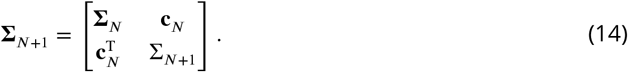

Exploiting this nested structure, the LFI of *N* + 1 neurons can be expressed recursively in terms of the LFI of *N* neurons as (text S8)

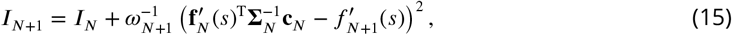

where 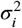. For a single neuron whose response follows a normal distribution with a mean *f*_1_(*s*) and variance 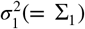, the LFI is given by 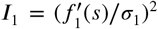, which is the squared standardized mean difference of two normal distributions for the finite difference case (Fig. 4A). Since *ω*_*N*+1_ > 0 is required for the covariance of *N* + 1 neurons to be positive definite, Eq. 15 indicates that the LFI along this nested sequence of subpopulations is a non-decreasing function of *N*. Further, Eq. 15 reveals two distinct ways to increase the LFI (Fig. 4A,B). First, when adding an independent neuron (i.e., **c**_*N*_ = **0**), the LFI increases by 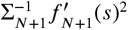, which is the LFI of the added neuron (Fig. 4B left). Second, the LFI increases even if 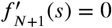 namely, even if we add a neuron whose activity is insensitive to the stimulus change, it will contribute to increasing information as long as 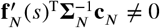. This is because adding a correlated neuron can remove the overlap of the noise distributions for the two close stimuli (Fig. 4B right), explaining an effect found in the sensory and frontal areas of rodents and primates (*26–30*).

**Fig. 4.**
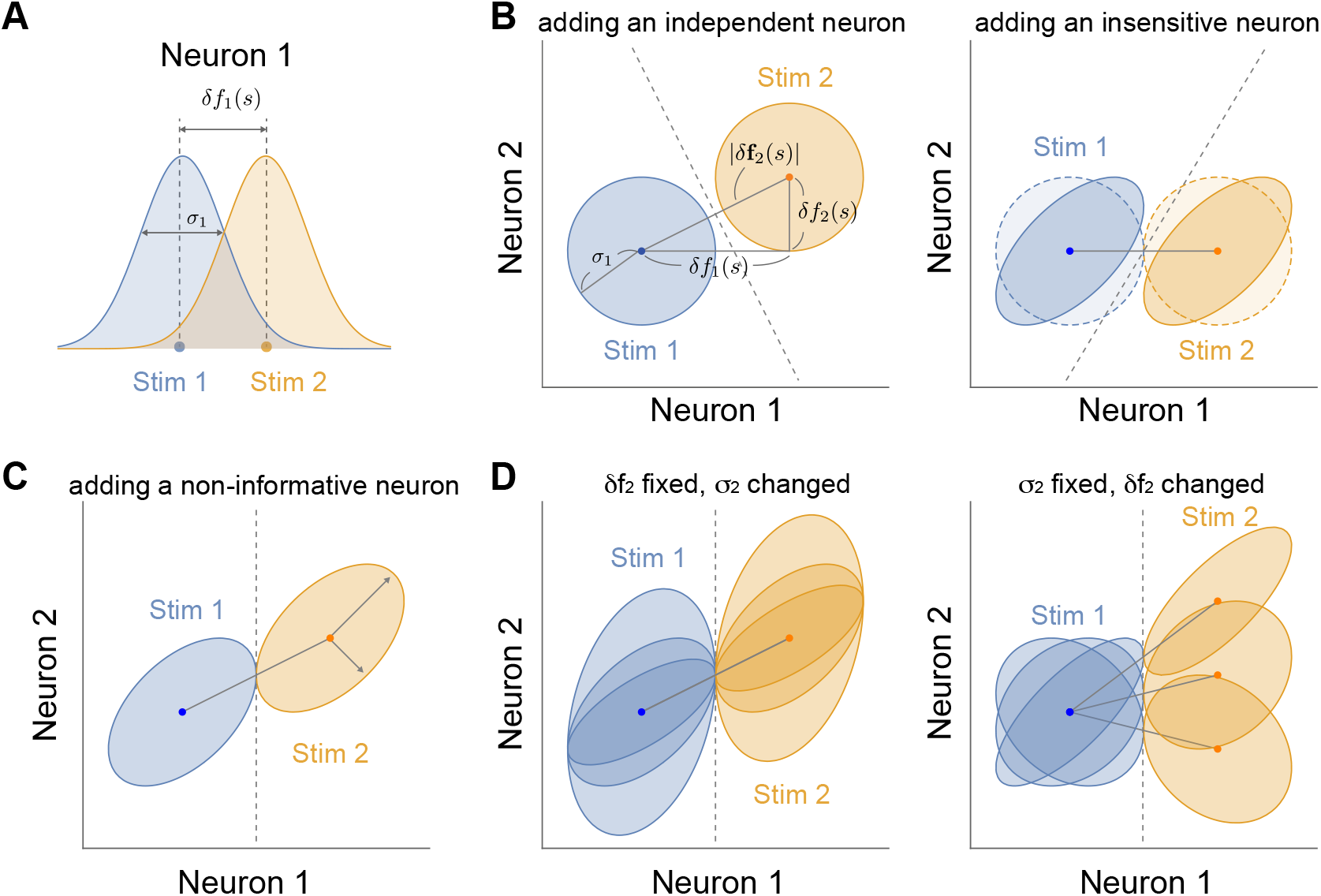
Incremental effect of adding a neuron on stimulus coding. **A**, Response distributions of a single neuron (Neuron 1) for two close stimuli (blue for Stimulus 1 and orange for Stimulus 2). The standardized mean difference *δf*_1_(*s*)/*σ*_1_ quantifies the ability to discriminate between the two stimuli. Here, *δf*_1_(*s*) = *σ*_1_. **B**, (Left) Joint distributions of two neurons after adding an independent, stimulus-sensitive neuron with the same variance (*σ*_2_ = *σ*_1_). The dashed line indicates the optimal linear decision boundary. The circle radius indicates one standard deviation. The distance between the mean responses for the two stimuli is 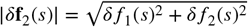. The horizontal separation between the two means remains *δf*_1_(*s*) = *σ*_1_, identical to panel A. (Right) Joint distributions after adding a correlated, stimulus-insensitive neuron (Neuron 2), having the same variance (*σ*_2_ = *σ*_1_). Neuron 2 does not change the mean activity, but correlation makes the distributions ellipsoidal. The dashed line indicates the optimal decision boundary for the correlated neurons. Light-colored circles show a case with adding an independent, stimulus-insensitive neuron for comparison. **C**, Joint distributions after adding a non-informative neuron (Neuron 2). Adding a correlated, stimulus-sensitive neuron does not increase discriminability from the single neuron’s performance when 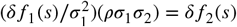 with *ρ* being the correlation coefficient. We impose 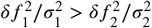 so that Neuron 2 has smaller LFI than Neuron 1. **D**, Variations of the joint distributions that preserve discriminability. (Left) *δf*_2_(*s*) is fixed, while *σ*_2_ and correlation vary. (Right) *σ*_2_ is fixed, while *δf*_2_(*s*) and correlation vary.

Moreover, the LFI does not increase if 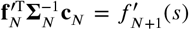. Figure 4C presents an illustration of the joint responses of two neurons for a finite stimulus difference that satisfies this condition, with Fig. 4D showing additional variations. In all cases, when the LFI remains unchanged, the decision boundary (dashed line) is independent of the activity of the added neuron. Intuitively, to keep the information unchanged, the ratio between the noise magnitude along 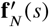 and the distance between two stimuli, 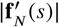, must remain the same as in the single-neuron case shown in Fig. 4A. That is, the standardized mean difference along the 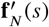 direction must be preserved. This intuition follows directly from the definition of LFI, which corresponds to the infinitesimal limit of the squared standardized mean difference along the direction that best discriminates the response distributions for the two stimuli (text S8).

Based on Eq. 15, we prove that, in general, the noise covariance of subsampled populations exhibiting increasing and saturating LFI with *I*_*N*_ < *I*_∞_ at every finite *N* can always be expressed in the form of Eq. 12. The proof follows by showing that 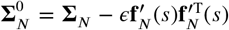 is positive definite for *ϵ* = 1/*I*_∞_ (for more details see text S9). In other words, such a noise covariance inherently contains differential correlations. This result justifies the widespread use of the concept of differential correlations in the analysis of subsampled populations (*24, 31, 32*).

Given the equivalence between differential correlation and information limiting under the subsampled conditions, we can reconfirm the necessity of the linear eigenvalues for the bounded LFI as a consequence of the differential correlations (*22, 31*). Namely, the addition of differential correlations induces linear eigenvalues in the covariance matrix. The number of linear eigenvalues depends on whether the LFI associated with 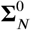 is unbounded for all signal directions or only for specific directions (text S10). However, to further achieve information boundedness, the alignment of the linear eigenvectors with the signal must occur faster than the rate set by the sublinear noise components (text S4) as well as the decay rate of the linear eigenvalue slopes with rank, i.e., boundedness requires each nonnegative LFI contribution to remain bounded. More importantly, the above proof mitigates the need to directly search for differential correlations to confirm or refute the boundedness of information.

Finally, this equivalence between information-limiting conditions and differential correlations is specific to subsampled populations. Differential correlations can be absent in more general noise covariances of non-subsampled neuronal populations, even when the LFI approaches a finite limit (e.g., when it approaches that limit from above; text S11). Such a scenario might occur in growing neuronal populations, for example through neurogenesis.

## Discussion

We proposed a theory of population coding based on the scaling properties of neural systems, which allowed us to predict the boundedness of information. This theory critically considers the constraints introduced by subsampling, leading to the following key findings: (i) The squared distance between mean responses of randomly subsampled *N* neurons to two close stimuli increases proportionally to *N* (*16, 18*). (ii) Random subsampling makes the total variance increase linearly with *N*, which allows a finite or countably infinite family of fixed-rank eigenvalue sequences to scale linearly, while eigenvalues whose ranks grow with *N* may scale sublinearly. (iii) For a finite linear family, the LFI can be limited if the sublinear eigenvalues’ angular occupation drops sufficiently rapidly. For an infinite linear family, the linear-eigenvalue contribution must additionally remain bounded. For a scale-invariant power-law eigenspectrum, these constraints further imply that (iv) an eigenspectrum with *α* > 0 or *α* = 0 necessitates that the spectrum decays with an exponent *μ* greater or less than one, respectively. (v) Only the scale invariance with *α* > 0 allows for linear eigenvalues, which can potentially limit the information. (vi) By contrast, *α* = 0 only allows for sublinear eigenvalues, resulting in unbounded LFI.

Through examination of the mouse data, we found the scale-invariant noise eigenspectra with *α* > 0 (Fig. 2D–F) and another power-law scaling relation between the eigenvalues and angular occupations with an exponent *γ* (Fig. 2G). Our analysis identified the critical exponent above and below which the LFI becomes bounded or remains unbounded. We found that the directions of the small noise components in the mouse data overlap with the signal direction at a sufficient rate, leading to a sublinear yet unbounded increase in LFI. Consequently, we conclude that the information encoded in the mouse V1 neurons can continue to increase without bound, going beyond the previous study on this dataset that reported no apparent signs of saturation in the LFI lower bound up to the recorded number of neurons (*21*).

Previous studies (*15, 16, 18, 21*) reporting bounded information in the limit of large networks are based on inadequate assumptions about covariance scaling. First, the study that collected the data analyzed here and reported high-precision coding (*21*) relied on extrapolation through a simple parametric scaling with 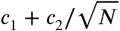 fitted to the decoding performance. Our results suggest that the reported bounded precisions in the limit of large *N* by a finite positive *c*_1_ were conservative estimates, as predicted by the authors. Second, ignoring or regularizing the activities related to smaller variances can lead to different and potentially inaccurate conclusions. Rumyantsev et al. reported saturating information using a dimension-reduction method (*16*). However, applying the same method to simulated data whose LFI is theoretically unbounded, we observed saturating LFI (text S7, figs. S9 and S10). Third, one cannot use a finite-size fit of Eq. 13 alone to infer bounded LFI. It is possible to find a saturating function that fits finite-size data well because *I*_0_ may be any increasing unbounded function and *ϵ* is unknown (text S9). Therefore, excellent fits of Eq. 13 to empirical LFI (*15, 16, 18*) cannot, by themselves, establish asymptotic boundedness.

We found scale invariance with a positive *α* and a fast-decaying power-law exponent *μ*(= 1 + *α*) > 1 in the recorded neural activities (Fig. 2D–F, and figs. S1–S5D–F). In addition, we observed a scaling exponent *μ* < 1 from the covariances of normalized neural activities (fig. S8), providing further evidence for unbounded information coding. This latter result aligns with other studies that have reported scale-invariant eigenspectra with scaling exponents less than 1 (*4–6, 33*). While these results might indicate that noise in neural activities does not generally confound information coding, further rigorous assessment of the scaling parameters *α* and *μ* across brain regions and species is necessary to reveal the foundational coding capacities of neurons.

The ongoing activity in the mouse forebrain, including the visual cortex, is high-dimensional (*1*). Furthermore, its reliable dimension increases without apparent saturation up to about a million neurons (*3*). These findings are consistent with the high-dimensional noise covariances because the ongoing activities driven by the animal’s behavior and internal dynamics are likely to cause variability in repeated stimulus-evoked activities (*34–34*). The above large-scale analyses showed that the ongoing activities overlap with signal correlations through one- or few-dimensional subspaces, suggesting multiplexed coding by the nearly orthogonal stimulus and spontaneous activities. The present study also supports that the noise correlations, including those arising from measurement noise, do not limit local signal representation. However, in agreement with positive correlations between the signal and noise in early sensory areas reported in other studies (*35–35*), we found broader directional overlaps between the leading noise components and signal (Fig. 2G), which is nevertheless insufficient to limit the information.

More generally, it remains unclear how the noise under brain states and behavioral conditions other than passive visual stimulation affects large-scale population coding. In smaller populations, it is known that cortical states (*40*), locomotion (*17, 41*), and attention (*42*) modulate noise correlations in a way that improves coding capacity. We showed that population coding is affected by the scaling of smaller noise variances in the population activity and how their directions align with the signal direction. Thus, we cannot rule out the idea that even small changes caused by the behavior and internal dynamics (*5*) in small noise components of the high-dimensional noise have a substantial impact on the stimulus coding fidelity in large-scale neuronal populations.

## Materials and Methods

### Analysis of mouse V1 data

We analyzed two-photon calcium imaging data of mouse V1 neurons from the “full-field” dataset by Stringer et al. (*21*) (see Data, code, and materials availability). The dataset includes recordings from five awake head-fixed mice, passively viewing static grating stimuli with up to ~1000 trials per degree. Each trial presented a static grating stimulus for 750 ms, followed by a gray screen for ~650 ms. The static grating stimuli shared the same spatial frequency (0.05 cycles per degree) with orientations ranging from 43° to 47°. The individual neural activities extracted by the Suite2p toolbox consisted of deconvolved fluorescence traces responding to the 750 ms stimulus presentation.

We divided the trials into two groups based on the orientation of the grating stimuli. Trials with orientations from 43° to 45°, including 45° (*s*_1_), comprise the first group, while trials with orientations above 45° and up to 47° (*s*_2_) comprise the second. The numbers of trials for Stimuli 1 and 2, *L*_1_ and *L*_2_, differed by a small margin in all datasets (< 50 trials in each dataset). We balanced the number of trials for both stimuli by randomly choosing min{*L*_1_, *L*_2_} trials (without replacement) from the stimulus with the larger number of trials. Following this procedure, the number of trials was 1855– 2654. Each dataset had *N*_*t*_ = 17,631–21,469 neurons.

We computed the activation means and covariance as follows. Let 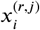 denote the activity of neuron *i* on trial *r* (= 1, …, *R*) for stimulus group *j* ∈ {1, 2}. For each group, we estimated the trial-averaged population activity as

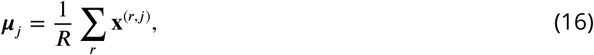

and the covariance matrix as

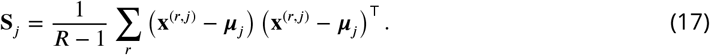

We defined the unscaled mean-response difference as *δ****μ*** = ***μ***_2_ − ***μ***_1_ and estimated 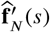 as

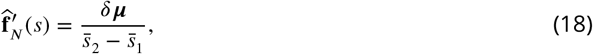

where 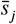 denotes the trial-averaged stimulus orientation within group *j*. The population covariance was estimated as

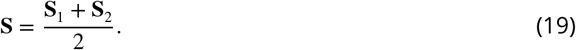

To characterize the eigenvalues and angular occupations for a subsample of size *N*, we randomly selected 200 subpopulations of size *N* and averaged the resulting eigenvalues and angular occupations.

Since the raw activity traces of neurons might include confounding factors such as different expression levels of fluorescence proteins or positions of neurons in the scanning system, we also constructed the activation means and covariance using the standardized data, by normalizing each neuron’s activity across all trials by its standard deviation (*4, 5*) or by the mean of its nonzero activities (*6*).

### Fitting scaling parameters and a scale-invariant function to an eigenspectrum

The scaling parameter of the scale-invariant eigenvalues (Eq. 5) was fitted as follows. The eigenvalues were evaluated at population sizes spaced by 50 neurons. Given the scale invariance, we must have:

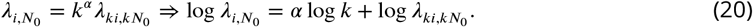

For each reference size *N*_0_, we used *k* = 2, …, *K*(*N*_0_), where *K*(*N*_0_) = ⌊*N*_avail_/*N*_0_⌋ and *N*_avail_ is the largest available population size. Therefore, we define the following cost function for *α* as

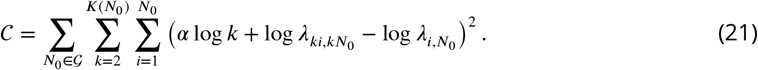

The minimum of the cost function is obtained as a solution of 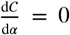 from which we obtain an estimated 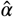:

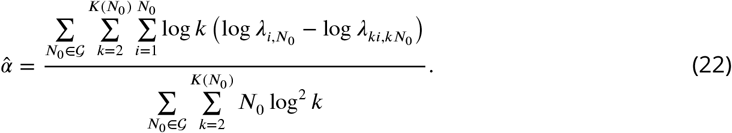

We computed 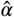 for each recording using Eq. 22. The reference-size grid *G* began at 50 and extended up to one quarter of the maximum trial number.

We used the jackknife resampling method to obtain the estimation error of *α*. For each *r* = 1, 2, …, *R*, jackknife replicates 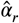 were generated by computing the mean eigenvalues 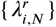 over all but the *r*th repetition, and using them in Eq. 22. Then, the jackknife estimate of the standard deviation of 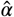 was obtained as 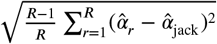, where 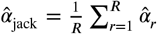.

Using the estimated *α*, we obtained a collapsed curve by plotting *N*^*α*^*λ*_*i,N*_ against *x*_*i*_ = *i*/*N*, using the mean eigenvalues over *R* repetitions. We then fitted the parametric scale-invariant function *g*(*x*_*i*_) (Eq. 6) to the collapsed eigenspectra in logarithmic coordinates, using all ranks and all evaluated population sizes.

The scaling parameter *γ* (Eq. 7) was estimated as follows. Using *t* randomly chosen trials for both stimuli (where *t* was varied from 500 up to the largest number of trials in the recording), we obtained *γ*(*t, N*) as the slope of a straight line fit to cos^2^(*θ*_*i,N*_) vs *λ*_*i,N*_ in the log-log plot. For each *N*, we estimated *γ*_∞_(*N*) as the y-intercept of a straight line fit to the plot of *γ*(*t, N*) vs 1/*t*. The final estimate 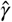 and its standard deviation were obtained as the average and standard deviation of *γ*_∞_(*N*) for *N* > 1000.

### LFI scaling under a power-law relation between angular occupation and eigenvalues

Here we outline the derivation of the critical exponent *γ*_*c*_ (Eq. 8) and the LFI scaling (Eq. 9); the full proof is given in text S3. Combining the LFI (Eq. 1) with the scale-invariant eigenspectrum *λ*_*i,N*_ = *N*^−*α*^*g*(*x*_*i*_) (*α* > 0; Eq. 5) and the power-law relation between angular occupation and eigenvalues (Eq. 7) gives

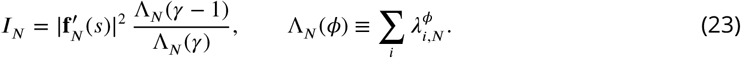

Using the leading-edge power law *g*(*x*) ~ *x*^−(1+*α*)^, the integral approximation yields

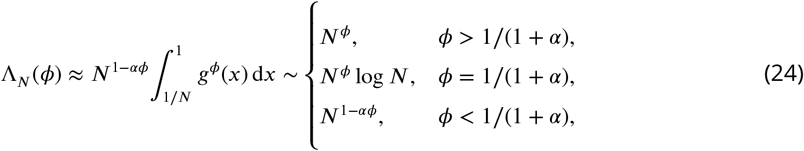

where the case split reflects whether the lower integration limit makes the integral diverge with *N*. Since Λ_*N*_ (*γ*) ~ *N*^*γ*^ and 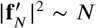 for *γ* > 1, we have *I*_*N*_ ~ *N*^1−*γ*^ Λ_*N*_ (*γ* − 1); substituting *ϕ* = *γ* − 1 then reproduces Eq. 9, with the transition at *γ* − 1 = 1/(1 + *α*), i.e. *γ*_*c*_ = (2 + *α*)/(1 + *α*) (Eq. 8).

### Bias correcting the eigenvalues and angular occupations

#### Eigenvalue bias correction

Finite-sample estimates of covariance matrices are known to produce biased eigenvalues: small eigenvalues are typically underestimated, while large ones are overestimated (*43*). To address this, we developed an iterative shrinkage method. We began by computing a naive estimate of the covariance matrix, 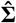. This estimate was treated as a sample from a Wishart distribution

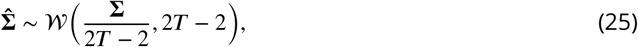

where **Σ** is the true covariance and *T* is the number of trials. Our goal was to infer the set of true eigenvalues, 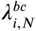, that could produce the observed biased sample eigenvalues.

Bias-corrected eigenvalues were obtained iteratively in the principal component basis, in which the true covariance is diagonal. Starting with the naive eigenvalues, 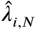, as the initial guess, we repeatedly sampled synthetic covariance matrices from the Wishart distribution with the current covariance iterate as its mean:

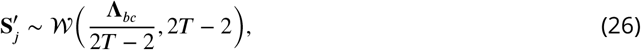

where **Λ**_*bc*_ is the diagonal matrix of current eigenvalue estimates. The ranked eigenvalues of these synthetic matrices were averaged across samples to obtain 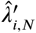. These averages were then compared with the naive estimates, and the bias-corrected eigenvalues were updated according to

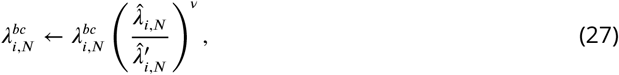

where *ν* is a small exponent controlling the update step.

This process was repeated until convergence, at which point the averaged synthetic eigenvalues matched the naive estimates. The resulting 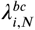 provide a bias-corrected spectrum that removes finite-sample bias from the covariance matrix. A detailed description of the method is presented in text S12.

#### Angular occupation bias correction

The error in estimating the angles *θ*_*i,N*_ between the vector *δ***f** and the eigenvectors of the covariance matrix **u**_*i,N*_ leads to biased estimates of the angular occupations cos^2^ *θ*_*i,N*_. This bias arises from two sources. First, the estimated eigenvector *û*_*i,N*_ is oriented differently from the true eigenvector **u**_*i,N*_. Second, the estimated vector *δ****μ***, used as an approximation of 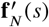, has a different orientation from the true vector. We denote 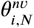, the naive estimate of *θ*_*i,N*_, as the angle between *δ****μ*** and 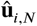. Similarly, we denote the angle between *δ****μ*** and the actual eigenvector **u**_*i,N*_ as 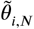. To correct the bias in angular occupations, we use a two-step method. First, we make the correction resulting from the error in the estimate of **u**_*i,N*_; the goal of this step is to find 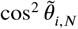. In the second step, we correct 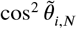 for the bias caused by the error in the estimation of the signal. A detailed description of this step is presented in text S13.

##### Eigenvector bias correction

In the first step of angular occupation bias correction, we only have access to the naive estimates of the squared cosines 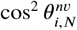. To find 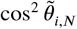, we use an iterative method similar to the eigenvalue bias correction. Let 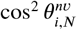 denote the naive angular occupation estimates obtained from the estimated eigenvectors over *m* different subpopulations of the same size *N*. The goal of the algorithm is to use these naive estimates to recover bias-corrected values 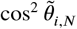 that more accurately reflect the alignment with the true covariance eigenvectors. We work in the same framework as in eigenvalue bias correction.

The algorithm proceeds as follows. First, the naive estimates are used as an initial guess for the bias-corrected squared cosines. We then generate *m* covariance matrices 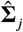 from a Wishart distribution,

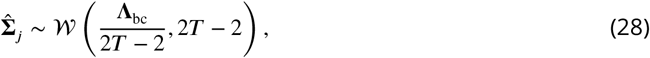

where **Λ**_bc_ is the diagonal matrix of bias-corrected eigenvalues obtained from the eigenvalue correction procedure. For each simulated covariance matrix, we compute the eigenvectors and their squared cosines with the candidate signal direction defined by the current angular-occupation estimates. These values are averaged across all *m* samples to produce predicted squared cosines 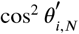 that represent the expected naive estimates under the current guess of the true cosines.

The bias-corrected squared cosines are then updated according to

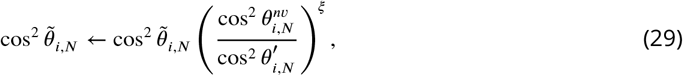

where *ξ* serves as a learning rate controlling the update magnitude. The squared cosines are subsequently normalized so that

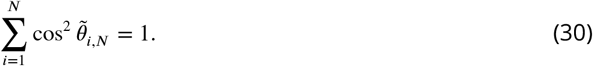

Steps of simulation, averaging, updating, and normalization are repeated until convergence, defined as the condition 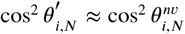 is satisfied for all *i*.

This procedure yields bias-corrected squared cosines that account for sampling variability in the estimated eigenvectors and provide a more accurate representation of the alignment between *δ****μ*** and the population eigenvectors. The resulting estimates can then be used in subsequent analyses requiring precise directional information, such as perturbative bias corrections or projections along principal components.

##### Signal bias correction

We modeled the observed signal difference *δ****μ*** as the sum of the true underlying difference *δ***f** and additive Gaussian noise,

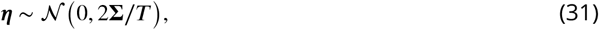

where **Σ** is the known noise covariance and *T* is the number of trials. The eigenvectors **u**_*i,N*_ and eigenvalues *λ*_*i,N*_ of **Σ** were used to define principal directions of variability, and the alignment of the signal difference with each eigenvector was quantified by the squared cosine cos^2^ *θ*_*i,N*_. We define

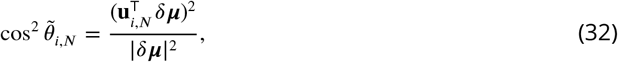

which captures the fraction of the signal along each direction but is biased due to measurement noise.

To correct for this bias, we performed a perturbative expansion in 1/*T*, which shows that the expected value of the naive squared cosine is shifted by terms proportional to *λ*_*i,N*_. Neglecting higher-order terms, the bias-corrected estimate is

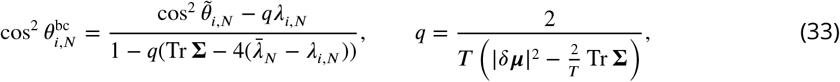

where 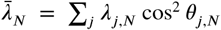. Because 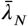 depends on the cos^2^ *θ*_*i,N*_ values themselves, we implemented a few iterations in which 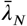 was updated using the current estimates of cos^2^ *θ*_*i,N*_. During iteration, any negative bias-corrected values were clipped to zero. This procedure yields nonnegative signal fractions along each principal direction that account for leading-order finite-sample noise.

### Scaling relations under linear scaling of the total variance

We sketch the derivation of the scaling relations summarized in Table 1; the full derivation is given in text S5. We write a scale-invariant eigenspectrum with a power-law leading edge as

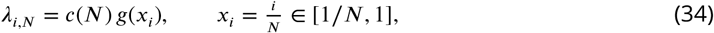

where *x*_*i*_ denotes the rank fraction. For small *x* we assume a power-law leading edge,

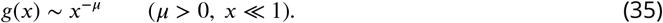

The linearly scaling total variance then reads

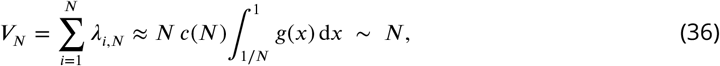

Thus, the *N*-dependence of the integral, and hence the constraint on *c*(*N*), is determined by the small-*x* behavior of *g* through the lower integration limit:

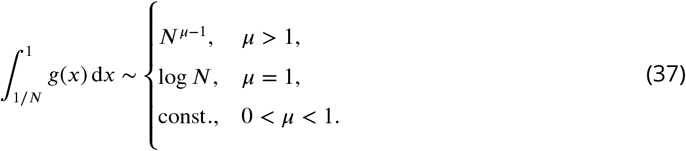

For *μ* > 1 this forces *c*(*N*) = *N*^−*α*^ with *α* = *μ* − 1 > 0, giving linearly scaling leading eigenvalues *λ*_*i,N*_ ~ *N*^*μ*−*α*^ = *N*. Such linear noise modes can bound the LFI only if the (also linearly growing) signal direction aligns with them; otherwise, the LFI is unbounded. For 0 < *μ* < 1 the integral converges, so *c*(*N*) is constant (*α* = 0) and the leading eigenvalues scale sublinearly, *λ*_*i,N*_ ~ *N*^*μ*^. In the absence of linearly scaling noise modes, the LFI is unbounded under the linear scaling of the signal 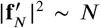. The two regimes correspond to the rows of Table 1 (*α* > 0 ⇒ *μ* = 1 + *α*, linear; *α* = 0 ⇒ *μ* < 1, sublinear), with *μ* = 1 a marginal case (*c*(*N*) ~ (log *N*)^−1^).

### Simulation of neural activity with bounded and unbounded LFI

We simulated neural responses to two stimuli by sampling from two multivariate Gaussian distributions, with means **f**_1_ = diag(**Σ**) and **f**_2_ = **f**_1_ + Δ**f** and common covariance **Σ**:

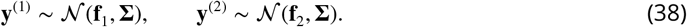

We chose **f**_1_ to approximate a Poissonian mean–variance relationship. We generated neural activity separately for each population size. The eigenvalues of **Σ** were obtained from Eqs. 5 and 6 for *α* > 0, and from the modified *α* = 0 eigenspectrum described in the main text.

We constructed **Σ** using the following procedure: (1) sample reference firing rates *r*_*i*_ from a lognormal distribution fitted to the mean responses in mouse TX42 data, (2) construct a positive-definite candidate matrix **M** using the square root (elementwise) of the outer product of the sampled reference firing rates, with diagonal entries *r*_*i*_ and off-diagonal entries 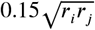 (3) sample from the inverse-Wishart distribution with **M** as the mean; and (4) obtain eigenvectors **U** of the resulting matrix and combine them with the diagonal matrix **Λ** of prescribed eigenvalues to construct **Σ** = **UΛU**^T^. Using these eigenvectors, we generated the mean-response difference Δ**f** using the positive square roots of the angular occupations given by Eq. 7. Specifically, we set 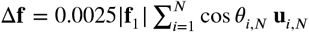.

## Ethics statement

This study used only previously published, publicly available data (*21, 44*) and involved no new animal experiments. All original procedures were conducted under protocols reviewed by the IACUC, as reported in (*21*).

## Acknowledgments

The authors thank Rubén Moreno Bote, Kenneth Harris, and Quan Wen for valuable discussion, and Magalie Tatischeff and Bingyue Zhu for preliminary data analysis.

## Author contributions

S.A.M. and H.S. designed the research; S.A.M., S.S.R.H., and H.S. performed the theoretical analysis. S.S.R.H. and S.A.M. performed the data analysis; H.S. supervised the work. All authors contributed to writing the manuscript.

## Data, code, and materials availability

All data needed to evaluate the conclusions in the paper are present in the paper and/or the Supplementary Materials. The two-photon calcium imaging recordings analyzed in this study are publicly available from Pachitariu, Michaelos, and Stringer (2019), “Recordings of 20,000 neurons from V1 in response to oriented stimuli”, https://doi.org/10.25378/janelia.8279387. The code for the data analysis is archived on Zenodo (https://doi.org/10.5281/zenodo.19504768) and linked to the GitHub repository: https://github.com/Sai-Sumedh/scaling-population-coding. This study did not generate new materials.

## Supplementary Materials

## text S1. Linear Fisher information

In this note, we present the eigenvector representation of the linear Fisher information (LFI) for neuronal populations. This expression provides the basis for our later analysis of how covariance scaling and signal alignment determine whether information remains bounded or grows without bound.

Let **f**_*N*_ (*s*) be an *N*-dimensional vector whose elements are mean activity or tuning functions *f*_*i*_(*s*) (*i* = 1, …, *N*) of individual neurons at a given stimulus *s*. Assuming that the covariance matrix of neuronal activity is invertible, the LFI about the stimulus *s* can be written as:

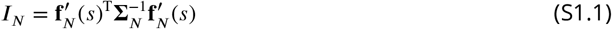

where 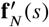 is the *N*-dimensional vector composed of the derivatives of tuning functions with respect to *s*, assuming that the tuning functions are differentiable, and **Σ**_*N*_ is the covariance matrix.

We denote the *i*-th eigenvalue of the covariance matrix **Σ**_*N*_ in the *N*-dimensional space as *λ*_*i,N*_. Here we index the eigenvalues in a descending order (from large to small): *λ*_1,*N*_ ≥ *λ*_2,*N*_ ≥ ⋯ ≥ *λ*_*N,N*_. We write the covariance matrix in terms of its eigenvalues *λ*_*i,N*_ and eigenvectors **u**_*i,N*_ as

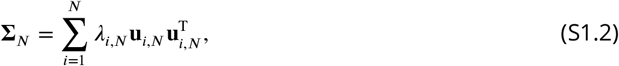

where T is the transposition operator. We can also write 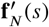 as a linear combination of eigenvectors of the covariance matrix as

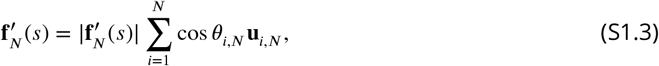

where *θ*_*i,N*_ is the angle between the vector 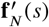 and the *i*-th eigenvector of the covariance matrix **u**_*i,N*_. Using the above equations, we can rewrite the LFI as

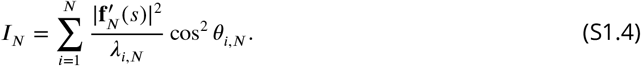

Eq. S1.4 indicates that whether the LFI is limited or not depends on the combination of the scaling of 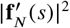, the eigenvalue *λ*_*i,N*_, and the angular occupation cos^2^ *θ*_*i,N*_. Each obeys a certain constraint, resulting in a characteristic scaling structure. Most importantly, the signal magnitude 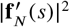 is expected to scale with *N* as long as each element 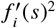 is sampled from a common distribution having a finite nonzero mean *η* because 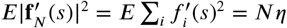.

Although the LFI behavior with respect to the population size *N* generally depends on how the eigenvalues and angular occupations scale relative to the linearly scaling signal magnitude, a simple conclusion can be drawn without knowing the angular occupations when all eigenvalues scale sublinearly. In this case, the largest eigenvalue satisfies *λ*_1,*N*_ /*N* → 0 as *N* → ∞. Since *λ*_*i,N*_ ≤ *λ*_1,*N*_ and 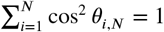, Eq. S1.4 implies

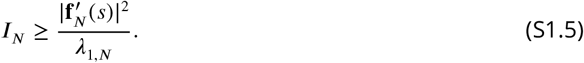

If 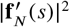 scales linearly with *N*, the right-hand side diverges. Thus, in this special case, the LFI is unbounded regardless of the signal direction.

## text S2. Linear scaling of the total variance constrains the scaling of eigenvalues

In this note, we discuss the linear scaling of the total variance in subsampled neurons, categorize the scaling behavior of eigenvalues, and outline constraints on eigenvalue scaling when the total variance scales linearly.

For any number of neurons *N*, the total variance *V*_*N*_ is equal to the sum of variances of individual neurons, which is also equal to the sum of all eigenvalues of the covariance matrix, i.e.,

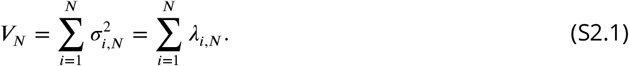

Here *σ*_*i,N*_ is the standard deviation in the activity of neuron *i* in a population of size *N*. In a sub-sampled population of neurons, the variance of an individual neuron’s activity is independent of the subpopulation size *N*, i.e., 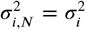. We can consider a distribution of 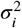 across neurons with a positive mean 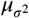 and a bounded support, i.e., the variance of a given neuron is nonnegative by definition and cannot exceed an upper bound by physiological constraints. Looking at the variability in the activity of a randomly selected neuron in recordings of a neuronal population is equivalent to sampling 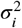 from its distribution. Under this random-sampling model, the law of large numbers gives 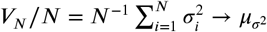 as *N* → ∞. Because 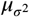 is finite and nonzero, we have

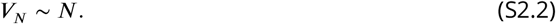

This adds important constraints on the scaling of the eigenvalues with respect to *N*.

To see this, we categorize the eigenvalues into those that scale linearly, sublinearly, and supralinearly, defined as follows. For each fixed rank *j* ∈ N, consider the eigenvalue sequence {*λ*_*j,N*_}_*j*≤*N*_, and assume that lim_*N*→∞_ *λ*_*j,N*_ /*N* exists as an extended nonnegative real number. The set of indices corresponding to linearly scaling eigenvalues is defined as

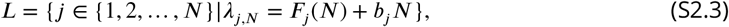

where *b*_*j*_ > 0 is a constant positive number, and *F*_*j*_(*N*) is a function satisfying

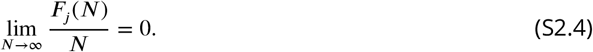

Here, we refer to *λ*_*j,N*_ = *F*_*j*_(*N*) + *b*_*j*_*N* as linear eigenvalues because the linear term will dominate the behavior at large *N*. Next, we define a set of sublinear eigenvalues that grow slower than the linear function in the limit as

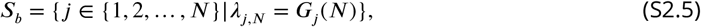

where *G*_*j*_(*N*) is a function satisfying

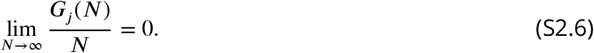

The sublinear eigenvalues may increase sublinearly, remain constant, or decrease as a function of *N*. Similarly, we define a set of indices corresponding to supralinear eigenvalues as

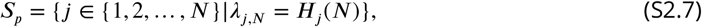

in which *H*_*j*_(*N*) is a function of *N* with the following property

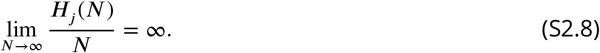

Given these definitions, an immediate consequence of the constraint given by Eq. S2.2 is that the supralinear fixed-rank eigenvalue sequences cannot exist under the linear scaling of the total variance. For linear eigenvalues, if *b*_*j*_ values exhibit a constant positive lower bound *b*_*j*_ > *b*_0_ > 0, the number of linear eigenvalues must converge to a fixed finite number as *N* increases, i.e., the set *L* is finite. Otherwise, if the number of linear eigenvalues increases with *N*, their summed contribution to the total variance grows faster than linearly. For example, if their number grows as *N*^*ϕ*^, the lower bound *b*_*j*_ > *b*_0_ implies that their summed contribution is at least *b*_0_*N*^*ϕ*+1^, which is supralinear for *ϕ* > 0. Thus, the number of linear eigenvalues must remain finite. On the other hand, if the *b*_*j*_ values decrease with rank sufficiently rapidly to be summable, then the contribution of the linear eigenvalues is compatible with linear total-variance scaling. For example, *b*_*j*_ ~ *j*^−(1+*α*)^ is summable when *α* > 0.

In sum, at large *N*, if *b*_*j*_ > *b*_0_ > 0, the number of elements in set *L* approaches a constant integer *c* ≥ 0, and consequently the number of elements in set *S*_*b*_ linearly increases with *N*, while the set *S*_*p*_ becomes empty, i.e.,

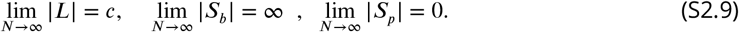

On the other hand, if *L* is countably infinite, linear total-variance scaling requires its positive slopes to be summable:

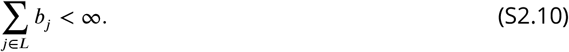

## text S3. Scaling of LFI under a scale-invariant eigenspectrum with a power-law relation to the angular occupation

Considering the scaling ansatz in Eqs. 5 and 6 with *α* > 0 together with the following power-law relation (Eq. 7):

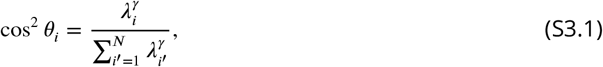

we can write the LFI as

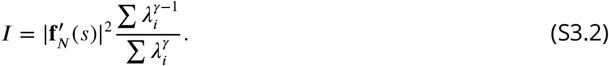

In this note, we establish an analytical expression of the above LFI and prove that the information is a bounded function of *N* if

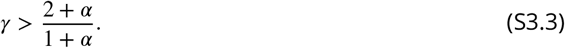

Before proving the above statement, we argue, more generally, that regardless of the scaling of the eigenvalues, if *γ* = 1, we have

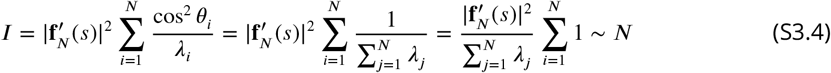

Therefore, for *γ* = 1, the information is a linearly increasing function of *N*. The last step is obtained by considering the fact that

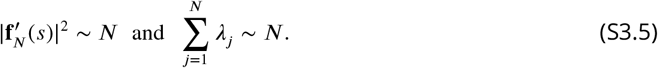

See text S2 for more details. On the other hand, if *γ* = 2,

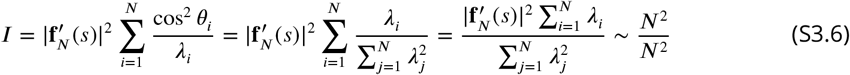

that is a bounded function of *N*. It is clear that the numerator, which is the product of the total variance and the squared norm of 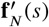, scales as *N*^2^. The denominator scaling, however, needs more attention. Since

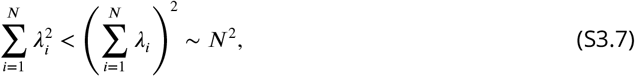

we have 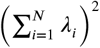 as an upper bound of 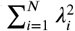. In addition, if there exists at least one eigenvalue that is a linearly increasing function of *N*, say *λ*_1_ ~ *N*, then we have

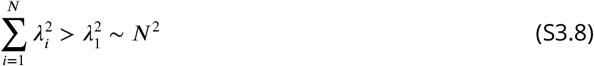

that specifies a lower bound for 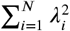; as a result, because both the upper and lower bounds scale with *N*^2^, we can conclude that

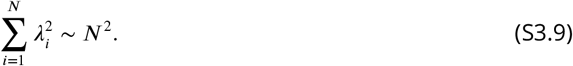

Up to this point, we showed that for *γ* = 1 the information is unbounded and for *γ* = 2 it is bounded. An immediate conclusion is that there must exist a 1 < *γ* < 2 at which we have a transition from unbounded to bounded information. In the rest of this note, we prove that the critical value of *γ* = *γ*_*c*_ at which the transition happens is identified by the following exponent relation:

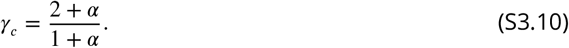

To prove that, we first focus on the scaling of

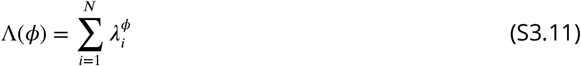

in which *ϕ* is a positive exponent. Considering the scaling ansatz (Eq. 5) with the scale-invariant function *g*(*x*) defined in Eq. 6, we can write

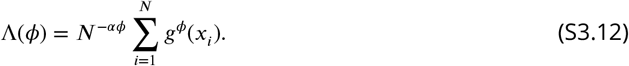

Using the integral approximation of the summation, we have

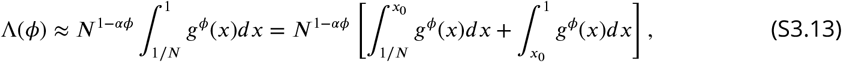

in which *x*_0_ is a fixed small number (1/*N* < *x*_0_ ≪ 1) for which we can approximate

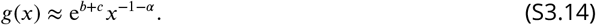

In Eq. S3.13, the second integral in the brackets is independent of *N*, and the first integral can be approximated as

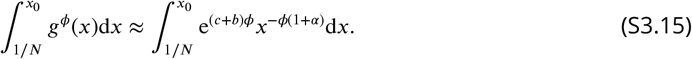

By integrating directly and substituting the result into Eq. S3.13, we get

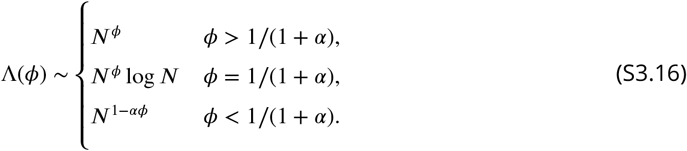

See below for the derivations.

Using Eqs. S3.2 and S3.16, we can rewrite the LFI in terms of the function Λ(*γ*) and Λ(*γ* − 1) as

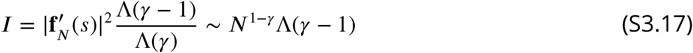

because Λ(*γ*) ~ *N*^*γ*^ for *γ* > 1. Using Eq. S3.16, we can write

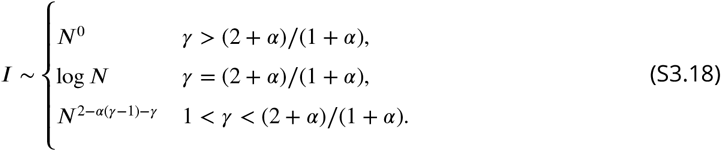

That is an unbounded function of *N* for 1 < *γ* ≤ (2 + *α*)/(1 + *α*) and bounded for *γ* > (2 + *α*)/(1 + *α*).

Finally, we note that cos^2^ *θ*_*i,N*_ is computed as

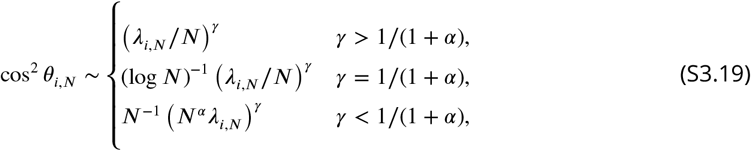

which exhibits the scale invariance except for *γ* = 1/(1 + *α*):

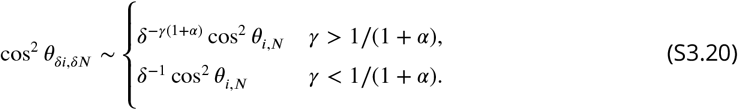

Derivation of Λ(*ϕ*) is as follows:

**Case of** *ϕ* > 1/(1 + *α*):

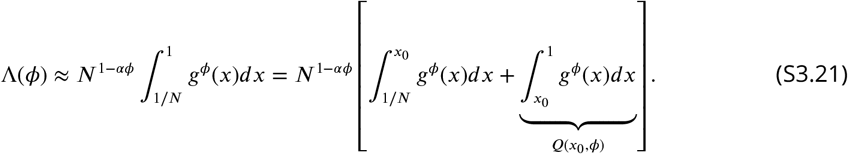

Using the approximation of *g*(*x*) for small *x*, we have

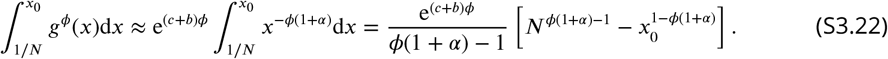

Replacing in Eq. S3.21, we get

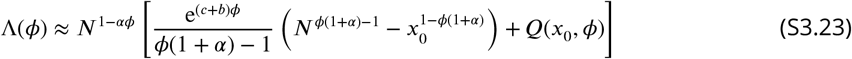

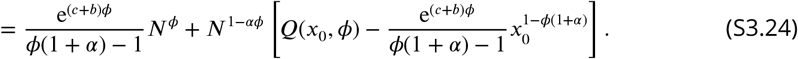

Because *ϕ* > 1 − *αϕ*, in the large-*N* limit

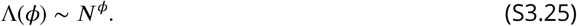

**Case of** *ϕ* = 1/(1 + *α*):

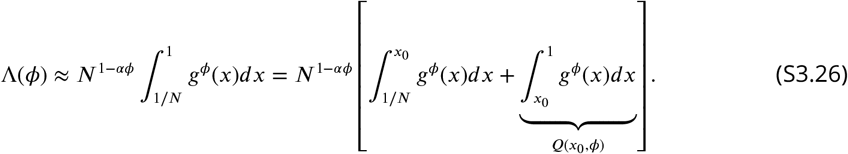

Using the approximation of *g*(*x*) for small *x*, we have

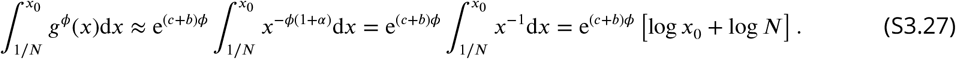

Replacing in Eq. S3.26, we get

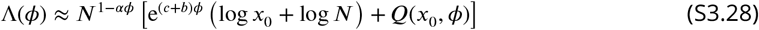

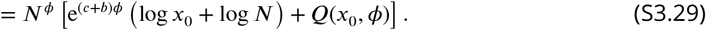

In the last step we used 1 − *αϕ* = *ϕ*. Therefore, in the large-*N* limit

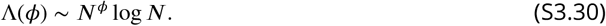

**Case of** *ϕ* < 1/(1 + *α*):

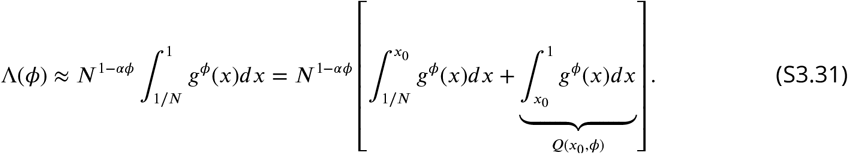

Using the approximation of *g*(*x*) for small *x*, we have

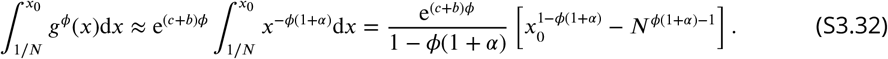

Replacing in Eq. S3.31, we get

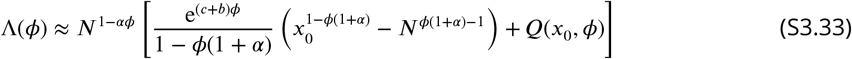

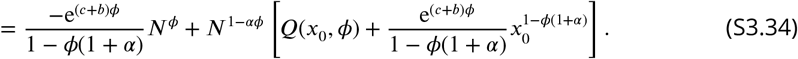

Because *ϕ* < 1 − *αϕ*, in the large-*N* limit

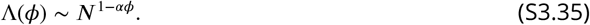

## text S4. Scaling conditions for bounded and unbounded LFI

In this note, we derive the necessary and sufficient conditions on the covariance matrix that limit information about a stimulus when the size of a neuronal population *N* increases, given that the covariance matrices exhibit the linear scaling of the total variance with respect to *N*. Here, we assume the slopes of linear eigenvalues exhibit a positive lower bound that leads to a finite number of linear eigenvalues under the linear scaling of the total variance.

First, as explained in text S1, 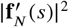 is expected to scale with *N*

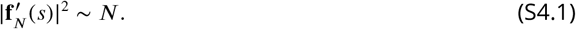

Second, the angular occupation is constrained by

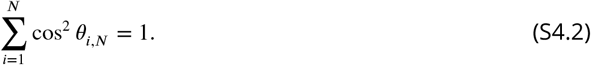

We will explain the consequences of this constraint on angular occupation at the end of this note.

Third, the total sum of eigenvalues for the subsampled population scales linearly:

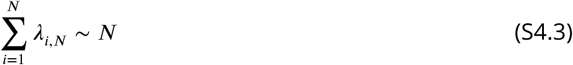

As discussed in text S2, under the assumed positive lower bound on the linear-eigenvalue slopes, this constraint requires |*L*| to remain finite and |*S*_*b*_| to increase linearly with *N*, while prohibiting supralinear eigenvalues. Throughout this note, |*L*| denotes the set of linearly scaling eigenvalues, and we assume |*L*| < ∞.

The LFI based on the above definitions can be written in terms of the contributions of the two defined sets as

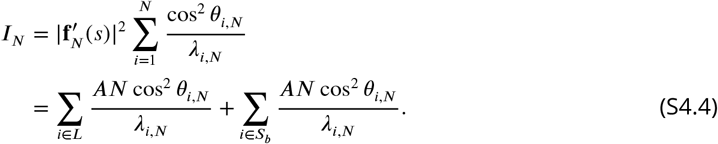

where we used 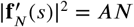. For future reference, we define

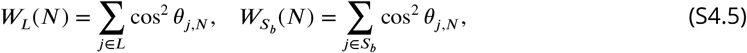

as the total angular occupation by these two sets.

Because the two summation terms on the right-hand side of Eq. S4.4 are nonnegative, the LFI is bounded if and only if both terms remain bounded. Here, we study the two terms. Using the linear-eigenvalue definition *λ*_*i,N*_ = *F*_*i*_(*N*) + *b*_*i*_*N* for *i* ∈ *L* from text S2 (Eq. S2.3), together with cos^2^ *θ*_*i,N*_ ≤ 1, the first term is bounded as

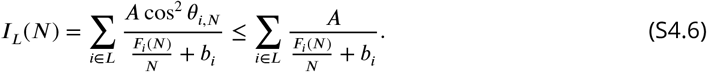

Hence, applying the definition of the sublinear function (Eq. S2.4), individual terms in the summation on the right-hand side of the equation above approach a finite value in the large-*N* limit. Since the number of linear eigenvalues |*L*| is finite by the finite-*L* assumption of this note, their sum is finite. Therefore, *I*_*L*_(*N*) is bounded, regardless of the behavior of cos^2^ *θ*_*i,N*_ as a function of *N*.

The second term on the right-hand side of Eq. S4.4 is

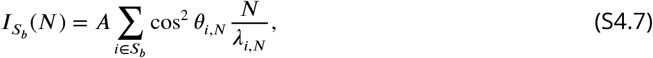

where *λ*_*i,N*_ = *G*_*i*_(*N*) for *i* ∈ *S*_*b*_ (Eq. S2.6). Based on the definition of 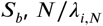 in the above summation are unbounded functions of *N*. Therefore, to have an upper bound for 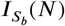, it is necessary that for all terms in this summation, cos^2^ *θ*_*i,N*_ must decay asymptotically at least as fast as *λ*_*i,N*_ /*N*. Not only that, the summation itself must remain bounded. In other words, to have an upper bound on 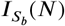, the required asymptotic decay of cos^2^ *θ*_*i,N*_ (∀*i* ∈ *S*_*b*_) as a function of *N* depends not only on the behavior of *λ*_*i,N*_ /*N* but also on the number of terms in the summation, i.e., the number of elements in *S*_*b*_. Note that, under the assumptions of this note, the number of sublinear noise components |*S*_*b*_| increases linearly with *N*.

Accordingly, the following two conditions comprise the necessary and sufficient conditions for bounded information. First, 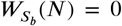 is sufficient for information to be bounded. Second, if 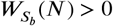, the second term can be written as

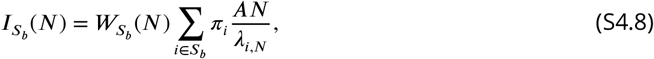

where

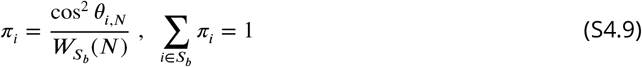

forms the probability mass function. Therefore, the summation term in Eq. S4.8 is a weighted average whose smallest term is *AN*/*λ* _|*L*| +1,*N*_ because the eigenvalues are ordered. Since *λ* _|*L*| +1,*N*_, the largest sublinear eigenvalue, grows more slowly than *N*, this smallest term grows without bound, and therefore so does the weighted average. Below, we denote this expectation by ⟨⋅⟩. Thus the necessary and sufficient condition that 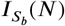 (as well as *I*_*N*_) is bounded is

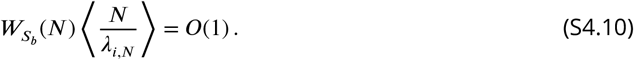

Here and below, *O*(*N*^*p*^) denotes a term whose magnitude divided by *N*^*p*^ remains bounded as *N* → ∞; *O*(1) corresponds to *p* = 0. Note that due to the definition of the sublinear eigenvalues and finiteness of *L*, ⟨*N*/*λ*_*i,N*_ ⟩^−1^ tends to zero in the limit. Thus, 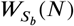 must converge to zero at a rate equal to or faster than ⟨*N*/*λ*_*i,N*_ ⟩^−1^.

In summary, the necessary and sufficient condition that the information is bounded is that either 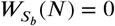 or 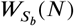 converges to zero at a rate equal to or faster than ⟨*N*/*λ*_*i,N*_ ⟩^−1^. The LFI is unbounded if and only if 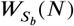 fails to approach zero rapidly enough to compensate for the growth of *N*/*λ*_*i,N*_, so that their product does not remain bounded as *N* increases.

### Necessary condition

Consequently, the necessary condition for the bounded information is that 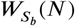 converges to zero, equivalently *W*_*L*_(*N*) converges to one:

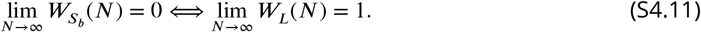

Further, one can consider a set that covers the linear eigenvalues and let the angular occupation by this set approach one to satisfy a necessary condition. We consider a positive integer *κ*(*N*) ≤ *N* such that a set of *i* = 1, …, *κ*(*N*) includes all linear eigenvalues. We define their angular occupation as 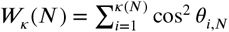. For the information to be bounded, it is necessary that *W*_*κ*_ (*N*) converge to one:

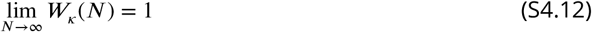

The correlations that violate this necessary condition result in unbounded information.

### Sufficient conditions

An exemplary sufficient condition for the bounded information is obtained as follows. Since *λ*_*i,N*_ ≥ *λ*_*N,N*_, we obtain the upper bound of ⟨*N*/*λ*_*i,N*_⟩ by replacing *λ*_*i,N*_ (*i* = 1, …, *N* − 1) with *λ*_*N,N*_ : ⟨*N*/*λ*_*i,N*_⟩ ≤ *N*/*λ*_*N,N*_. By multiplying 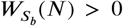, we have the upper bound of 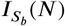

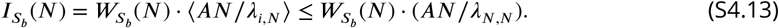

Thus, it is sufficient that 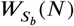 approach zero at least as fast as *λ*_*N,N*_ /*N*. Further, it is also sufficient to find a set containing all sublinear eigenvalues whose total angular occupation approaches zero at least as fast as *λ*_*N,N*_ /*N*.

Below, we present exemplary covariance matrices that yield bounded and unbounded LFI, assuming specific scaling conditions for the eigenvalues and angular occupation (except for Example 1).

### Example 1: Unbounded LFI

If the eigenvalues of the covariance matrix are all sublinear functions, we have 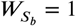, which clearly violates the necessary condition. Thus, the LFI for such populations is unbounded. Trial-shuffled covariance matrices provide a simple example. In the infinite-trial limit, shuffling makes the covariance matrix diagonal, so its eigenvalues are the individual-neuron noise variances. As assumed for subsampled populations in text S2, these variances are bounded above by a fixed constant independent of *N*. Hence, *λ*_1,*N*_ = *O*(1), so *λ*_1,*N*_ /*N* → 0 and all eigenvalues scale sublinearly.

### Example 2: Bounded/unbounded LFI

We consider only one sublinear (*i* = *N*) eigenvalue that has a finite cosine direction (cos^2^ *θ*_*N,N*_ > 0), while the rest of the sublinear eigenvalues have no angular occupation (cos^2^ *θ*_*i,N*_ = 0 for *i* ≠ *N* and *i* ∈ *S*_*b*_). The linear eigenvalues occupy the rest of the angular occupation: *W*_*L*_ = 1 − cos^2^ *θ*_*N,N*_. Then, for the LFI to be bounded, 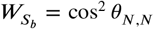 must decrease at a rate equal to or faster than *λ*_*N,N*_ /*N*. If cos^2^ *θ*_*N,N*_ = *N*^−δ^ and *λ*_*N,N*_ ~ *N*^*ζ*^ (*ζ* < 1), the exponent must satisfy −δ ≤ *ζ* − 1, namely δ ≥ 1 − *ζ*. Otherwise, if δ < 1 − *ζ*, the LFI is unbounded.

### Example 3: Bounded/unbounded LFI

We consider the first *N*_*c*_ eigenvalues to be linear, and the rest are sublinear eigenvalues given by *λ*_*i,N*_ ~ *N*^*ζ*^ with *ζ* ≤ 0 (*i* = *N*_*c*_ + 1, …, *N*). Note that *ζ* ≤ 0 is necessary for linearly scaling total variance. We then assign cos^2^ *θ*_*i,N*_ = *N*^−δ^ for *i* = *N*_*c*_ + 1, …, *N*. The angular occupation by sublinear eigenvalues is

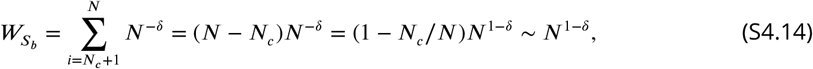

from which we find δ must satisfy δ ≥ 1 so that their total angular occupation does not exceed one as *N* grows. For the LFI to be bounded, 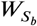 must decay at least as fast as ⟨*N*/*λ*_*i,N*_^−1^⟩ ~ *N*^*ζ*−1^. Thus if 1 − δ ≤ *ζ* − 1, namely δ ≥ 2 − *ζ*, the information is bounded. Otherwise, if δ < 2 − *ζ*, the LFI is unbounded.

As seen in these examples, one can construct the populations with bounded and unbounded LFI by appropriately assigning the angular occupations cos^2^ *θ*_*i,N*_ to eigenvectors associated with linear or sublinear eigenvalues. However, the assignment must be done by keeping the sum of the angular occupations one (Eq. S4.2). Here, we summarize its consequences, including the cases when the angular occupations scale with *N*.

1. Assuming monotonicity, the angular occupation must be either (i) a function of *N* approaching a nonzero constant equal to or less than one in the large-*N* limit, or (ii) a decreasing function of *N* that vanishes to zero in the large-*N* limit or a constant zero. For the following classification, assume that, for every fixed *j*, the angular occupation has a limit *c*_*j*_ ≡ lim_*N*→∞_ cos^2^ *θ*_*j,N*_.
2. Let *C* be the set with the nonzero angular occupations in the large-*N* limit:

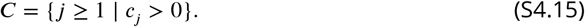

The set *C* may be finite or countably infinite, with ∑_*i*∈*C*_ *c*_*i*_ ≤ 1. For bounded LFI under the finite-*L* assumption, the necessary condition *W*_*L*_(*N*) → 1 implies *C* ⊆ *L*. Consequently, *C* is finite and ∑_*i*∈*C*_ *c*_*i*_ = 1. Some linear eigenvectors may have limiting angular occupation zero, so *C* may contain fewer eigenvectors than *L*. The angular occupations of the complement of *C* (*i* ∉ *C*) are zeros or functions vanishing to zero in the large-*N* limit.
3. At population size *N*, only the indices 1, …, *N* are present. Let *C*_*N*_ = *C* ∩ {1, …, *N*} denote those belonging to *C*. We define the total angular occupation by the set *C* as

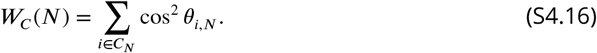

Suppose that, in the large-*N* limit, the components in *C*_*N*_ do not carry all the angular occupation. The remaining components must then carry a nonzero total occupation. More precisely, if *W*_*C*_ (*N*) → *w* < 1, then their total occupation, 1 − *W*_*C*_ (*N*), tends to 1 − *w* > 0. To illustrate how many small occupations can produce such a nonzero total, consider a group of *m*_δ_(*N*) components, each with angular occupation of order *N*^−δ^, with the same bounds applying to every component in the group. The group’s total occupation is of order *m*_δ_(*N*)*N*^−δ^. For this group to retain a positive total occupation, *m*_δ_(*N*) must grow in proportion to *N*^δ^.
4. In contrast, suppose *W*_*C*_ (*N*) → 1, so the total angular occupation of the remaining components, 1 − *W*_*C*_ (*N*), tends to zero. For a group with the same scaling as above, the total occupation is of order *m*_δ_(*N*)*N*^−δ^ and must now vanish. Therefore, unlike in the preceding case, *m*_δ_(*N*) must grow more slowly than *N*^δ^. In particular, if the group grows linearly, *m*_δ_(*N*) ∝ *N*, its total occupation vanishes when δ > 1.

These consequences can be utilized to set a range of scaling parameters for the angular occupations when constructing populations with the bounded and unbounded LFI. For example, for the LFI to be bounded, the nonzero limiting angular occupations must be assigned to linear eigen-values, and their sum, *W*_*C*_ (*N*), must converge to one to satisfy the necessary condition (Eq. S4.11). Example 3 applies item #4 by assigning the decreasing angular occupations (cos^2^ *θ*_*i,N*_ = *N*^−δ^) to the set of sublinear eigenvalues *i* = *N*_*c*_ + 1, …, *N*, whose number increases linearly. In this case, the decreasing angular occupations must scale with δ > 1, allowing us to set the range of the parameter δ. Note that this is a consequence of the necessary condition for bounded LFI under Eq. S4.2. The LFI is bounded only if δ ≥ 2 − *ζ*, as derived in Example 3.

Interestingly, under the finite-*L* assumption of this note, if *C* is an empty set, namely if all cos^2^ *θ*_*i,N*_ are decreasing functions of *N* that vanish to zero or constant zeros, the LFI is unbounded regardless of the composition of the eigenspectrum. Since the set *L* is assumed to be finite in this note, the sum of their angular occupations diminishes to zero, which violates the necessary condition (Eq. S4.11) for the bounded LFI. The trial-shuffled covariance matrices in Example 1, for which *L* = ∅, also illustrate the angular-occupation behavior described above. In the infinite-trial limit, their eigenvectors can be chosen as the single-neuron axes. For the axis associated with neuron *j*, the angular occupation is 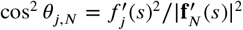. Since the individual squared signal components remain bounded while the total squared signal magnitude grows proportionally to *N*, every angular occupation is *O*(*N*^−1^) and vanishes. Thus, *C* = ∅, consistent with the unbounded LFI established in Example 1.

Finally, the necessary and sufficient condition for bounded LFI derived here assumes linearly growing total noise variance and a finite set *L* of linearly scaling eigenvalues. These assumptions constrain the covariance matrix at each *N*, but not how the covariance matrices at different population sizes are related. If the populations form a nested sequence created by successively adding neurons, each covariance matrix is contained in the next, and the LFI cannot decrease with *N* (text S8). Because the boundedness condition derived here does not require this nested structure, it also applies to more general covariance sequences, including those for which the LFI decreases with *N* (text S11).

## text S5. Scale-invariant eigenspectrum under the linear scaling of the total variance

In this note, we clarify the relationship between the scale invariance of the eigenspectrum, the exponent of the power-law spectrum, and the eigenvalue scaling under the linear scaling of the total variance.

We start by assuming that the eigenspectrum is written in the form:

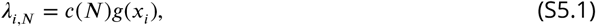

where *c*(*N*) is the scaling factor depending on the population size *N* and *g*(*x*_*i*_) is the scale-free structure of the eigenspectrum on the normalized index, *x*_*i*_ = *i*/*N*.

As discussed in text S2, subsampled populations satisfy linear total-variance scaling:

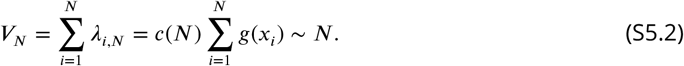

Using the integral approximation, the above summation is approximated as

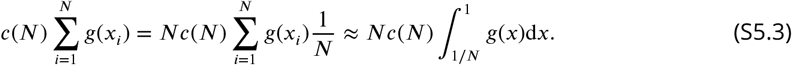

Note that the bounds of the integral are chosen over the domain of the function *g*(*x*), which is [1/*N*, 1]. We break the above integral into two parts:

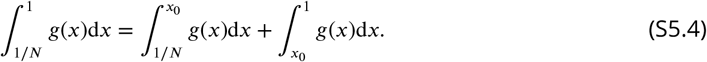

While the second integral on the right-hand side of the above equation is independent of *N*, the first integral depends on *N*.

Suppose that we have the power-law spectrum *g*(*x*) ~ *x*^−*μ*^ (*μ* > 0) at the limit of early indices of the eigenspectrum (*x* ≪ 1). Because the eigenspectrum decreases with rank, its slope is negative, implying *μ* > 0. If *μ* ≠ 1, the above integral is approximated as

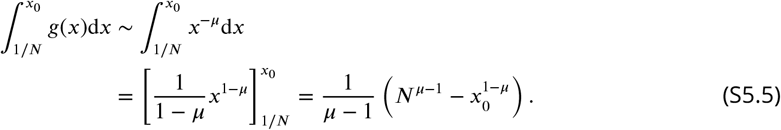

Further, if *μ* > 1, the above integral, Eq. S5.5, scales as *N*^*μ*−1^, resulting in

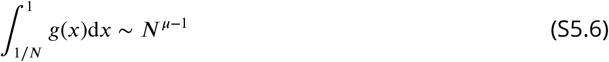

From Eqs. S5.3 and S5.6, the total variance scales as *V*_*N*_ ∝ *c*(*N*)*N*^*μ*^. The constraint of the linearly scaling total variance imposes *c*(*N*) = *N*^−*α*^ with *α* = *μ* − 1 > 0, resulting in the following scale invariance:

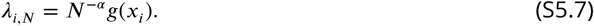

In this case, the eigenvalues at every fixed rank *i* scale linearly, *λ*_*i,N*_ ~ *N*^−*α*^(*i*/*N*)^−*μ*^ = *N*^*μ*−*α*^ ~ *N* with *μ* = 1 + *α*. These linearly scaling noise components can potentially limit the LFI.

By contrast, if 0 < *μ* < 1, the above integral, Eq. S5.5, converges to a positive constant. Then, the total variance scales as *V*_*N*_ ~ *Nc*(*N*), making *c*(*N*) constant under the linear-scaling total variance constraint. In this case, we have the following scale-invariant eigenspectrum:

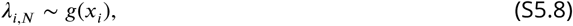

which can also be written in the form of Eq. S5.7 using *α* = 0. The eigenvalues at the limit of early indices scale sublinearly, *λ*_*i,N*_ ~ *N*^*μ*^ for a fixed *i*(≪ *N*). Since the eigenspectrum is constructed in descending order, the later eigenvalues cannot exceed the earlier eigenvalues, from which we conclude that all of the eigenvalues are sublinear functions of *N* if *α* = 0. Without the presence of the linear scaling eigenvalues, the LFI cannot be limited (text S1). This result implies that when neural populations show scale-invariant eigenspectra with *α* = 0 in their noise covariance, their LFI is unbounded as long as the signal magnitude 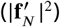 scales linearly with *N*.

Finally, the critical exponent *μ* = 1 results in *V*_*N*_ ~ *Nc*(*N*) log *N*, resulting in *c*(*N*) ~ (log *N*)^−1^. It is a decreasing function of *N* for *N* > 1. The eigenvalues at early indices scale as *λ*_*i,N*_ ~ *N*/ log *N*.

In summary, if we observe a scale-invariant eigenspectrum with an exponent *α* > 0 for large *N*, the exponent *μ* of the power-law scaling must be 1 + *α* in the limit of early indices, and the leading eigenvalues scale linearly. By contrast, if we observe a scale-invariant eigenspectrum with an exponent *α* = 0, the exponent of the power-law scaling *μ* must be less than one, and all eigenvalues scale sublinearly. We note that the same constraints are obtained by requiring the finite-*N* empirical eigenvalue mean to approach a finite, nonzero limit (text S6).

## text S6. Scale invariance and LFI in terms of eigenvalue distributions

In this note, we show that the relation between the scaling parameter *α* and the power-law exponent *μ* of the eigenvalue spectrum under the linear total variance is obtained by requiring the mean eigenvalue to have a finite, positive limit as *N* increases.

Let 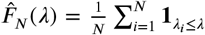 be an empirical cumulative distribution function (CDF) of the eigenvalues. Here, 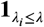 is an indicator function that takes one if the condition of the subscript is satisfied. Let *x*_*i*_ = *i*/*N* for *i* = 1, …, *N* be the normalized index of the eigenvalue *λ*_*i*_. Since we ordered the eigenvalues in descending order, it is related to the empirical CDF as

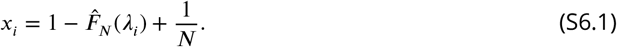

For simplicity, we assume that no two eigenvalues are exactly equal. We approximate 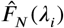 by a smooth CDF *F*_*N*_ (*λ*) for each finite *N*, where *λ* is a nonnegative real value. Then, the eigenvalue spectrum and the CDF *F*_*N*_ (*λ*) of the eigenvalues are related as follows.

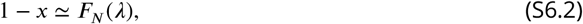

where we introduced *x* ∈ (0, 1]. Similarly, we approximate the empirical density of the eigenvalues 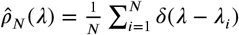 by a continuous density function 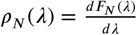.

Thus, under the scaling-invariant eigenspectrum satisfying

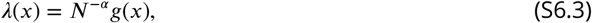

the eigenvalue density is related to *g*(*x*) via

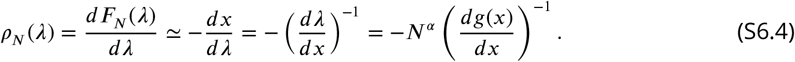

In the following, we assume the power-law for the eigenvalue spectrum at low indices: *g*(*x*) = *bx*^−*μ*^ for *x* ≪ 1. Then, if *μ* ≠ 1, the eigenvalue density follows the power-law with exponent 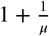 at *x* ≪ 1:

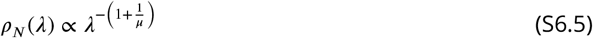

This is because, from *λ* = *N*^−*α*^*bx*^−*μ*^, we have

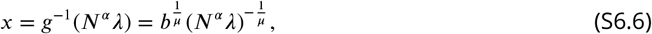

hence

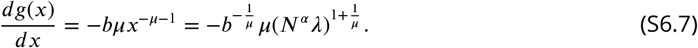

The constraint of the linear scaling of the total variance or the sum of eigenvalues translates into the condition that the mean eigenvalue approaches a finite positive value. More specifically, assuming that the smoothing error in the mean vanishes as *N* increases, we note that the mean of the sampled eigenvalues is related to the eigenvalue spectrum as follows:

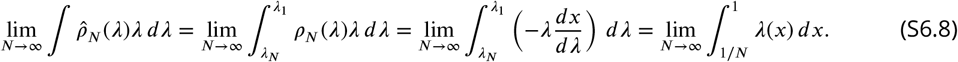

For *μ* ≠ 1 and *α* > 0, the right-hand side of the above equation is given as

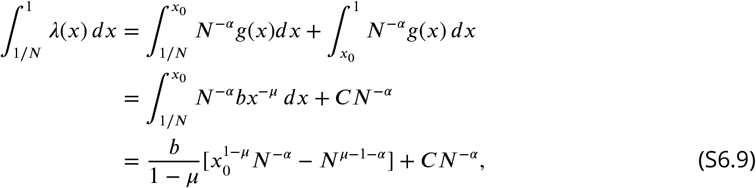

where *C* is a constant value. Note that *N*^−*α*^ → 0 in the large-*N* limit for *α* > 0. Thus, for the above equation to approach a finite, nonzero value, we need to impose *μ* − 1 − *α* = 0, or *μ* = 1 + *α*. In this case, *μ* > 1, which makes the integral positive and finite.

For *μ* ≠ 1 and *α* = 0, the right-hand side of the above equation is given as

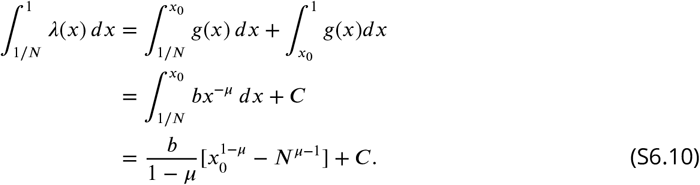

For the above equation to have a finite limit, *μ* must satisfy *μ* < 1 so that *N*^*μ*−1^ → 0 in the large-*N* limit. These results are consistent with the scaling relations obtained under the linear scaling of the total variance in text S5 (The case with *μ* = 1 is treated separately there).

Finally, we also note that the LFI can be computed using the finite-*N* distributions of *λ*_*i,N*_ and cos^2^ *θ*_*i,N*_ as follows, where *c* = cos^2^ *θ* and *p*_*N*_ (*λ, c*) is their joint distribution:

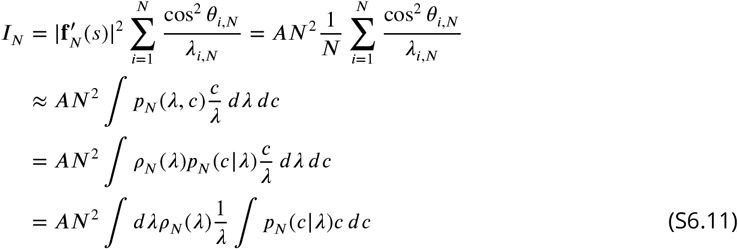

Here we note that the power-law relation between the eigenvalue and angular occupation (Eq. 7 or Eq. S3.1) specifies the conditional expectation of the angular occupation. Then, following the definition of Λ(*γ*) in text S3, we obtain

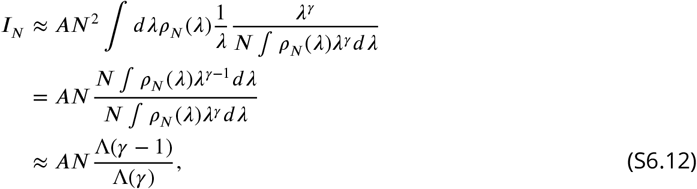

which recovers, within the continuum approximation, Eq. S3.17.

## text S7. Subspace analysis by partial least squares may incorrectly report saturated LFI

In this note, we show that analyzing LFI using a dimension-reduction method, known as the Partial Least Squares (PLS) discriminant analysis, can result in an incorrect conclusion about its boundedness, while our method correctly predicts its behavior in the large-*N* limit.

The PLS discriminant analysis identifies directions within a subspace of measurements that most effectively discriminate between two stimuli. The study by Rumyantsev et al. (*16*) applied the PLS analysis to distill the high-dimensional neural space into a latent space of five dimensions obtained from stimulus-informative PLS directions. They discovered that the noise eigenvector within this reduced space most aligned with the signal direction had an eigenvalue that scaled linearly with the population size across five examined mice, and concluded that these and other eigenvalues that increase with population size were responsible for the saturated behavior of the LFI, denoted as *d*^′^ in their analysis. However, the PLS directions are not generally principal eigenvectors of the full covariance, and the covariance eigenvalues within the PLS subspace need not be the leading eigenvalues of the full covariance. Thus, restricting the LFI to the PLS subspace may miss collective contributions from signal components distributed across many full-space noise eigenvectors, including those with sublinearly scaling eigenvalues. Consequently, the full-space LFI can be unbounded even when the reduced-space LFI appears bounded.

To test this hypothesis, we simulated neural populations exhibiting unbounded LFI theoretically. For this goal, for each population of size *N*, we constructed *N* eigenvalues from the eigenspectrum given by Eqs. 5 and 6 with the parameters derived from mouse TX42 whose estimated scaling parameter was *α* = 0.164±0.002. To define angular occupations, we employed the relation between the angular occupation and eigenvalues given by Eq. 7 with an exponent *γ* = 1, which results in unbounded LFI for any *α* > 0 (text S3).

More specifically, we constructed the covariance matrix **Σ**_*N*_ by associating random orthogonal vectors with the eigenvalues obtained from Eqs. 5 and 6. To construct **f**_*N*_ (*s*) and **f**_*N*_ (*s* + δ*s*), we assumed that the activity means are equal to the variance (i.e., the diagonal of **Σ**_*N*_). We then constructed 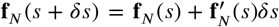, where 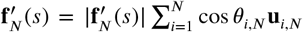 is constructed using the cos^2^ *θ*_*i,N*_ that comes from the power-law relation given by Eq. 7 using *γ* = 1. We then sampled neural activities from the Gaussian distribution *N*(**f**_*N*_ (*s*), **Σ**_*N*_) and *N* (**f**_*N*_ (*s* +δ*s*), **Σ**_*N*_). The sample covariance matrix and signal for a subpopulation of size *N* were constructed from 2600 samples (trials). The eigenspectra and angular occupations for subpopulation size *N* were obtained by averaging the results from 20 independently generated distributions. To compare the PLS and scaling-based analyses under the same finite-sample conditions, we used raw sample eigenvalues and angular occupations without bias correction.

The leading eigenvalues of the simulated dataset scale linearly as expected for *α* > 0 (fig. S9A). As expected, the data show both the collapse of the eigenvalue spectrum by the scaling and the power-law relation between eigenvalues and angular occupations (fig. S9B,C). The scaling parameter and the slope were estimated as *α* = 0.203 ± 0.0001 and *γ* = 1.009 ± 0.011. By applying the PLS method to the unbounded data, we find that the 10 eigenvalues extracted by the PLS method all scale linearly with population size (fig. S9D), and only partially recover the scaling behavior of the full system (fig. S9E,F).

Figure S9G shows the estimated LFI based on the PLS and our scaling-based method. The estimate of the LFI based on the PLS analysis (red) shows the saturated behavior, which follows the curve predicted by the differential correlations (dotted line). To the contrary, our scaling-based method (green) with estimated *α* and *γ* correctly predicts that the LFI of the population is unbounded (fig. S9H). We note that the large eigenvalues from finite samples are biased upwards, compared to the original values (*43*). While this results in the overestimation of *α*, using the estimated *α* is sufficient to conclude the unboundedness of the LFI because the critical *γ*_*c*_ is a decreasing function of *α*. For comparison, we provided the bias-corrected LFI estimate (*17, 25*), which estimates the LFI precisely. However, we note that the method cannot predict the behavior of LFI beyond the number set by the number of trials and neurons.

To further investigate the behavior of the PLS analysis, we simulated neural populations exhibiting unbounded LFI, assuming that the eigenvalues follow the scale invariance given by Eq. 5 with *α* = 0. We proved that the LFI of subsampled populations is unbounded under this condition (text S5). All the eigenvalues, including the leading eigenvalues, scale sublinearly. We then sampled eigenvalues from the same modified *α* = 0 eigenspectrum with the remaining parameters derived from mouse TX42. We set the leading-edge exponent to *μ* = 0.5 and the angular-occupation exponent to *γ* = 1. Thus, *μ* was chosen independently rather than constrained to equal 1 + *α* in this simulation.

As expected, the leading eigenvalues of the sampled data scaled sublinearly (fig. S10A). The eigenvalue spectrum in the normalized indices was consistent with *α* = 0 under the scaling-collapse fit (fig. S10B). The fitted exponent *γ* was *γ* = 0.968 ± 0.016 (fig. S10C), which is close to 1, but the error was larger than the previous example, possibly due to the lack of a large magnitude of *λ*_*i,N*_ because all eigenvalues scale sublinearly.

When we apply the PLS method to these data, we find that the largest eigenvalue scales closer to a linear line than the original eigenvalues (fig. S10D), despite the fact that there are no linearly scaling eigenvalues in the original dataset. Similarly to the previous example, the method did not fully recover the scaling behavior present in the full system. Accordingly, the estimate of the LFI based on the PLS analysis for these unbounded data shows again the saturated behavior (fig. S10G). Although our scaling-based analysis overestimated the magnitude of the LFI for *α* = 0, it correctly predicted its unbounded behavior (fig. S10H).

## text S8. LFI of the subsampled populations is a non-decreasing function of population size

In this note, we derive a recurrence relation for the LFI of subsampled populations. This relation clarifies how the LFI changes when a neuron is added to a subpopulation, demonstrating that the LFI of subsampled populations is a non-decreasing function of population size.

When we increase the number of neurons by adding a neuron to an existing subpopulation of size *N*, the covariance of the larger population of size *N* + 1, **Σ**_*N*+1_, contains the covariance of the population of size *N*, **Σ**_*N*_. Let Σ_*N*+1_ be the variance of the (*N* + 1)-th neuron and **c**_*N*_ be the covariance between the existing *N* neurons and the (*N* + 1)th neuron. The covariance matrix of the *N* + 1 neurons is written as

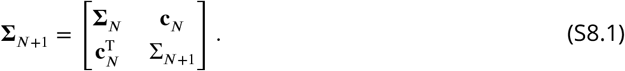

The determinant of the sub-matrix **Σ**_*N*_ is the leading principal minor of order *N* of the matrix **Σ**_*N*+1_. Further, the determinants of Σ_1_, **Σ**_2_, …, **Σ**_*N*_, **Σ**_*N*+1_ constitute the leading principal minors of **Σ**_*N*+1_. Sylvester’s criterion states that **Σ**_*N*+1_ is positive definite if and only if all of the leading principal minors are positive.

From the block determinant formula, we have

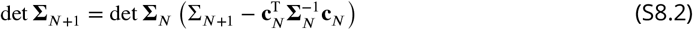

Given that **Σ**_*N*_ is positive definite, we have det **Σ**_*N*_ > 0. Then, **Σ**_*N*+1_ is positive definite if and only if

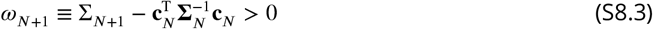

Starting from a single neuron with a positive variance Σ_1_, Sylvester’s criterion is satisfied if we construct the covariance matrix recursively while satisfying *ω*_*N*+1_ > 0. Indeed, this equation must be satisfied for a subsampled population of any size because the covariance of the subsampled population is positive definite.

We now consider the relation between the LFI of the population with size *N* and *N* + 1. First, we calculate the inverse of the covariance matrix. Using the following formula for the inversion of a block matrix

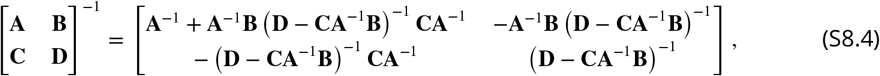

we obtain

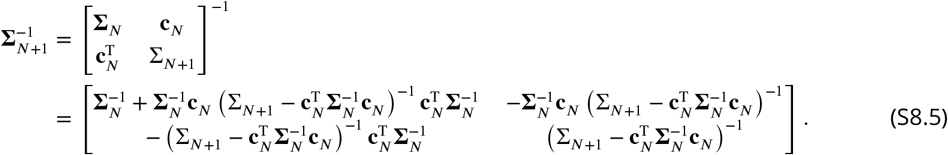

Using *ω*_*N*+1_ and

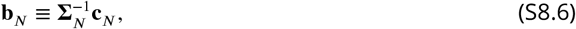

the inverse of the covariance matrix is written as

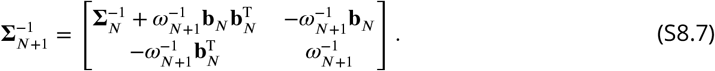

Similarly to the covariance matrix, we separate 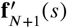 as

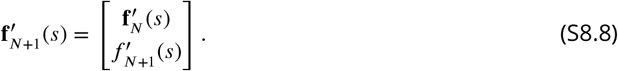

With this, the LFI is obtained as

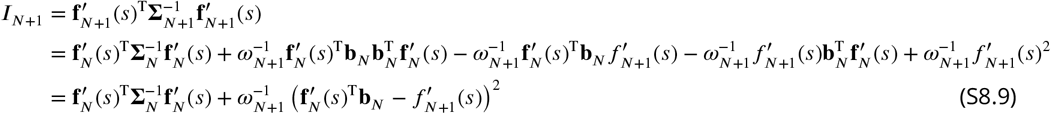

from which we have

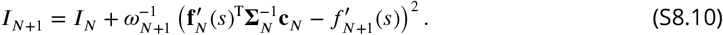

Since *ω*_*N*+1_ > 0, the second term of the above equation is nonnegative. Hence, we obtain

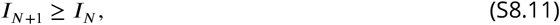

where the equality holds for 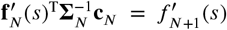. This proves that the LFI of the subsampled population is a non-decreasing function of *N*.

Eq. S8.10 provides insights into how the information increases or does not increase by adding a neuron. First, if **c**_*N*_ = **0**, namely, if an added neuron is independent of the other *N* neurons, the LFI increases by 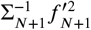. Further, if 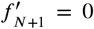, namely, even if we add a neuron that is not sensitive to a stimulus, the LFI increases by 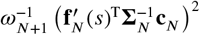. Finally, the LFI does not increase if 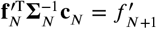.

Note that the LFI can be interpreted as an infinitesimal value of the squared standardized mean difference along the direction that discriminates the distributions of the neural activity responding to two close stimulus conditions *s* and *s* + *ds*. If we have two multivariate Gaussian distributions with means **f**_*N*_ (*s*) and **f**_*N*_ (*s* + *ds*) and the same covariance matrix **Σ**_*N*_, linear discriminant analysis shows that the direction 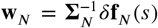, where δ**f**_*N*_ (*s*) = **f**_*N*_ (*s* + *ds*) − **f**_*N*_ (*s*), best discriminates the stimuli. One can project the data on this line by 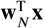, resulting in univariate distributions with 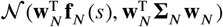 or 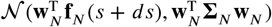, depending on the stimuli. The separability of these two distributions is quantified by the standardized mean difference,

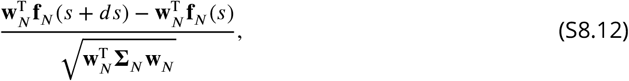

and its square is given as

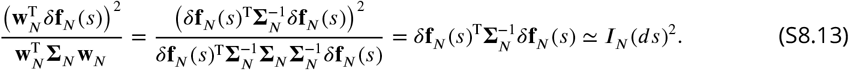

Thus, *I*_*N*_ = *I*_*N*+1_, namely 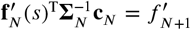, is the condition under which adding the neuron does not improve the optimal linear discriminability between the two stimuli.

## text S9. Noise of bounded subsampled populations contains differential correlations

In this note, we prove that the noise covariance of subsampled populations inducing saturating LFI and satisfying *I*_*N*_ < *I*_∞_ at every finite *N* can be decomposed into a covariance that induces unbounded LFI and differential correlations. This demonstrates that the boundedness of LFI under subsampling conditions is directly linked to the presence of differential correlations.

Given an (invertible) covariance matrix 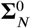 whose LFI 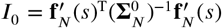 diverges as *N* goes to infinity, Moreno-Bote et al. (*22*) proved that the following specific form of correlations always limits the information:

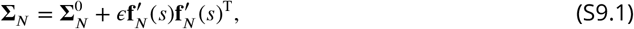

where 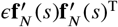 represents differential correlations, with *ϵ* being a (positive) constant. It can be confirmed as follows. Applying the Sherman–Morrison formula, the inverse of **Σ**_*N*_ is given as

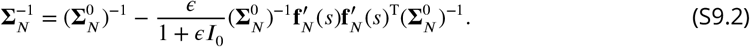

Consequently, the LFI 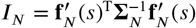 is given by

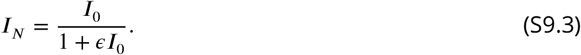

Given that *I*_0_(*N*) → ∞, the above equation is a saturating function that converges to 1/*ϵ* as *N* goes to infinity. Thus, the differential correlations limit the information.

It was further demonstrated that the differential correlations are special because adding an outer product of any other direction to 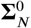 cannot limit the information. Therefore, they proposed that differential correlations are the only information-limiting correlations. The intuition behind this is that the noise structure 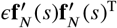 aligns with the direction of a signal change. No linear decoder can use such a structure to improve estimation accuracy. However, because the proof considers only rank-one additions to 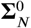, it does not exclude information limitation caused by higher-rank additions or by other covariance transformations, such as rotations of 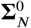.

Here, we remark that it is impossible to directly test for the presence of differential correlations by fitting Eq. S9.3 to empirical LFI from finite-size populations because *I*_0_ is an increasing, unbounded function whose form is unspecified a priori, and *ϵ* is unknown. Namely, because *I*_0_ is determined by the eigenvalues of 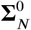 and the corresponding angular occupations (text S4), its scaling depends on the scaling of these quantities. Consequently, Eq. S9.3 can describe any increasing and saturating function of *N* that remains below the unknown limiting value 1/*ϵ*. Thus, fitting this equation to the empirical LFI cannot, by itself, prove or disprove the presence of differential correlations.

We now examine whether the covariance matrix of subsampled populations resulting in saturation of the LFI contains differential correlations. We start with the assumptions that the covariance matrix **Σ**_*N*_ is positive definite for any finite *N* and that 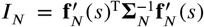 saturates as *N* → ∞: *I*_*N*_ → *I*_∞_, where *I*_∞_ is a finite value. To test for the presence of differential correlations, we ask whether

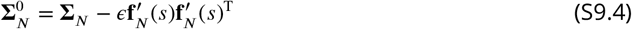

is a positive definite (and therefore invertible) covariance matrix. If 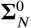 is positive definite, the Sherman–Morrison formula gives

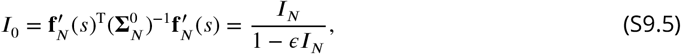

or equivalently,

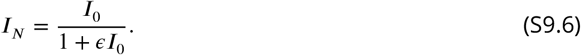

This is precisely the form derived for differential correlations (Eq. S9.3). Choosing *ϵ* = 1/*I*_∞_ yields the limiting value *I*_∞_ = 1/*ϵ*. Moreover, *I*_0_ diverges as *I*_*N*_ → *I*_∞_ = 1/*ϵ*, showing that the first term in the covariance decomposition (Eq. S9.1) indeed has unbounded LFI. Thus, if 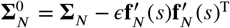 is positive definite for *ϵ* = 1/*I*_∞_, then **Σ**_*N*_ admits a decomposition into an unbounded covariance matrix and a differential-correlation term.

We first note the necessity of the strict inequality *I*_*N*_ < *I*_∞_ for the positive definiteness of 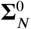. If 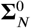 is positive definite, then *I*_0_ given by Eq. S9.5 must be finite and nonnegative, which requires 1 − *ϵI*_*N*_ > 0. With *ϵ* = 1/*I*_∞_, this condition is equivalent to *I*_*N*_ < *I*_∞_. The monotonicity result for subsampled populations gives *I*_*N*+1_ ≥ *I*_*N*_ (text S8); hence a saturating subsampled LFI satisfies *I*_*N*_ ≤ *I*_∞_ for every finite *N*. The equality case is degenerate: if *I*_*N*_ = *I*_∞_ for some finite *N*, monotonicity implies *I*_*k*_ = *I*_∞_ for all *k* ≥ *N*, and 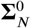 is singular because 1 − *ϵI*_*N*_ = 0. We therefore restrict our analysis to the nondegenerate saturating case in which *I*_*N*_ < *I*_∞_ for every finite *N*.

The remaining task is to prove the converse direction, namely that this strict inequality is sufficient for positive definiteness of 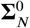. We prove this below by exploiting the nested structure of subsampled covariance matrices.

Given a positive definite covariance matrix **Σ**_*N*+1_, we ask if

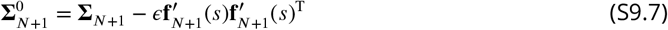

is positive definite or not. If so, **Σ**_*N*+1_ can be decomposed into a covariance matrix and differential correlations. Since we consider an incremental increase in the population size from *N* to *N* + 1, 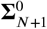 can be written using Eq. S8.1 as

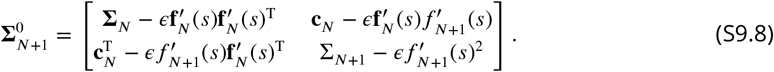

Note that the upper left block matrix is 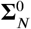 (Eq. S9.4). Hence, by applying the block determinant formula, the determinant of 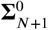 is given by

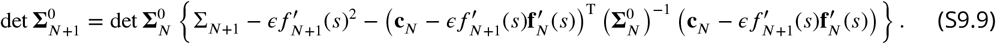

From this equation, the ratio between the determinant of 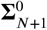 and 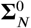 is given as

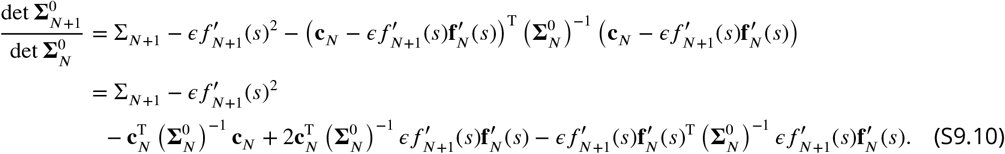

Using 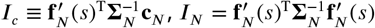, and the Sherman–Morrison formula for,

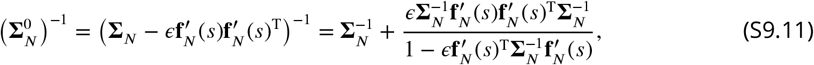

the above equation becomes

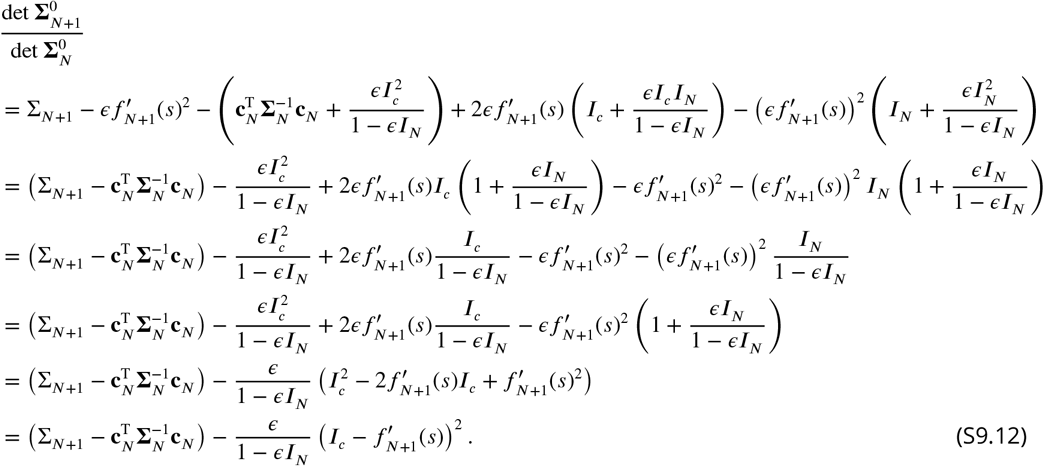

Using Eq. S8.10, the equation above becomes

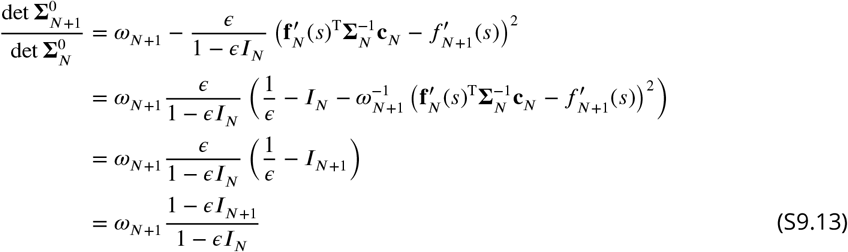

This equation can also be obtained by directly applying the matrix determinant lemma, det 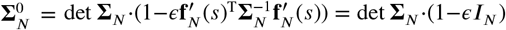, to both det 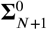 and det 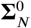, together with det **Σ**_*N*+1_/ det **Σ**_*N*_ = *ω*_*N*+1_ (Eq. S8.3).

Note that *ω*_*N*+1_ > 0 from Eq. S8.3. Furthermore, with *ϵ* = 1/*I*_∞_, the strict bounds *I*_*N*_ < *I*_∞_ and *I*_*N*+1_ < *I*_∞_ imply that both the numerator and denominator below are positive:

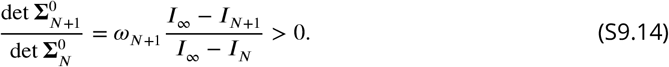

Therefore, given det 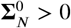, we obtain det 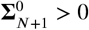. A single neuron (*N* = 1) with a positive variance Σ_1_ satisfies det 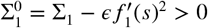 since 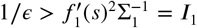. Thus, det 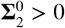 for the population of two neurons. Then, according to Sylvester’s criterion, 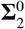 is positive definite. Recursively applying this relation to increasing the number of neurons up to *N* + 1 guarantees that all of the leading principal minors of 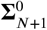 are positive, proving that 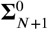 is positive definite for any *N*.

## text S10. Differential correlations induce linearly scaling eigenvalues

In this note, we show that if 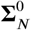 leads to the unbounded LFI for all directions of 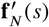, by adding differential correlations, only one of the eigenvectors of Σ_*N*_ will be substantially aligned with 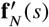 and only the eigenvalue associated with this eigenvector will linearly increase with *N*. Otherwise, if *I*_0_ is unbounded only for specific signal directions and 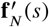 retains a nonvanishing component outside the linear-eigenvalue subspace of 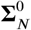, there are multiple linear eigenvalues in Σ_*N*_ whose eigenvectors asymptotically span 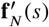. These results show that alignment of the signal with the linear-eigenvalue subspace is necessary to limit the information.

If differential correlations exist, for any subsample of neurons, 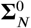 must exist and because it is a covariance matrix, it can be written in terms of its eigenvalues *γ*_*i,N*_ and eigenvectors **v**_*i,N*_ as

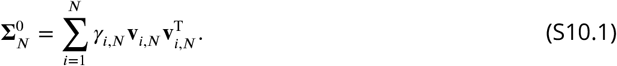

Since the set of orthonormal eigenvectors is complete, we can write 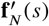 as a linear combination of the eigenvectors of 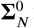:

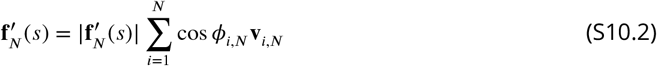

with ∑_*i*_ cos^2^ *ϕ*_*i,N*_ = 1, in which *ϕ*_*i,N*_ is the angle between the *i*-th eigenvector of 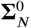 and 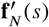. We can also write

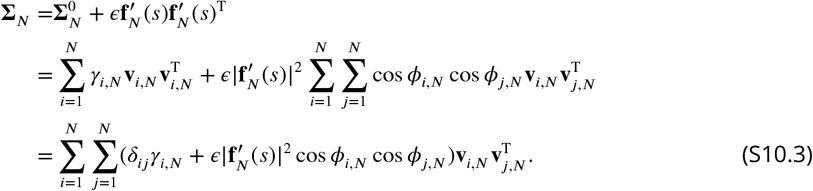

We multiply the above equation from the right side by 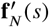

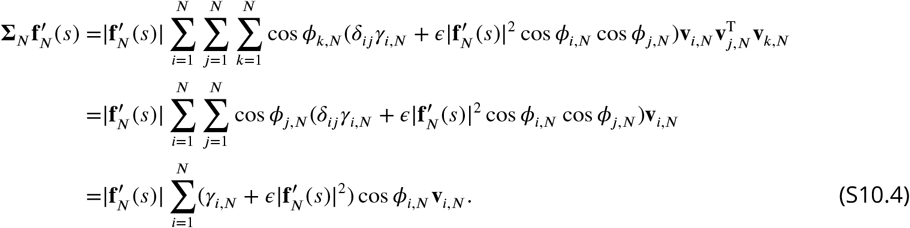

Furthermore, if 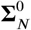 is independent of 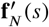 and gives unbounded LFI for every signal direction with 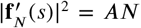, then all *γ*_*i,N*_ must be sublinear functions of *N*. Indeed, choosing 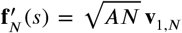 gives *I*_0_(*N*) = *AN*/*γ*_1,*N*_ → ∞, so *γ*_1,*N*_ /*N* → 0; by eigenvalue ordering, all *γ*_*i,N*_ are sublinear.

Therefore,

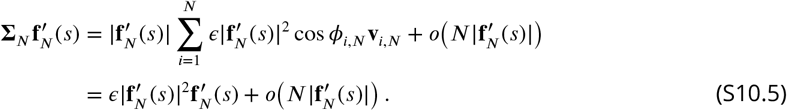

Here and below, *o*(*a*_*N*_) denotes a term whose magnitude divided by *a*_*N*_ tends to zero as *N* → ∞. Because the largest eigenvalue of 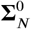 is *o*(*N*), adding 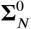 changes the eigenvalues of the rank-one differential-correlation term by at most *o*(*N*). Therefore, 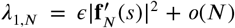 and *λ*_*i,N*_ = *o*(*N*) for *i* ≥ 2. This result states that, in this limit, 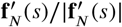 asymptotically aligns with the leading eigenvector of Σ_*N*_. In other words, 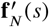 is spanned by only this specific eigenvector of Σ_*N*_. Namely, if 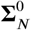 is independent of 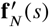 and gives unbounded LFI for every signal direction, we expect that only one of the eigenvectors becomes increasingly aligned with 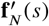 as *N* increases. Specifically, one of the direction cosines (*i* = 1) dominates the spectrum of cosines (i.e., lim_*N*→∞_ cos^2^ *θ*_1,*N*_ = 1) and the rest of the cosines go to zero as *N* increases. In other words, in the large-*N* limit, we have

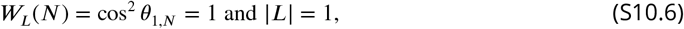

in which |*L*| is the number of elements (cardinality) of the set *L*.

On the other hand, if 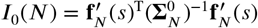 is unbounded only for specific signal directions, not all *γ*_*i,N*_ are sublinear functions. If 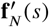 retains a nonvanishing component outside the linear-eigenvalue subspace of 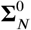, the differential-correlation term produces an additional linearly scaling eigenvalue. Consequently, the linear-eigenvalue subspace of Σ_*N*_ is multidimensional and asymptotically contains 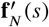. A particularly interesting case is Σ_*N*_ exhibiting infinitely many linear eigenvalues for which the LFI boundedness is equivalent to restricting the orientation of the signal vector to a particular subspace, e.g., *γ* > *γ*_*c*_ if 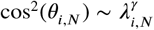.

## text S11. Noise of populations expressing finite limiting LFI may not contain differential correlations

In this note, we demonstrate that the differential correlations are not the unique form of information-limiting correlations for covariance matrices even if they are subject to the linearly increasing total variance. A specific decomposition of the covariance matrix Σ_*N*_ is shown below, which resembles the structure of differential correlations (having the sum of two matrices, one that limits information and the other that does not limit), but has a potentially higher-rank information-limiting matrix added to the non-information-limiting component. We then give a formal disproof.

Given that the linear and sublinear eigenvalues of Σ_*N*_ are respectively written as *λ*_*i,N*_ = *b*_*i*_*N* + *F*_*i*_(*N*) (*i* ∈ *L*) and *λ*_*i,N*_ = *G*_*i*_(*N*) (*i* ∈ *S*_*b*_) (See text S2), Σ_*N*_ can in general be decomposed as

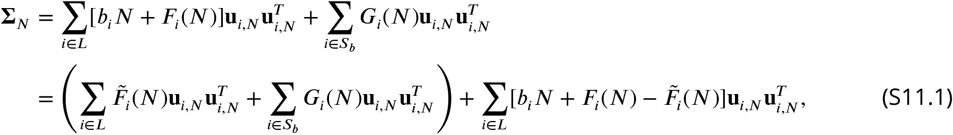

where 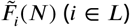 are sublinear functions satisfying the conditions below. Hence, we obtain the following expression for Σ_*N*_ :

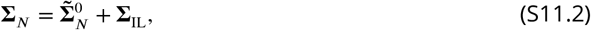

where the first term is

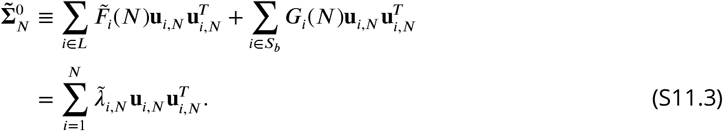

Here we introduced 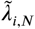 defined as 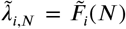 if *i* ∈ *L* and 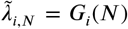 if *i* ∈ *S*_*b*_. Note that we can choose 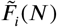 to be sublinear and satisfy 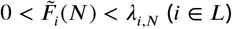. Then, 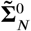 is a positive definite matrix. While 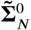 has the same set of eigenvectors as in Σ_*N*_, all of its eigenvalues are sublinear functions of *N*. Consequently, 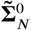 results in unbounded LFI for any 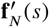 (text S1). The second term is

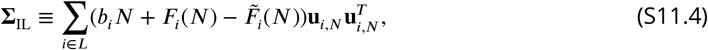

which is a rank |*L*| matrix and is positive semidefinite because 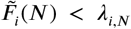 for all *i* ∈ *L*. This decomposition is applicable to any covariance matrix, including the case that 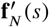 and Σ_*N*_ result in the saturation of information. In this case, the covariance matrix Σ_*N*_ was expressed as the sum of two matrices, where 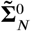 leads to unbounded LFI irrespective of the orientation of 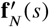, and a rank |*L*| matrix that is responsible for limiting the information. Note that this decomposition is not unique: We can construct various decompositions using sublinear functions 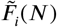 satisfying 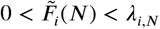.

An example of decomposing a more specific covariance matrix disproves the presence of differential correlation in a covariance matrix subject to the linearly increasing total variance. Suppose that the *k*th eigenvector **u**_*k,N*_ (*k* ∈ *L*) of Σ_*N*_ perfectly aligns with the direction of 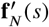. Such a covariance matrix results in the bounded information because *I*_*N*_ = *AN*/[*b*_*k*_*N* +*F*_*k*_(*N*)] → *A*/*b*_*k*_. Here, we note that, since 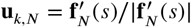, we have

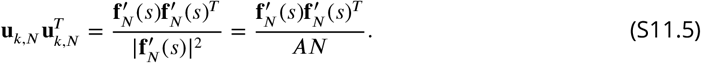

Using this relation, the covariance matrix is decomposed as

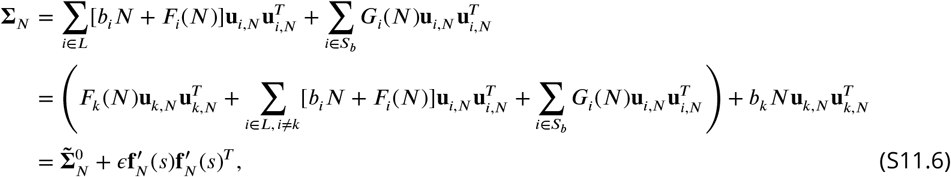

where we defined *ϵ* ≡ *b*_*k*_/*A* and

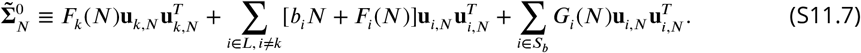

If *F*_*k*_(*N*) > 0, this matrix is positive definite. Unlike the previous decomposition, we did not exclude the linear eigenvalues except for *k* from 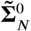. This is because **u**_*i,N*_ (*i* ∈ *L, i* ≠ *k*) are orthogonal to 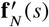; therefore, 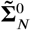 still results in unbounded LFI because *I*_0_(*N*) = *AN*/*F*_*k*_(*N*) → ∞ (See also text S10). Thus, such a covariance Σ_*N*_ is written as a sum of a non-saturating (invertible) covariance matrix and the differential correlations. In other words, Σ_*N*_ contains differential correlations.

However, *F*_*k*_(*N*) can be nonpositive. The eigenvalues of Σ_*N*_, including the linear eigenvalues *λ*_*i,N*_ = *b*_*i*_*N* +*F*_*i*_(*N*) (*i* ∈ *L*), must be positive so that Σ_*N*_ is positive definite. Hence, the sublinear functions *F*_*i*_(*N*) in the linear eigenvalues must satisfy *F*_*i*_(*N*) > −*b*_*i*_*N*, which allows *F*_*i*_(*N*) to be either zero or negative. If 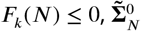 is no longer a positive definite matrix. If *F*_*k*_(*N*) = 0, i.e., *λ*_*k,N*_ = *b*_*k*_*N*, the rank of 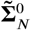 becomes *N* − 1, and 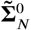 is not invertible. With *ϵ* = 1/*I*_∞_ = *b*_*k*_/*A*, the bounded covariance Σ_*N*_ having these properties cannot be decomposed into a sum of an (invertible) covariance matrix with unbounded LFI and the differential correlations. This example disproves the statement that differential correlations are the only information-limiting correlations.

In what follows, we give a formal disproof of the differential correlations using the example above, following an approach that proved the presence of differential correlations for the sub-sampled populations with saturating LFI (text S9). We assume that we have Σ_*N*_ that results in the bounded information with its limit being given by lim_*N*→∞_ *I*_*N*_ = 1/*ϵ*, namely *ϵ* = 1/ lim_*N*→∞_ *I*_*N*_. The presence of the differential correlations is guaranteed if we can decompose Σ_*N*_ in the form of Eq. S9.1, where 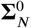 is a positive definite covariance matrix.

Using eigenvectors of Σ_*N*_ and 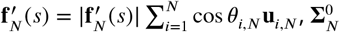 is written as

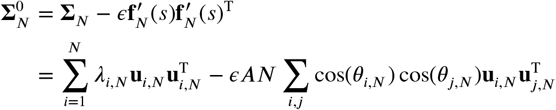

A positive definite matrix Σ_*N*_ can be diagonalized by an orthogonal matrix as 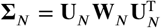, where **W**_*N*_ is a diagonal matrix with its entries being the eigenvalues of Σ_*N*_. The *i*th element of the *j*th column **w**_*j,N*_ of **W**_*N*_ is given by *w*_*ij,N*_ = *λ*_*i,N*_ *δ*_*ij*_. Then from

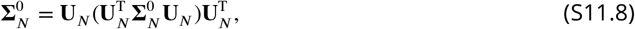

we investigate 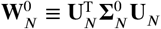 because if 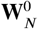 is a positive definite matrix, so is 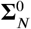. Noting **U**_*N*_ **W**_*N*_ = Σ_*N*_ **U**_*N*_ or **U**_*N*_ [**w**_1,*N*_, …, **w**_*N,N*_] = Σ_*N*_ [**u**_1,*N*_, …, **u**_*N,N*_], we have **U**_*N*_ **w**_*i,N*_ = Σ_*N*_ **u**_*i,N*_, from which we obtain

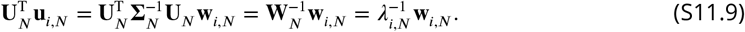

Hence,

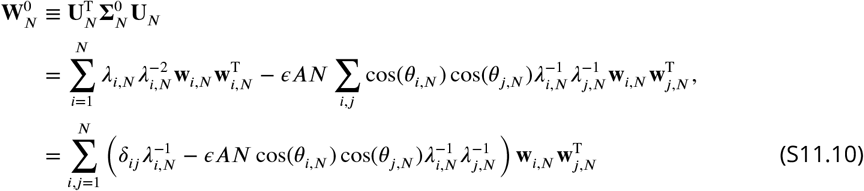

The (*k, l*) element of 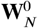 is computed as

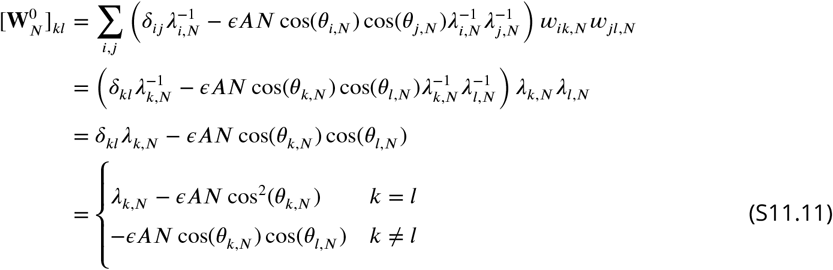

Hence, for 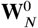 to be a positive definite matrix, it must satisfy

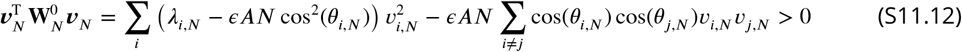

for any nonzero real vector ***v***_*N*_ (***v***_*N*_ ≠ 0) when *ϵ* = 1/*I*_∞_.

We examine this condition with the simple case discussed above, in which one linear eigenvalue occupies a cosine direction to 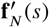. For simplicity, we consider that the first linear eigenvalue occupies the cosine direction (cos^2^ *θ*_1,*N*_ = 1). In this case,

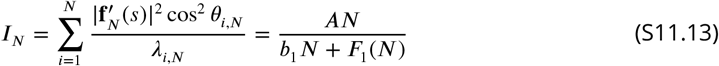

from which we obtain *I*_∞_ = *A*/*b*_1_. Hence we use *ϵ* = *b*_1_/*A*.

The quadratic term then becomes

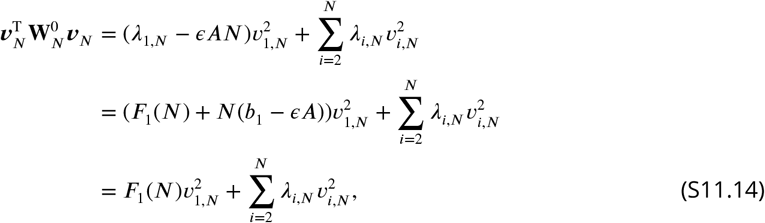

If 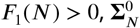 is positive definite. However, if 0 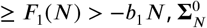 is not positive definite. When 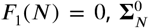 is not invertible, which also violates the condition for the differential correlations. We note that when *F*_1_(*N*) = *λ*_1,*N*_ − *ϵAN* < 0, we have

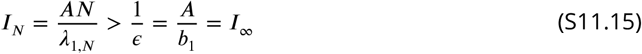

So the LFI approaches *I*_∞_ from above.

With this example, we corroborated that if a linear eigenvalue of the covariance matrix occupies the angular occupation, and the sublinear component of the linear eigenvalue is not positive, the covariance matrix does not contain the differential correlations with a positive-definite residual at *ϵ* = 1/*I*_∞_.

## text S12. Bias correction for covariance eigenvalue estimation

It is well known from the random matrix theory that the estimated covariance eigenvalues with finite sample size tend to push each other away on the real positive axis, so that the domain of the estimated eigenvalues is larger than the domain of the actual eigenvalues, leading to bias in their estimates, i.e., underestimation of small eigenvalues, and overestimation of large ones. Here, we present a method for correcting this bias. Throughout this note, we consider a fixed population size *N*. For notational simplicity, we therefore suppress the explicit *N*-dependence in quantities such as *λ*_*i,N*_ and write them as *λ*_*i*_.

Let *r*_*i,t*_(*s*_*j*_) denote the response of neuron *i* on trial *t* to stimulus *s*_*j*_. The population response vector for a subpopulation of size *N* is **r**_*t*_(*s*_*j*_) = [*r*_1,*t*_(*s*_*j*_), *r*_2,*t*_(*s*_*j*_), …, *r*_*N,t*_(*s*_*j*_)]^T^, its average across trials is ***μ***(*s*_*j*_), and the direct estimate of the covariance matrix is

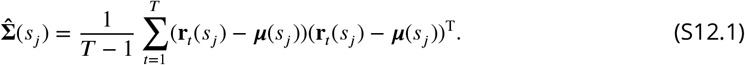

Here, we assume that this estimate is a sample of a Wishart distribution (*17*),

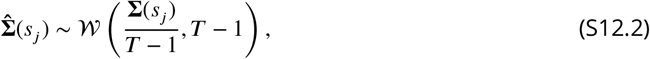

where Σ(*s*_*j*_) is the actual covariance matrix. In this paper, we have two sets of stimuli (*s*_1_, *s*_2_), and we use the estimate of the covariance as the average of the two covariance matrices, i.e., 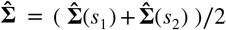. If the two stimulus groups have the same true covariance, Σ(*s*_1_) = Σ(*s*_2_) = Σ, then

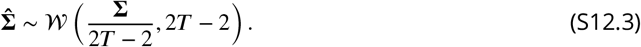

Eigenvalues of 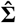 are the naive and biased estimates of the eigenvalues of Σ. Let 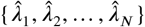 be the ranked eigenvalues of 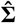 and {*λ*_1_, *λ*_2_, …, *λ*_*N*_} be those of Σ. Because the Wishart distribution is closed under orthogonal unitary transformations, we can consider the diagonalization of Σ as

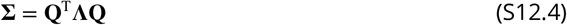

in which **Λ** is a diagonal matrix with {*λ*_1_, *λ*_2_, …, *λ*_*N*_} as its diagonal elements. To do the eigenvalue bias corrections, we work in the coordinate system in which Σ is diagonalized (principal component basis). In this coordinate system, we have

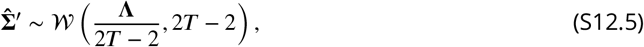

where 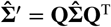. Note that the eigenvalues of 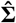 and 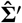 are the same.

Our bias correction approach is based on Eq. S12.5. Our goal is to find the set of ranked eigen-values {*λ*_1_, *λ*_2_, …, *λ*_*N*_} so that samples of the distribution exhibit the same eigenvalue spectrum as the naive estimates 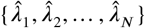. This goal was achieved by utilizing an iterative shrinkage algorithm. We start with an initial guess for **Λ**. The algorithm takes the following steps

1. Initial guess for bias-corrected eigenvalues is the naive estimates 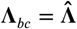
2. Generate *m* samples 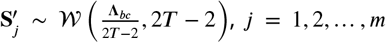, from the Wishart distribution in Eq. S12.5.
3. Evaluate the average of ranked eigenvalues of **S**^′^ across all *m* samples as 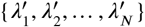.
4. Update the diagonal elements of **Λ**_*bc*_ by

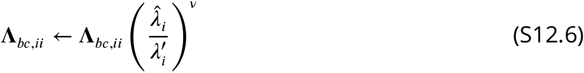

We repeat steps 2–4 until convergence. Here, *m* is equal to the number of population subsamples used in the naive estimation of the eigenvalues. This algorithm converges when the eigenvalues of **S**^′^ become equal to those of 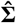. Note that if 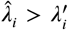 the bias-corrected eigenvalues get updated to larger numbers and vice versa if 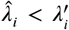. The exponent *ν* can be seen as a learning rate in the log scale, which is selected to be small enough to avoid large steps. In the end we find a set of bias-corrected eigenvalues ({*λ*_*bc*,1_, *λ*_*bc*,2_, …, *λ*_*bc,N*_}) for which the sampled eigenvalues 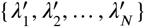 are equivalent to the naive estimates 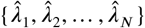 so that the ratio

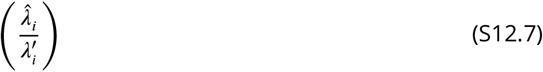

becomes equal to one for all components.

## text S13. Bias correction for estimating angular occupations

In this note, we provide a bias-correction algorithm for estimating angular occupations. As in text S12, we consider a fixed population size *N* and therefore suppress the explicit *N*-dependence of quantities for notational simplicity.

The error in estimating the angles *θ*_*i*_ between the signal vector *δ***f** and the eigenvectors of the covariance matrix **v**_*i*_ leads to biased estimates of the angular occupations cos^2^ *θ*_*i*_. This bias arises from two sources. First, as illustrated in the figure below, the estimated eigenvector 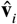 is oriented differently from the true eigenvector **v**_*i*_. Second, the estimated vector *δ****μ***, used as an approximation of *δ***f**, has a different orientation from the true vector.

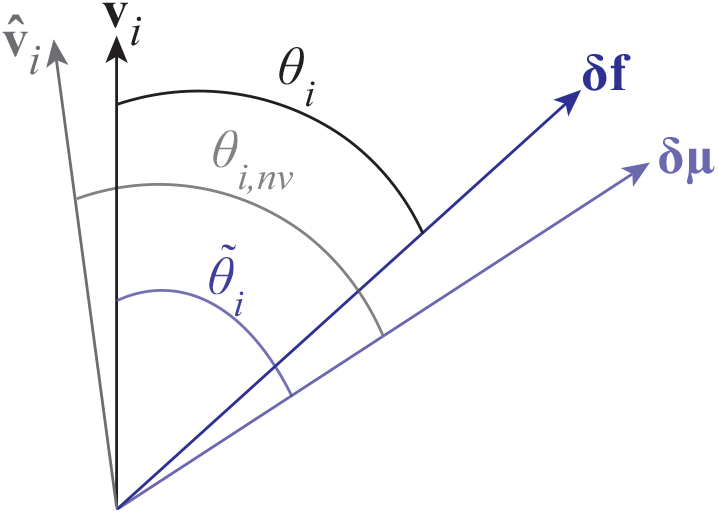

As shown in the above figure, we denote *θ*_*i,nv*_, the naive estimate of *θ*_*i*_, as the angle between *δ****μ*** and 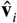. Similarly, we denote the angle between *δ****μ*** and the actual eigenvector **v**_*i*_ as 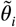. To correct the bias in angular occupations, we use a two-step method. First, we make the correction resulting from the error in the estimate of **v**_*i*_. In other words, the goal in the first step is to find 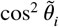. In the second step, we take 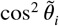 evaluated in the first step and correct for the bias caused by the error in estimation of *δ***f**.

### Angular occupation bias correction

To perform this correction, we adopt a recursive procedure analogous to the eigenvalue bias correction. Specifically, we seek the underlying true squared cosine values 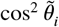 that would produce the observed naive estimates 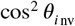 under the alignment implied by the Wishart samples of covariance eigenvectors. To do this, we work in the principal basis coordinates that give a diagonal representation of the true covariance matrix Σ. In this co-ordinate system, the normalized vector 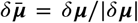 is represented by the vector of directional cosines that can have either positive or negative elements. Because we are focusing on cosine squares, the signs are irrelevant here, and we assume all cosines are positive. The algorithm proceeds as follows:

1. Initial guess for bias-corrected angular occupations is the naive estimates 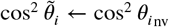
2. Generate *m* samples from the Wishart 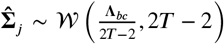 with *j* = 1, 2, …, *m*, where **Λ** is the diagonal matrix of bias-corrected eigenvalues obtained from the eigenvalue bias-correction algorithm.
3. Evaluate the eigenvectors of every sampled covariance matrix 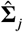 as 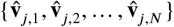 and find the sampled cosine squares as the squared dot product of these eigenvectors and 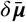 as 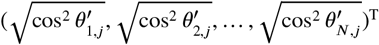.
4. Evaluate the average of 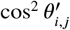 ranked according to eigenvalues of 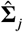 across all *m* samples as 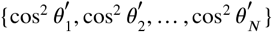.
5. Update the angular occupations by

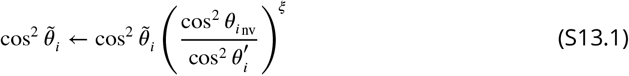
6. Normalize 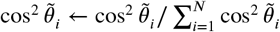.

We repeat steps 2–6 until convergence. Similar to the eigenvalue bias-correction algorithm, *m* is equal to the number of population subsamples used in the naive estimation of the cosine squares. This algorithm converges when 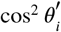 becomes equal to the naive estimate 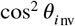 for all *i*. In the end we find a set of bias-corrected angular occupations 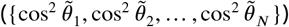 for which the sampled eigenvectors on average align with the naive estimates so that the ratio

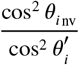

approaches one for all components. Similar to the eigenvalue bias-correction algorithm, the *ξ* exponent can be interpreted as a learning rate in log scale.

### Bias correction of signal estimation

Following Kafashan et al. (*17*), the estimated signal difference follows

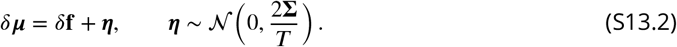

We assume that the covariance matrix Σ is known, and let *λ*_*i*_ and **v**_*i*_ be the eigenvalues and eigenvectors of Σ (i.e., Σ**v**_*i*_ = *λ*_*i*_**v**_*i*_). The naive estimate of the angular occupation is

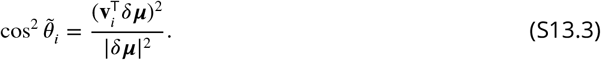

The noiseless cosine occupation is defined as

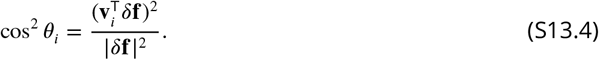

A perturbative expansion in *T* ^−1^ yields

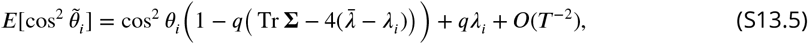

for *i* = 1, …, *N*, with

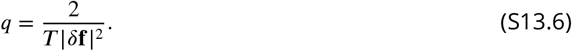

Throughout this note, big-*O* terms are bounded in magnitude by a constant multiple of the indicated order in the relevant limit. Here, 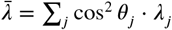 is the weighted average of eigenvalues by the angular occupations. We provide the derivation below. From the equation above, neglecting terms of order *O*(*T* ^−2^) and replacing the expectation by the naive estimate, we obtain the bias-correction formula:

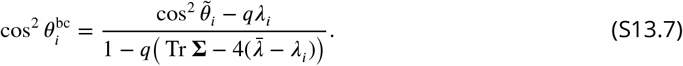

However, these are nonlinear equations with respect to cos^2^ *θ*_*i*_ because 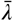 is a function of a set of 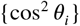. Here, in practice, we obtain 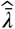 by the plug-in estimate

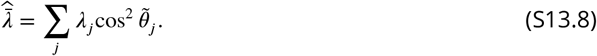

We apply a few iterations of the plug-in estimate by updating the angular occupation by the bias-corrected estimate. During this iteration, we clip the estimate at zero whenever the bias-correction returns a negative value. In the isotropic case, this bias-correction formula is equivalent, up to *O*(*T* ^−1^), to the one obtained under the factorization assumption described below (see the derivation at the end of this note). The bias correction with the term 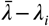 explicitly captures the direction-dependent contribution of the noise variance: in the general anisotropic case it removes eigenmode-specific bias, while in the isotropic limit it reduces to a uniform rescaling determined by Tr Σ.

Focusing on the contribution of the 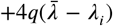 alone, which enters additively in the denominator, we note that it induces a differential rescaling of the estimated angular occupations. In particular, eigenmodes with 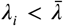 increase the denominator and therefore suppress the bias-corrected angular occupation, whereas modes with 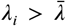 decrease the denominator and correspondingly increase the bias-corrected angular occupation.

*Proof of the perturbative expansion*

We assume that the noise ***η*** = *δ****μ*** − *δ***f** follows *N* (0, 2Σ/*T*). Its projection onto **v**_*i*_ yields

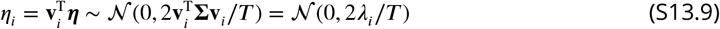

From the definition of eigenvalues, the noise variance along **v**_*i*_ is proportional to *λ*_*i*_. It also scales as *T* ^−1^. A perturbative expansion in *T* ^−1^ follows from expanding the ratio in powers of the small noise ***η***, whose covariance scales as Cov(***η***) = *O*(*T* ^−1^).

Let 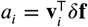 and note that 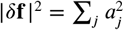. Then 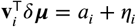, from which we obtain

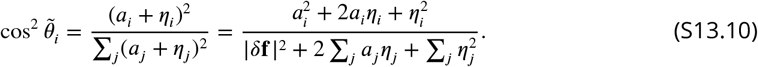

We expand this ratio through *O*(‖***η***‖^2^) as follows. Define

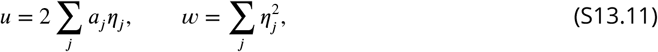

so that the denominator is |*δ***f** |^2^ + *u* + *w*, where *u* = *O*(‖***η***‖) and *w* = *O*(‖***η***‖^2^). Then

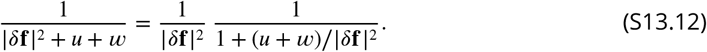

Using (1 + *z*)^−1^ = 1 − *z* + *z*^2^ + *O*(*z*^3^) with *z* = (*u* + *w*)/ |*δ***f**| ^2^, we get

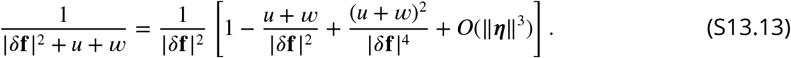

Since *u* = *O*(‖***η***‖) and *w* = *O*(‖***η***‖^2^), we have

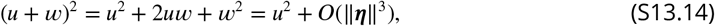

hence, up to *O*(‖***η***‖^2^),

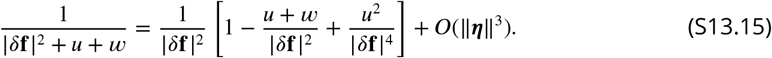

Now multiply by the numerator 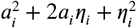:

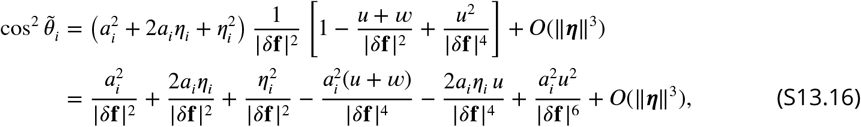

where we used that (2*a*_*i*_*η*_*i*_)*w* = *O*(‖***η***‖^3^), 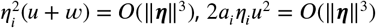, 2*a*_*i*_*η*_*i*_*u*^2^ = *O*(‖***η***‖^3^), and 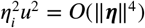.

We now take expectations term by term. Since the noise vector ***η*** is mean zero, all terms linear in ***η*** vanish upon averaging. In particular,

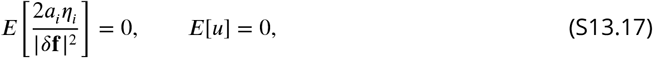

where *u* = 2 ∑_*j*_ *a*_*j*_*η*_*j*_.

The first nonvanishing contribution arises from the quadratic noise term in the numerator. Using 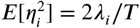, we obtain

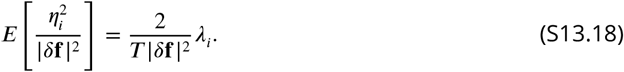

Next, consider the contribution from the expansion of the denominator proportional to 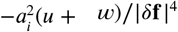, where 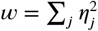. Since *E*[*u*] = 0, only the *w* term survives, giving

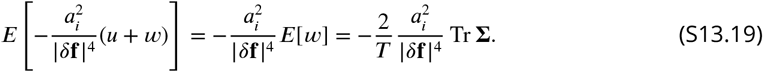

Next, the mixed quadratic terms give nonzero contributions. Since

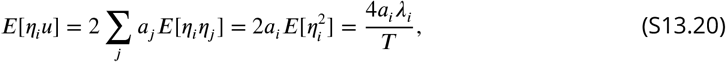

we have

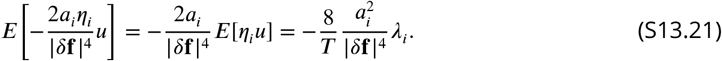

Moreover,

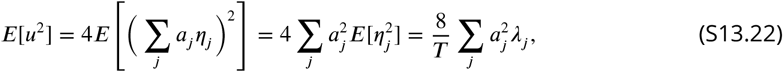

so

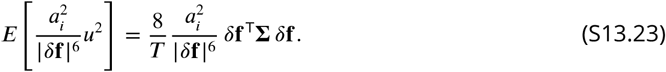

Collecting all terms, we obtain

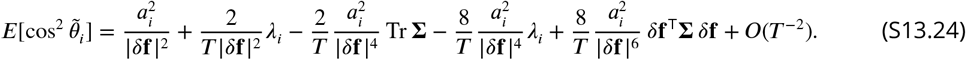

We note that the noiseless angular occupation is

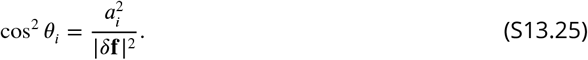

Using

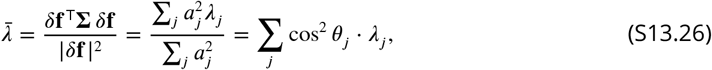

the expansion is written compactly as

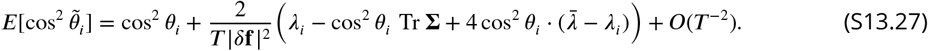

We define

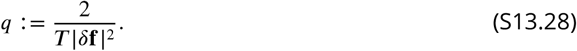

Neglecting terms of order *O*(*T* ^−2^), the expectation relation may be rewritten as

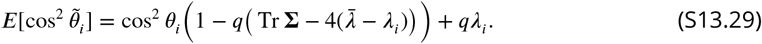

The equation above can be rewritten as

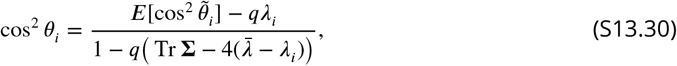

for *i* = 1, …, *N*. However, we note that these form nonlinear equations with respect to cos^2^ *θ*_*i*_ because 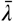 is a function of a set of {cos^2^ *θ*_*i*_}.

Replacing the unknown expectation by the observed value 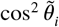 gives the bias-corrected plug-in estimator

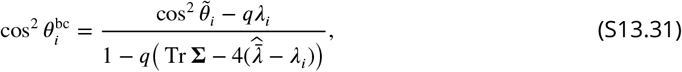

Where 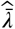 is obtained by the plug-in estimate

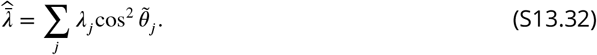

In practice, we apply a few iterations of the plug-in estimate by updating the angular occupation by the bias-corrected estimate.

If **f** |*δ*^2^| is unknown, a consistent estimator at order *T* ^−1^ is

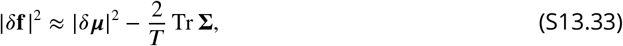

which yields

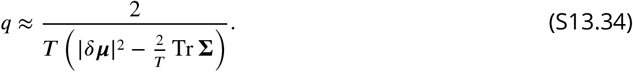

Finally, if the noise is isotropic (*λ*_*i*_ = *λ*) and there is no preferred directionality in the covariance eigenbasis, then we expect *E*[cos^2^ *θ*_*i*_] = 1/*N*, and we have 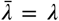. The bias correction equation is approximated by

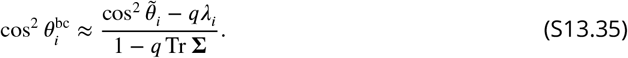

Since *q* = *O*(*T* ^−1^), we expand the denominator to first order:

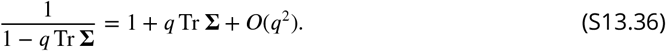

Substituting this expansion yields

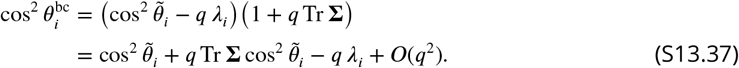

This isotropic expansion agrees, up to *O*(*T* ^−1^), with the factorization-based correction derived below. Recall that 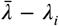 captures the direction-dependent noise contribution and vanishes in the isotropic limit, leaving a uniform correction.

### Bias-corrected angular occupations under factorization assumption

For comparison, we derive the simplified bias correction under the factorization assumption. Following Kafashan et al. (*17*), if the unbiased estimate of *δ****μ*** follows a normal distribution,

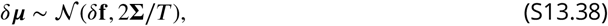

we have

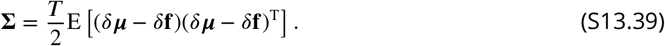

Multiplying on the left by an eigenvector transpose 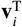 of Σ and on the right by **v**_*i*_, we get

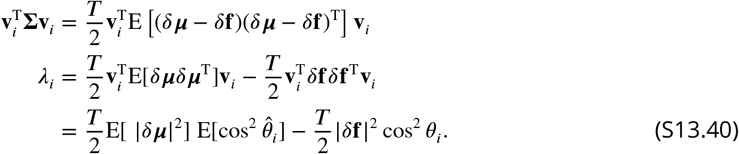

Here, 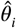 denotes the random angle between the fixed true eigenvector **v**_*i*_ and the noisy estimate *δ****μ***. At the last equality, we assumed the factorization

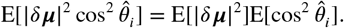

Summing over *i*, we get

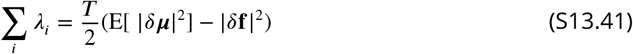

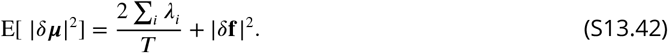

Substituting Eq. S13.42 into Eq. S13.40 and replacing 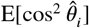 with the observed naive estimate 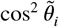, we find the bias-corrected estimate of the cosine squares as

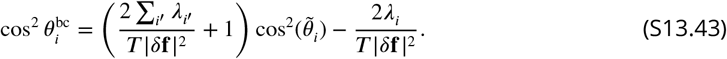

In the isotropic case, the factorization holds through first order in *q*, and the resulting correction agrees with the general perturbative correction in Eq. S13.37 to that order.

---

**Fig. S1.**
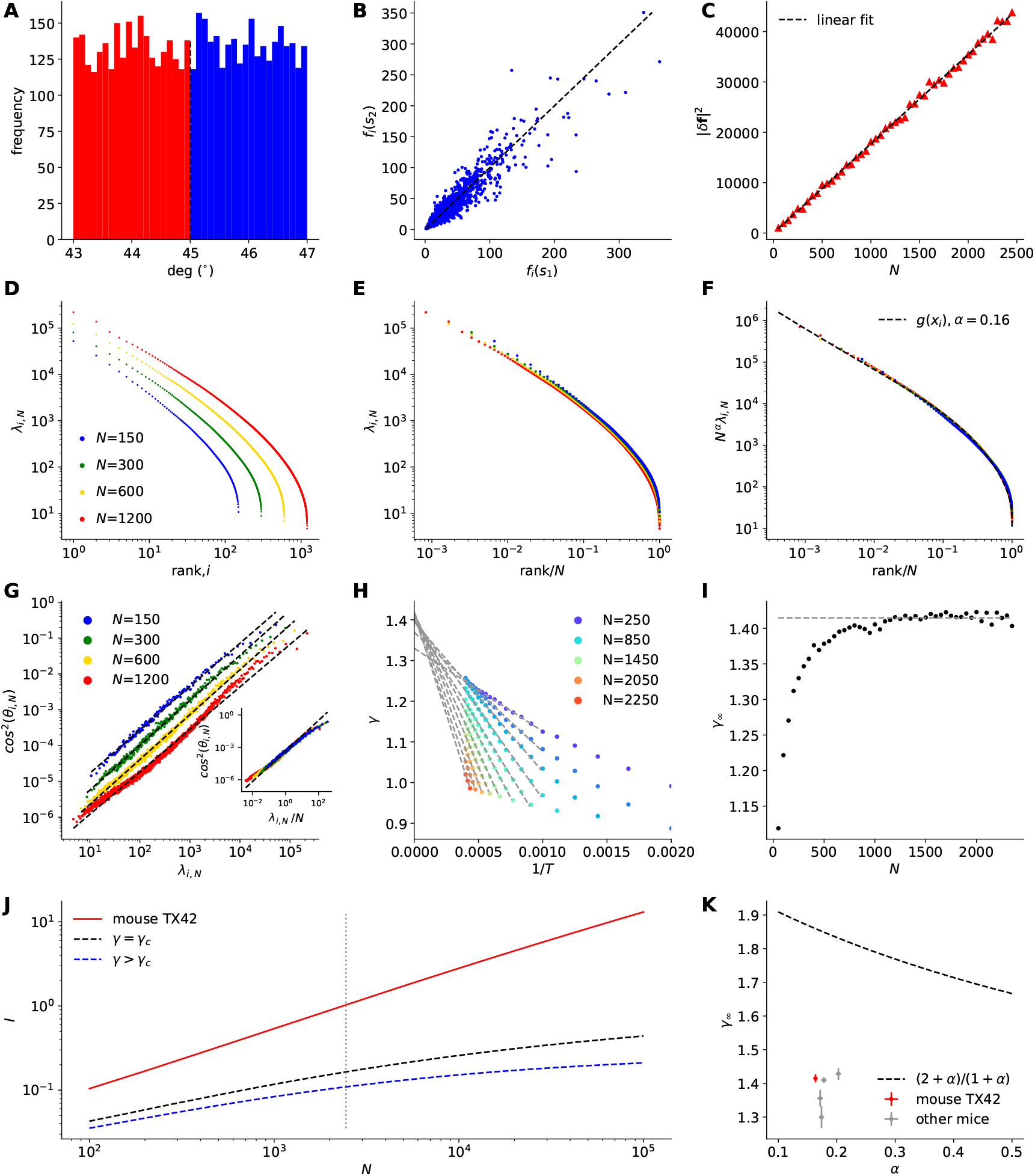
Analysis of mouse TX42. The figure styles follow Fig. 2 in the main text. The fitted scaling parameter is *α* = 0.164 ± 0.002. The fitted slope is *γ*_∞_ = 1.415 ± 0.006. **A**, A frequency histogram of grating stimulus orientations. **B**, Comparison of mean responses of neurons *f*_*i*_(*s*) to Stimuli 1 and 2. **C**, |*δ***f**_*N*_|^2^ vs the subpopulation size *N*. **D**, Noise eigenspectra for different *N*. **E**, The eigenspectra vs the normalized rank, (*x*_*i*_ =)*i*/*N*. **F**, The re-scaled eigenspectra by multiplying *N*^*α*^. The dashed line is the fitted *g*(*x*_*i*_). **G**, cos^2^ *θ*_*i,N*_ vs *λ*_*i,N*_ for different *N*. The dashed lines are linear fits. (Inset) cos^2^ *θ*_*i,N*_ vs the scaled eigenvalue, *λ*_*i,N*_ /*N*. **H**, The fitted *γ* as a function of 1/*T* for different *N*. The solid lines are fitted linear functions. **I**, The intercept *γ*_∞_ for different *N*. The average *γ*_∞_ (dashed line) was 1.415 ± 0.006. **J**, The inferred LFI (solid line) as a function of *N* in the log-log plot. The dashed lines show theoretical LFI. **K**, The estimated *γ*_∞_ vs *α*. The dashed line is a critical boundary.

**Fig. S2.**
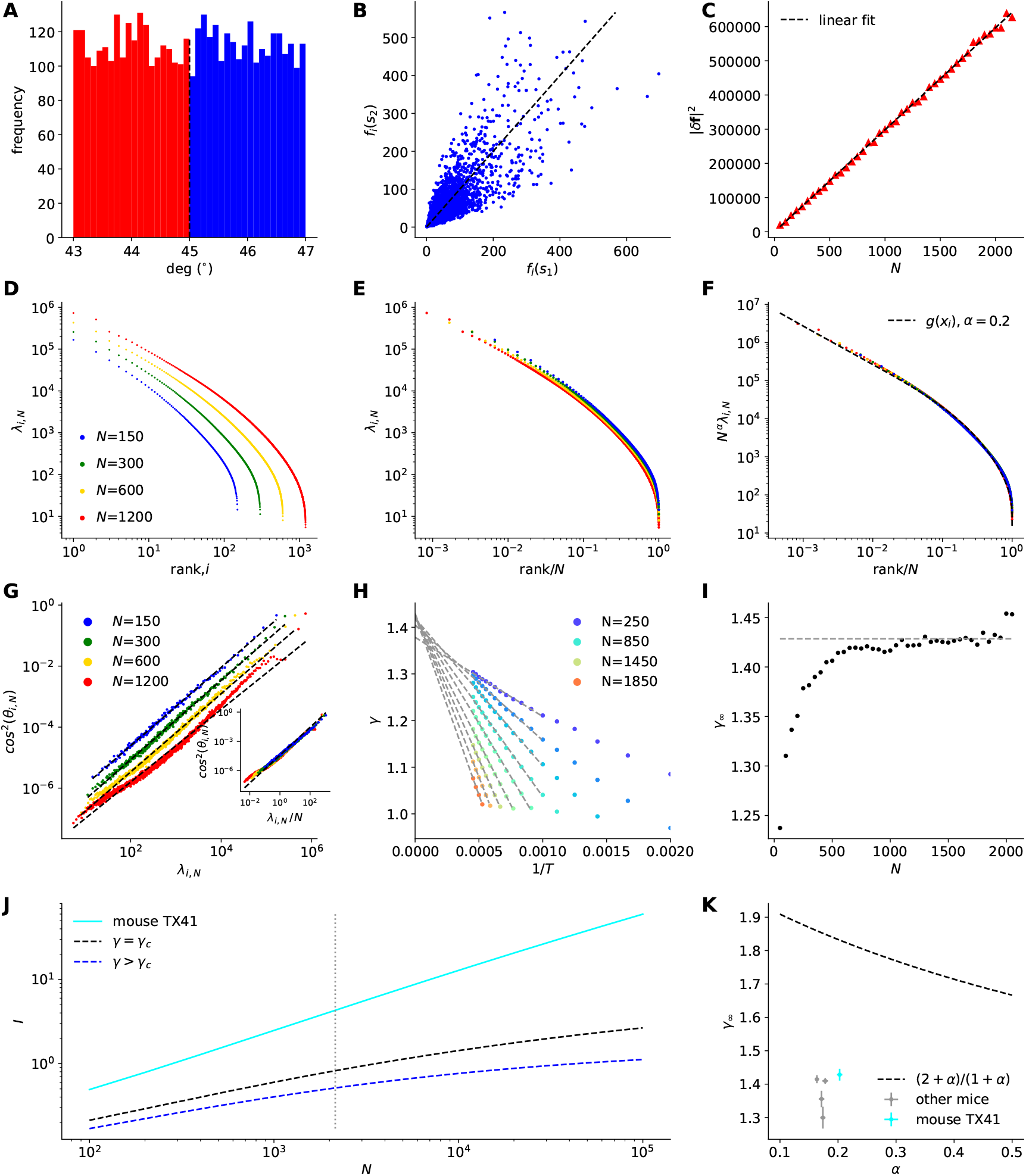
Analysis of mouse TX41. Same panels (A–K) as fig. S1. The fitted scaling parameter is *α* = 0.203 ± 0.002. The fitted slope is *γ*_∞_ = 1.428 ± 0.009.

**Fig. S3.**
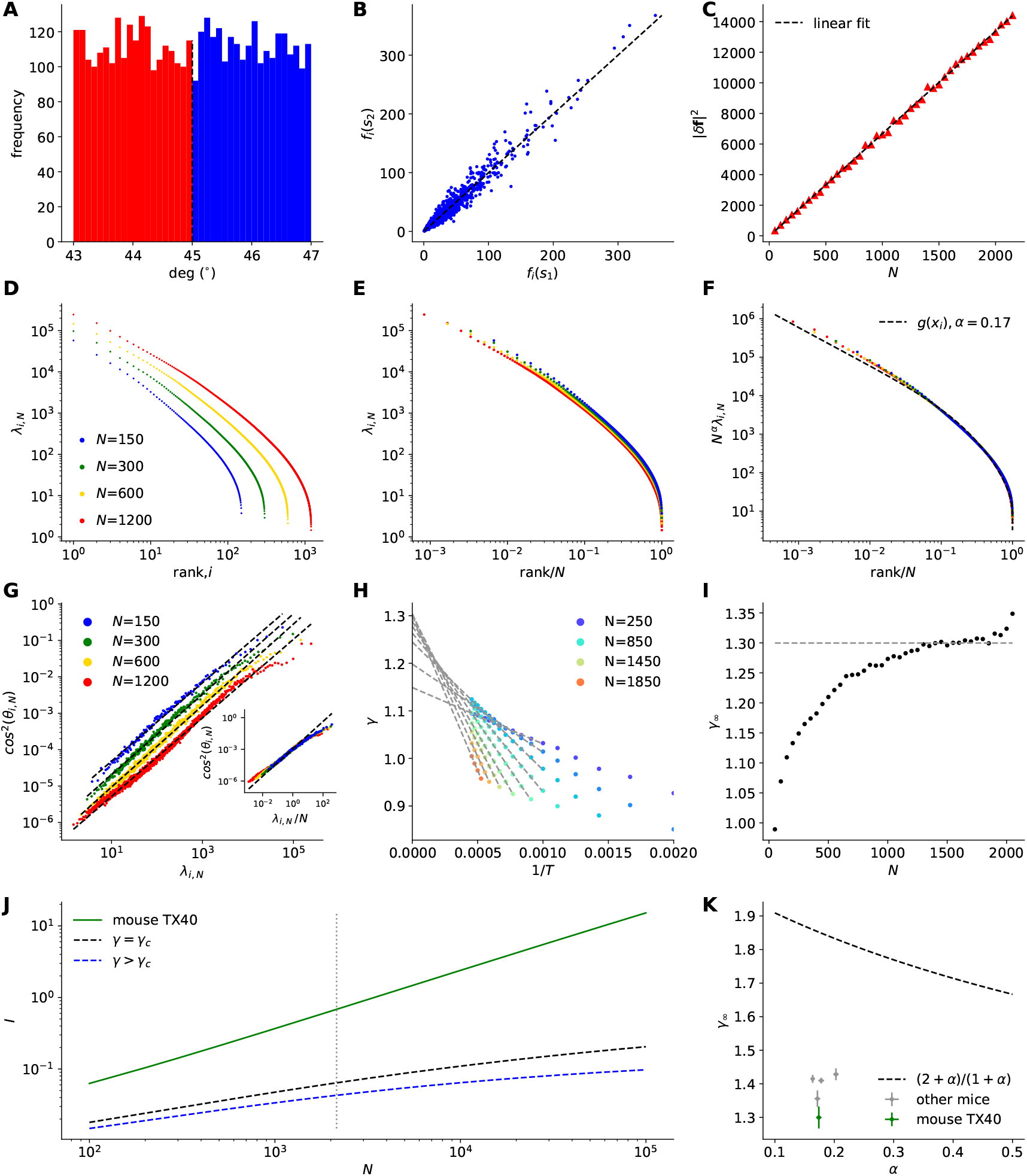
Analysis of mouse TX40. Same panels (A–K) as fig. S1. The fitted scaling parameter is *α* = 0.174 ± 0.002. The fitted slope is *γ*_∞_ = 1.3 ± 0.016.

**Fig. S4.**
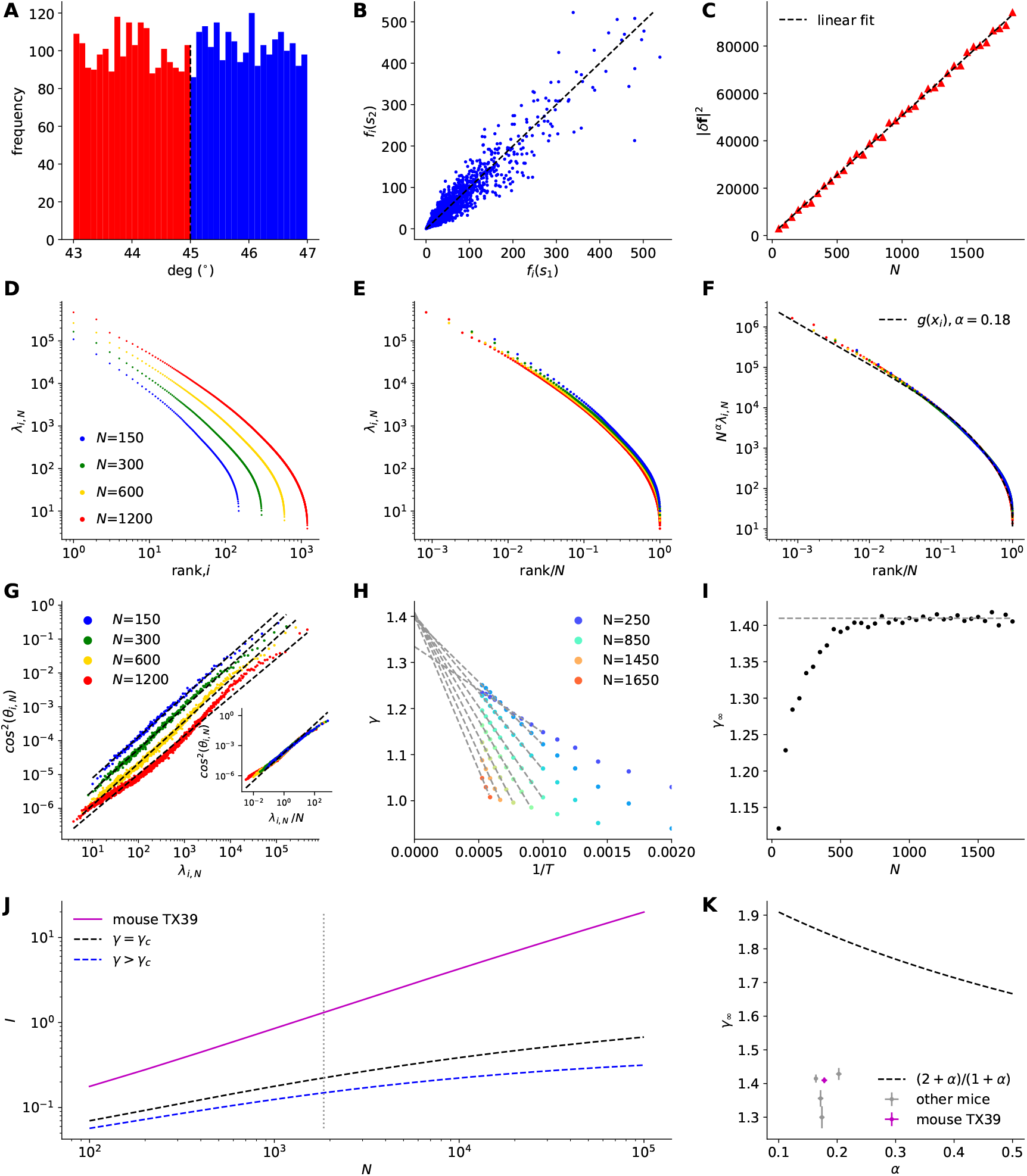
Analysis of mouse TX39. Same panels (A–K) as fig. S1. The fitted scaling parameter is *α* = 0.178 ± 0.002. The fitted slope is *γ*_∞_ = 1.41 ± 0.004.

**Fig. S5.**
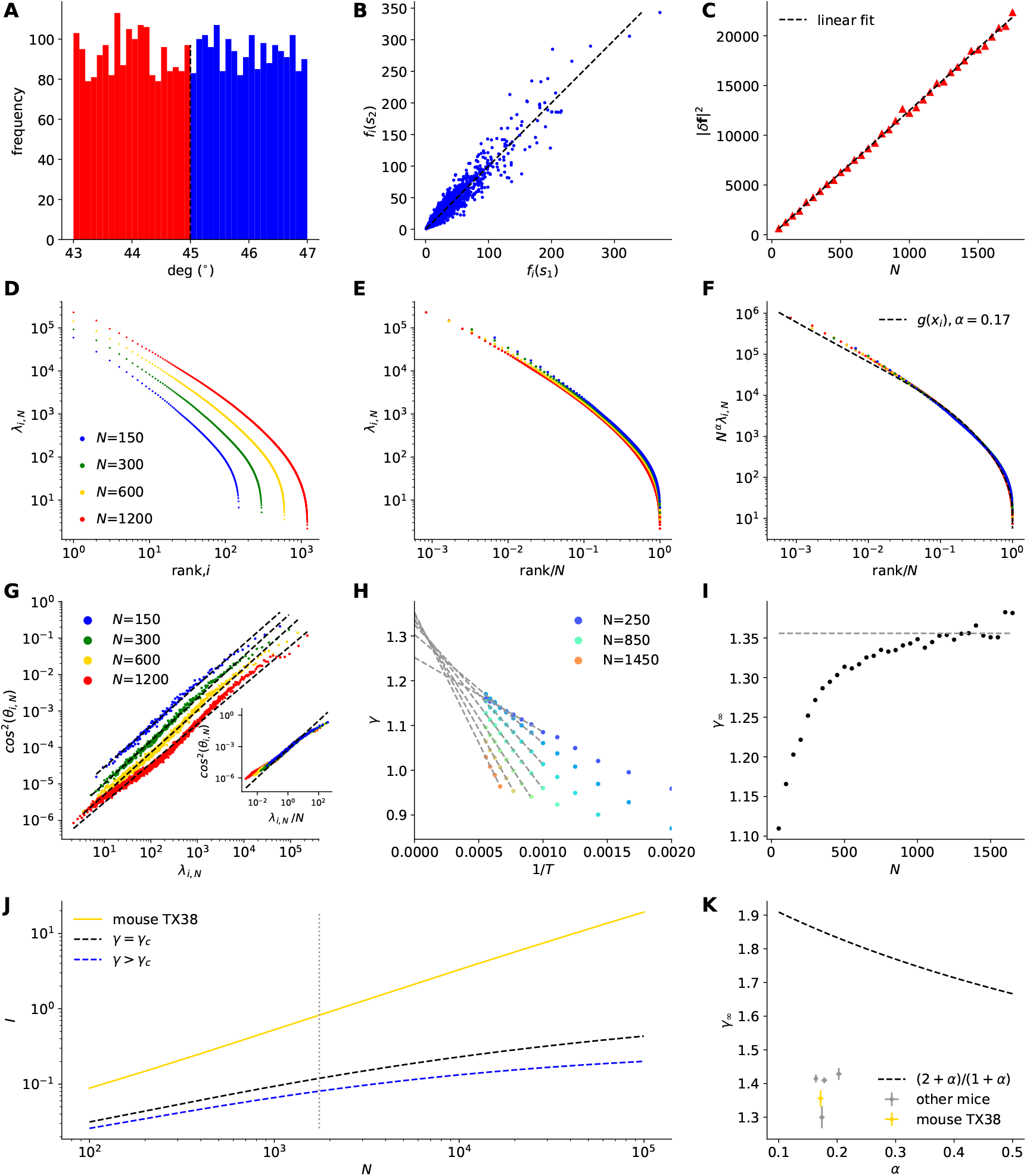
Analysis of mouse TX38. Same panels (A–K) as fig. S1. The fitted scaling parameter is *α* = 0.172 ± 0.002. The fitted slope is *γ*_∞_ = 1.356 ± 0.012.

**Fig. S6.**
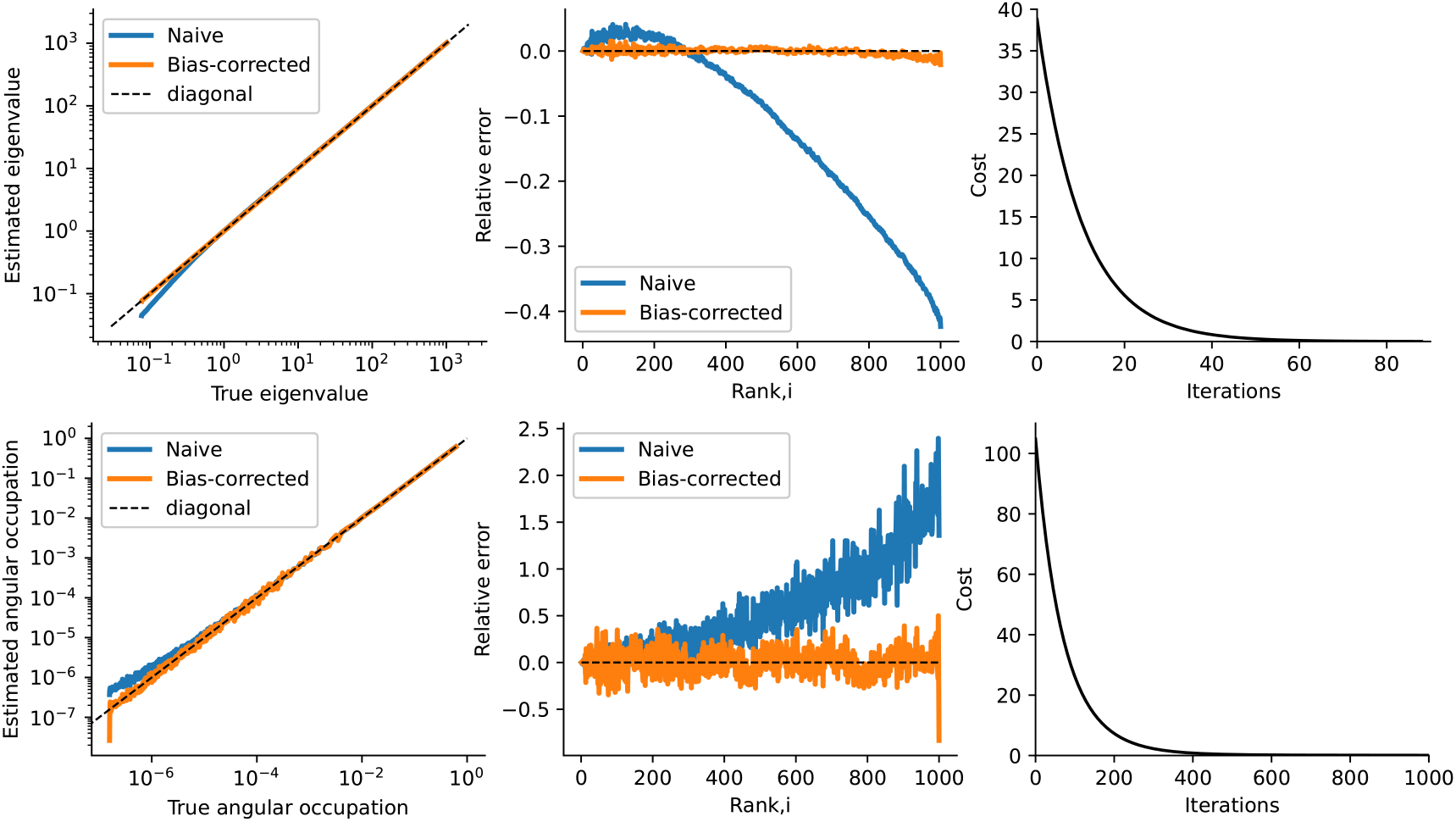
Validation of bias correction methods on simulated data. **A**, Covariance eigenvalue bias corrections. Left: Estimated versus true eigenvalues plotted on a log-log scale. The bias-corrected estimates fall on the diagonal, while the naive estimates deviate from it. Middle: To better visualize the error of the naive and bias-corrected estimates, the relative errors are shown versus the true eigenvalue ranks (lower ranks correspond to larger eigenvalues, *λ*_1_ > *λ*_2_ > ⋯ > *λ*_*N*_). Clearly, the naive estimates overstate the larger eigenvalues (yielding positive relative error) and understate the smaller eigenvalues (yielding negative relative error). The bias-corrected estimates show small errors. Right: The cost function of the algorithm 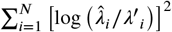 decreases to zero over iterations. **B**, The same as in **A**, but for the angular occupation corrections. The cost function plotted in the right panel is 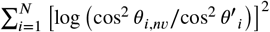 for the angular occupation bias-correction algorithm. In this simulation, the subsampled population size is *N* = 1000, the number of subsampled populations is *m* = 200, the number of trials is *T* = 2000, *α* = 0.25, and *γ* = 1.6.

**Fig. S7.**
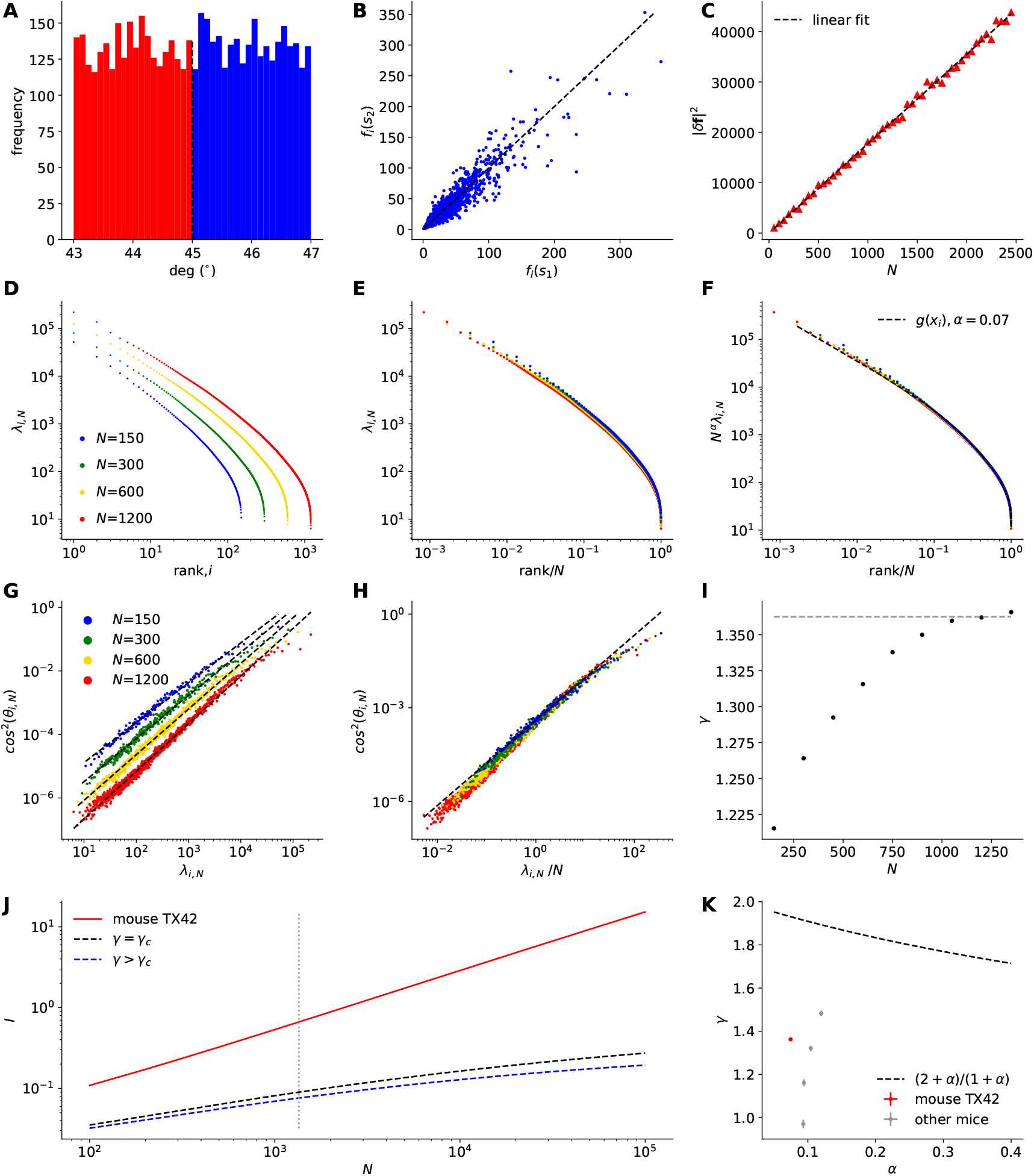
Bias-corrected analysis of mouse TX42 and the summary result from all mice. The figure styles follow Fig. 2 in the main text, except for **H** and **I**. The fitted scaling parameter is *α* = 0.074. The fitted slope is *γ* = 1.362 ± 0.002. **A**, A frequency histogram of grating stimulus orientations. **B**, Comparison of mean responses of neurons *f*_*i*_(*s*) to Stimuli 1 and 2. **C**, |*δ***f**_*N*_|^2^ vs the subpopulation size *N*. **D**, Noise eigenspectra for different *N*. **E**, The eigenspectra vs the normalized rank, (*x*_*i*_ =)*i*/*N*. The fitted scaling parameter is *α* = 0.074. The fitted parameters of *g*(*x*_*i*_) are *a* = 2.118, *b* = 2.078, *c* = 2.407, and *ν* = 0.213. **F**, The re-scaled eigenspectra by multiplying *N*^*α*^. The dashed line is the fitted *g*(*x*_*i*_). **G**, cos^2^ *θ*_*i,N*_ vs *λ*_*i,N*_ for different *N*. The dashed lines are linear fits. **H**, cos^2^ *θ*_*i,N*_ vs the scaled eigenvalue, *λ*_*i,N*_ /*N*. The dashed line is a linear fit. **I**, The estimated *γ* for different *N*. The average *γ* for *N* = 1000–1350 (dashed line) was 1.362 ± 0.002. **J**, The inferred LFI (solid line) as a function of *N* in the log-log plot. The dashed lines show theoretical LFI. The vertical dashed line is the number of neurons used to estimate the parameters. **K**, The estimated *γ* vs *α*. The dashed line is a critical boundary.

**Fig. S8.**
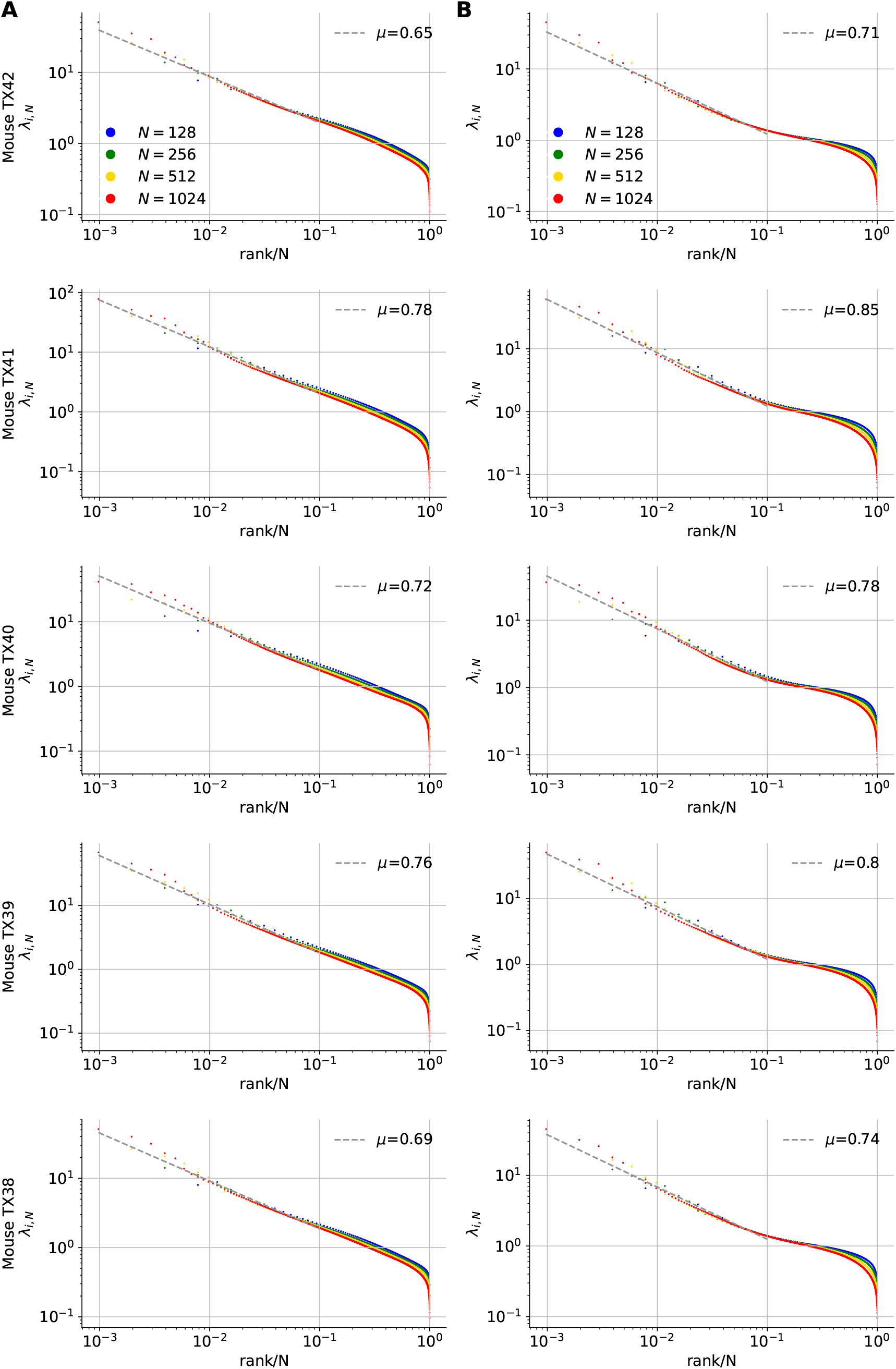
Eigenspectra of normalized covariance matrices for all mice. **A**, Eigenspectra of covariances of the activity normalized by nonzero mean for each neuron (Slope: *μ* = 0.72 ± 0.05 SD). The colors indicate *N* = 128, 256, 512, 1024. **B**, Eigenspectra of correlation coefficient matrices (Slope: *μ* = 0.78 ± 0.05 SD). The bias-correction methods were applied to this analysis.

**Fig. S9.**
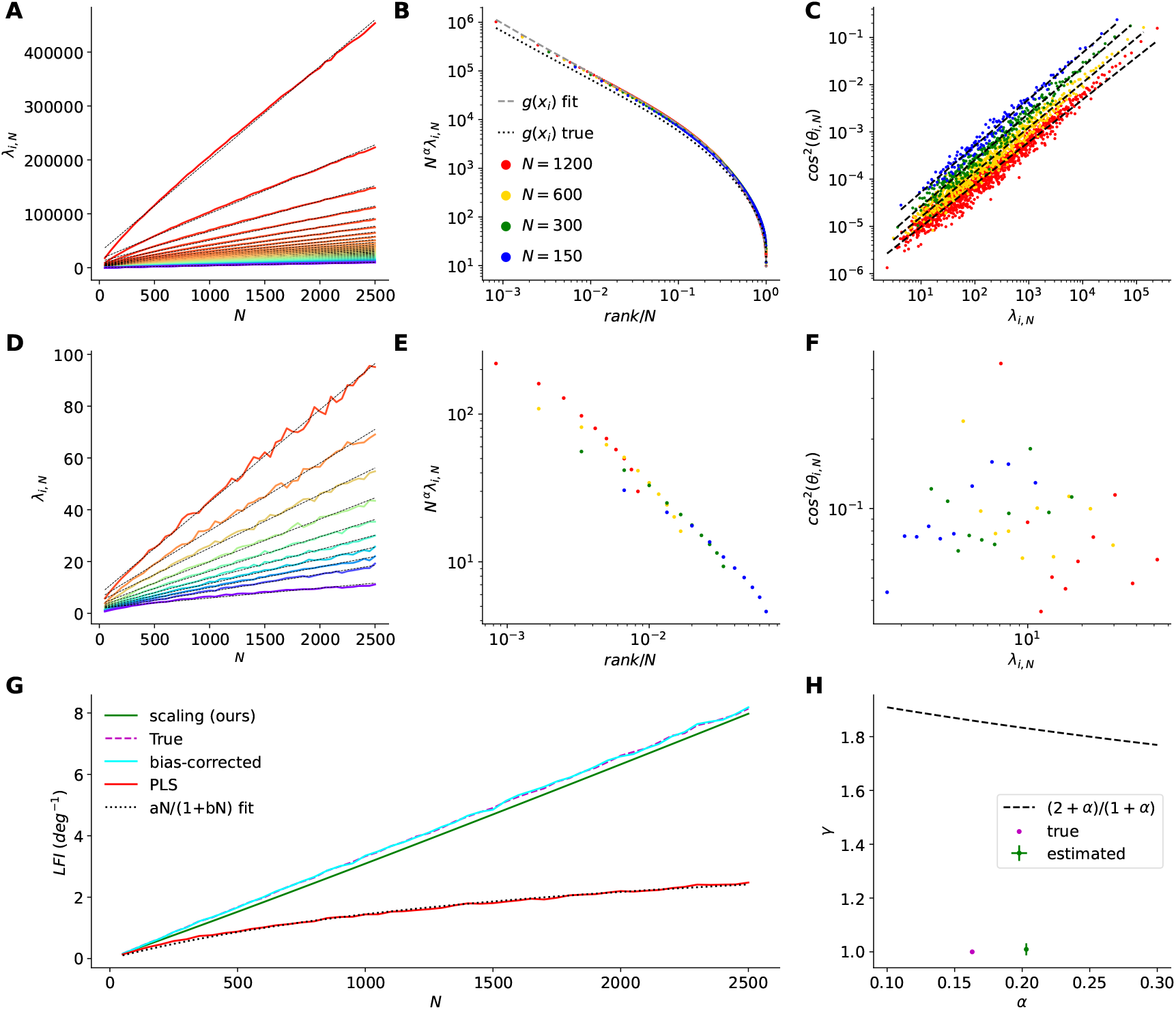
Comparison of the PLS analysis with the proposed scaling-based method using simulated data with nonzero *α*. The simulated data were generated from the eigenspectrum fitted to mouse TX42 (*α* = 0.163). Angular occupations were obtained using Eq. 7 with *γ* = 1. **A**, Eigenvalues as a function of population size *N*. Only the largest 50 eigenvalues are shown. The leading eigenvalues scale linearly for *α* > 0. Black lines, linear fits. **B**, Collapsed eigenspectrum of the simulated data (colored dots) with the true (dotted) and fitted (dashed) eigenspectra. The fitted scaling parameter is *α* = 0.203 ± 0.0001. **C**, Relation between the sampled eigenvalues and angular occupations. The estimated *γ* by the extrapolation method (see main text) is *γ* = 1.009 ± 0.011. **D**, Eigenvalues of the 10-dimensional subspace obtained by the PLS method as a function of population size *N*. Black lines, linear fits. **E**, Collapsed eigenspectrum of the subspaces obtained by the PLS method. **F**, Relation between the sampled eigenvalues and angular occupations in the subspaces obtained by the PLS method. **G**, Comparison of the LFI estimated by the scaling-based method (green) and PLS method (red) with the true LFI (dashed magenta). The PLS estimate underestimates the true LFI and is well fitted by the bounded function *aN*/(1 + *bN*) (dotted black), while the scaling-based method correctly estimates the true unbounded LFI. For comparison, the bias-corrected estimate of the LFI is shown (cyan). **H**, Estimated *γ*_∞_ vs *α* by the scaling-based method. The dotted line is a critical boundary (Eq. 8), below which the LFI is unbounded. Error bars, 2 SD.

**Fig. S10.**
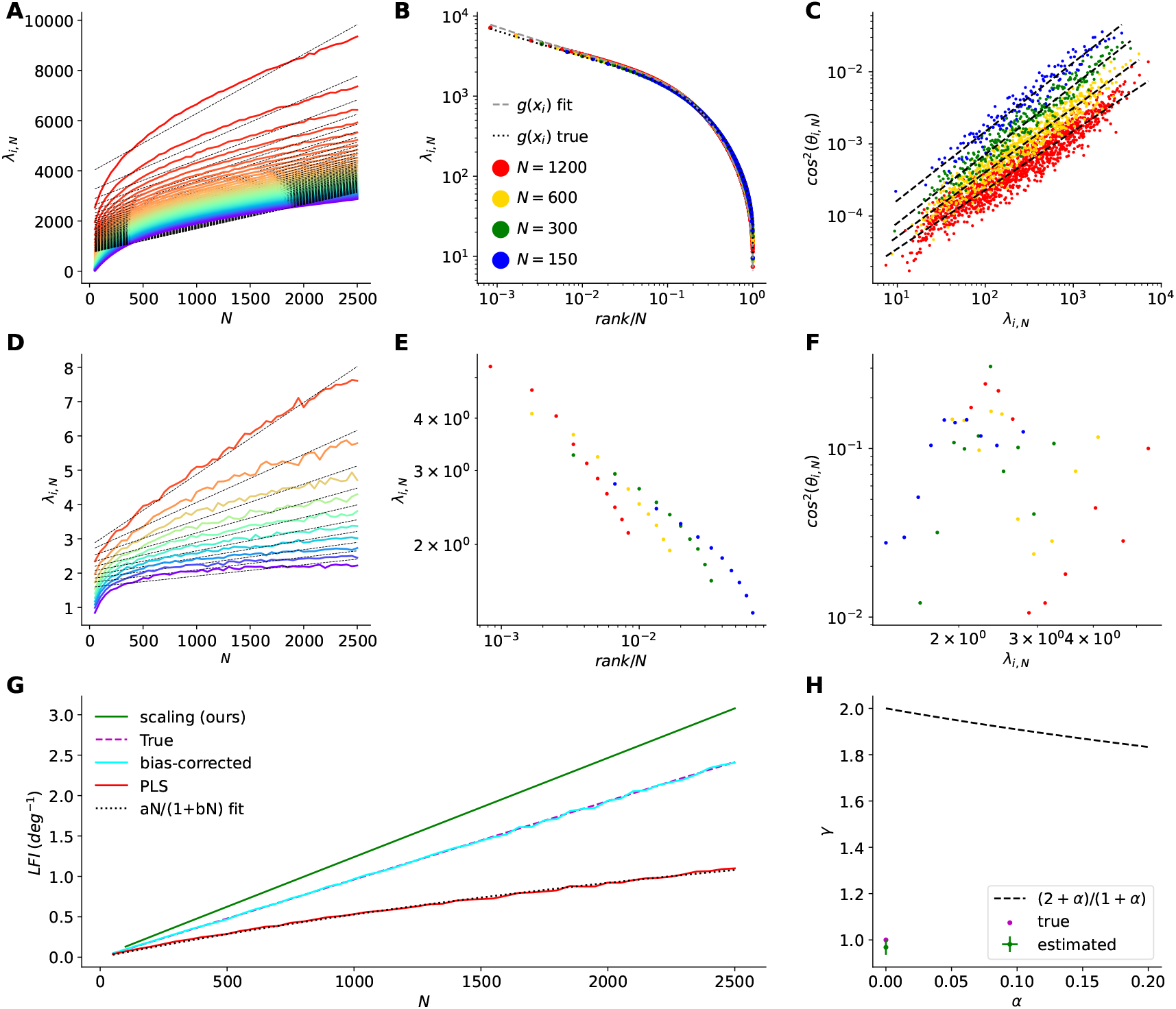
Comparison of the PLS analysis with the proposed scaling-based method using simulated data with *α* = 0. The analysis is the same as in fig. S9, but with simulated data generated using *α* = 0 and *g*(*x*), whose parameters were those fitted to mouse TX42 except for the power-law exponent, which was set to 0.5. Angular occupations were obtained using Eq. 7 with *γ* = 1. **A**, Eigenvalues as a function of population size *N*. Only the largest 50 eigenvalues are shown. All eigenvalues scale sublinearly for *α* = 0. Black lines, linear fits. **B**, Eigenspectrum of the simulated data (colored dots) with the true (dotted) and fitted (dashed) eigenspectra. The scaling-collapse fit is consistent with *α* = 0. **C**, Relation between the sampled eigenvalues and angular occupations. The estimated *γ* by the extrapolation method (see main text) is *γ* = 0.968 ± 0.016. **D**, Eigenvalues of the 10-dimensional subspace obtained by the PLS method as a function of population size *N*. One of the subspace eigenvalues scales linearly. Black lines, linear fits. **E**, Collapsed eigenspectrum of the subspaces obtained by the PLS method. **F**, Relation between the sampled eigenvalues and angular occupations in the subspaces obtained by the PLS method. **G**, Comparison of the LFI estimated by the scaling-based method (green) and PLS method (red) with the true LFI (dashed magenta). The PLS estimate underestimates the true LFI and is well fitted by the bounded function *aN*/(1 + *bN*) (dotted black), while the scaling-based method overestimates the true unbounded LFI. For comparison, the bias-corrected estimate of the LFI is shown (cyan). **H**, Estimated *γ*_∞_ vs *α* by the scaling-based method. The dotted line is a critical boundary (Eq. 8), below which the LFI is unbounded. Error bars, 2 SD.

## References

1 C. Stringer, M. Pachitariu, N. Steinmetz, C. B. Reddy, M. Carandini, K. D. Harris, Spontaneous behaviors drive multidimensional, brainwide activity. Science 364, eaav7893 (2019).

2 F. Lanore, N. A. Cayco-Gajic, H. Gurnani, D. Coyle, R. A. Silver, Cerebellar granule cell axons support high-dimensional representations. Nat. Neurosci. 24, 1142–1150 (2021).

3 J. Manley, S. Lu, K. Barber, J. Demas, H. Kim, D. Meyer, F. M. Traub, A. Vaziri, Simultaneous, cortex-wide dynamics of up to 1 million neurons reveal unbounded scaling of dimensionality with neuron number. Neuron 112, 1694–1709 (2024).

4 L. Meshulam, J. L. Gauthier, C. D. Brody, D. W. Tank, W. Bialek, Coarse graining, fixed points, and scaling in a large population of neurons. Phys. Rev. Lett. 123, 178103 (2019).

5 G. B. Morales, S. Di Santo, M. A. Muñoz, Quasiuniversal scaling in mouse-brain neuronal activity stems from edge-of-instability critical dynamics. Proc. Natl. Acad. Sci. U.S.A. 120, e2208998120 (2023).

6 Z. Wang, W. Mai, Y. Chai, K. Qi, H. Ren, C. Shen, S. Zhang, G. Tan, Y. Hu, Q. Wen, The geometry and dimensionality of brain-wide activity. Elife 14, RP100666 (2025).

7 G. Nicoletti, S. Suweis, A. Maritan, Scaling and criticality in a phenomenological renormalization group. Phys. Rev. Res 2, 023144 (2020).

8 Y. Hu, H. Sompolinsky, The spectrum of covariance matrices of randomly connected recurrent neuronal networks with linear dynamics. PLoS Comput. Biol. 18, e1010327 (2022).

9 K. H. Britten, M. N. Shadlen, W. T. Newsome, J. A. Movshon, The analysis of visual motion: a comparison of neuronal and psychophysical performance. J Neurosci. 12, 4745–4765 (1992).

10 E. Zohary, M. N. Shadlen, W. T. Newsome, Correlated neuronal discharge rate and its implications for psychophysical performance. Nature 370, 140–143 (1994).

11 B. B. Averbeck, P. E. Latham, A. Pouget, Neural correlations, population coding and computation. Nat. Rev. Neurosci. 7, 358–366 (2006).

12 M. N. Shadlen, W. T. Newsome, The variable discharge of cortical neurons: implications for connectivity, computation, and information coding. J Neurosci. 18, 3870–3896 (1998).

13 A. Kohn, R. Coen-Cagli, I. Kanitscheider, A. Pouget, Correlations and neuronal population information. Annu. Rev. Neurosci. 39, 237–256 (2016).

14 S. Panzeri, M. Moroni, H. Safaai, C. D. Harvey, The structures and functions of correlations in neural population codes. Nat. Rev. Neurosci. 23, 551–567 (2022).

15 R. Bartolo, R. C. Saunders, A. R. Mitz, B. B. Averbeck, Information-limiting correlations in large neural populations. J Neurosci. 40, 1668–1678 (2020).

16 O. I. Rumyantsev, J. A. Lecoq, O. Hernandez, Y. Zhang, J. Savall, R. Chrapkiewicz, J. Li, H. Zeng, S. Ganguli, M. J. Schnitzer, Fundamental bounds on the fidelity of sensory cortical coding. Nature 580, 100–105 (2020).

17 M. Kafashan, A. W. Jaffe, S. N. Chettih, R. Nogueira, I. Arandia-Romero, C. D. Harvey, R. Moreno-Bote, J. Drugowitsch, Scaling of sensory information in large neural populations shows signatures of information-limiting correlations. Nat. Commun. 12, 473 (2021).

18 O. Hazon, V. H. Minces, D. P. Tomàs, S. Ganguli, M. J. Schnitzer, P. E. Jercog, Noise correlations in neural ensemble activity limit the accuracy of hippocampal spatial representations. Nat. Commun. 13, 4276 (2022).

19 L. F. Abbott, P. Dayan, The effect of correlated variability on the accuracy of a population code. Neural Comput. 11, 91–101 (1999).

20 H. Sompolinsky, H. Yoon, K. Kang, M. Shamir, Population coding in neuronal systems with correlated noise. Phys. Rev. E 64, 051904 (2001).

21 C. Stringer, M. Michaelos, D. Tsyboulski, S. E. Lindo, M. Pachitariu, High-precision coding in visual cortex. Cell 184, 2767–2778 (2021).

22 R. Moreno-Bote, J. Beck, I. Kanitscheider, X. Pitkow, P. Latham, A. Pouget, Information-limiting correlations. Nat. Neurosci. 17, 1410–1417 (2014).

23 A. S. Ecker, P. Berens, A. S. Tolias, M. Bethge, The effect of noise correlations in populations of diversely tuned neurons. J Neurosci. 31, 14272–14283 (2011).

24 I. Kanitscheider, R. Coen-Cagli, A. Pouget, Origin of information-limiting noise correlations. Proc. Natl. Acad. Sci. U.S.A. 112, E6973–E6982 (2015).

25 I. Kanitscheider, R. Coen-Cagli, A. Kohn, A. Pouget, Measuring Fisher information accurately in correlated neural populations. PLoS Comput. Biol. 11, e1004218 (2015).

26 M. L. Leavitt, F. Pieper, A. J. Sachs, J. C. Martinez-Trujillo, Correlated variability modifies working memory fidelity in primate prefrontal neuronal ensembles. Proc. Natl. Acad. Sci. U.S.A. 114, E2494–E2503 (2017).

27 H. Safaai, M. von Heimendahl, J. M. Sorando, M. E. Diamond, M. Maravall, Coordinated population activity underlying texture discrimination in rat barrel cortex. J Neurosci. 33, 5843–5855 (2013).

28 M. N. Insanally, I. Carcea, R. E. Field, C. C. Rodgers, B. DePasquale, K. Rajan, M. R. DeWeese, B. F. Albanna, R. C. Froemke, Spike-timing-dependent ensemble encoding by non-classically responsive cortical neurons. Elife 8, e42409 (2019).

29 J. Zylberberg, The role of untuned neurons in sensory information coding. bioRxiv p. 134379 (2017).

30 Y. Osako, T. Ohnuki, Y. Tanisumi, K. Shiotani, H. Manabe, Y. Sakurai, J. Hirokawa, Contribution of non-sensory neurons in visual cortical areas to visually guided decisions in the rat. Curr. Biol. 31, 2757–2769 (2021).

31 J. S. Montijn, R. G. Liu, A. Aschner, A. Kohn, P. E. Latham, A. Pouget, Strong information-limiting correlations in early visual areas. bioRxiv p. 842724 (2019).

32 W.-H. Zhang, S. Wu, K. Josić, B. Doiron, Sampling-based Bayesian inference in recurrent circuits of stochastic spiking neurons. Nat. Commun. 14, 7074 (2023).

33 S. Camargo, D. A. Martin, E. J. A. Trejo, A. de Florian, M. A. Nowak, S. A. Cannas, T. S. Grigera, D. R. Chialvo, Scale-free correlations in the dynamics of a small (N 10000) cortical network. Phys. Rev. E 108, 034302 (2023).

34 A. Arieli, A. Sterkin, A. Grinvald, A. Aertsen, Dynamics of ongoing activity: explanation of the large variability in evoked cortical responses. Science 273, 1868–1871 (1996).

35 M. Tsodyks, T. Kenet, A. Grinvald, A. Arieli, Linking spontaneous activity of single cortical neurons and the underlying functional architecture. Science 286, 1943–1946 (1999).

36 T. Kenet, D. Bibitchkov, M. Tsodyks, A. Grinvald, A. Arieli, Spontaneously emerging cortical representations of visual attributes. Nature 425, 954–956 (2003).

37 A. Luczak, P. Barthó, K. D. Harris, Spontaneous events outline the realm of possible sensory responses in neocortical populations. Neuron 62, 413–425 (2009).

38 P. Berkes, G. Orbán, M. Lengyel, J. Fiser, Spontaneous cortical activity reveals hallmarks of an optimal internal model of the environment. Science 331, 83–87 (2011).

39 T. Matsui, T. Hashimoto, T. Murakami, M. Uemura, K. Kikuta, T. Kato, K. Ohki, Orthogonalization of spontaneous and stimulus-driven activity by hierarchical neocortical areal network in primates. Nature Communications 15, 1–13 (2024).

40 V. Minces, L. Pinto, Y. Dan, A. A. Chiba, Cholinergic shaping of neural correlations. Proc. Natl. Acad. Sci. U.S.A. 114, 5725–5730 (2017).

41 M. C. Dadarlat, M. P. Stryker, Locomotion enhances neural encoding of visual stimuli in mouse V1. J Neurosci. 37, 3764–3775 (2017).

42 M. R. Cohen, J. H. Maunsell, Attention improves performance primarily by reducing interneuronal correlations. Nat. Neurosci. 12, 1594–1600 (2009).

43 O. Ledoit, M. Wolf, A well-conditioned estimator for large-dimensional covariance matrices. Journal of multivariate analysis 88, 365–411 (2004).

44 M. Pachitariu, M. Michaelos, C. Stringer, Recordings of 20,000 neurons from V1 in response to oriented stimuli. figshare.org (2019), doi:10.25378/janelia.8279387, https://doi.org/10.25378/janelia.8279387.

